# Thalamic control of sensory enhancement and sleep spindle properties in a biophysical model of thalamoreticular microcircuitry

**DOI:** 10.1101/2022.02.28.482273

**Authors:** Elisabetta Iavarone, Jane Simko, Ying Shi, Marine Bertschy, María García-Amado, Polina Litvak, Anna-Kristin Kaufmann, Christian O’Reilly, Oren Amsalem, Marwan Abdellah, Grigori Chevtchenko, Benoît Coste, Jean-Denis Courcol, András Ecker, Cyrille Favreau, Adrien Christian Fleury, Werner Van Geit, Michael Gevaert, Nadir Román Guerrero, Joni Herttuainen, Genrich Ivaska, Samuel Kerrien, James G. King, Pramod Kumbhar, Patrycja Lurie, Ioannis Magkanaris, Vignayanandam Ravindernath Muddapu, Jayakrishnan Nair, Fernando L. Pereira, Rodrigo Perin, Fabien Petitjean, Rajnish Ranjan, Michael Reimann, Liviu Soltuzu, Mohameth François Sy, M. Anıl Tuncel, Alexander Ulbrich, Matthias Wolf, Francisco Clascá, Henry Markram, Sean L. Hill

## Abstract

Thalamoreticular circuitry is known to play a key role in attention, cognition and the generation of sleep spindles, and is implicated in numerous brain disorders, but the cellular and synaptic mechanisms remain intractable. Therefore, we developed the first detailed computational model of mouse thalamus and thalamic reticular nucleus microcircuitry that captures morphological and biophysical properties of ∼14,000 neurons connected via ∼6M synapses, and recreates biological synaptic and gap junction connectivity. Simulations recapitulate multiple independent network-level experimental findings across different brain states, providing a novel unifying cellular and synaptic account of spontaneous and evoked activity in both wakefulness and sleep. Furthermore, we found that: 1.) inhibitory rebound produces frequency-selective enhancement of thalamic responses during wakefulness, in addition to its role in spindle generation; 2.) thalamic interactions generate the characteristic waxing and waning of spindle oscillations; and 3.) changes in thalamic excitability (e.g. due to neuromodulation) control spindle frequency and occurrence. The model is openly available and provides a new tool to interpret spindle oscillations and test hypotheses of thalamoreticular circuit function and dysfunction across different network states in health and disease.

## 1 Introduction

The thalamus and thalamic reticular nucleus lie at the heart of the thalamocortical system in the mammalian brain and are tightly integrated with the neocortex via extensive reciprocal connections (Jones, 2007). Thalamic relay cells project to the cortex and form excitatory connections with thalamic reticular neurons, which in return send inhibitory projections to the thalamus, forming the thalamoreticular circuit (Ohara and Lieberman, 1985; Pinault, 2004; Scheibel and Scheibel, 1966). Thalamoreticular circuitry plays a key role in numerous functions, such as the transmission of sensory information to the cortex (Steriade et al., 1997) and the transition between brain states, including sleep and wakefulness (Jones, 2002; Rikhye et al., 2018; Steriade, 2003). Thalamoreticular circuitry has been implicated in attentional processes (McAlonan et al., 2008; Wimmer et al., 2015), the generation of the alpha rhythm (Hughes and Crunelli, 2005; Nestvogel and McCormick, 2021; Saalmann et al., 2012), and the generation of spindle oscillations during sleep (T. Bal et al., 1995; Contreras et al., 1997; Fernandez and Luthi, 2019; Steriade et al., 1987; von Krosigk et al., 1993). Alterations in thalamic neuron firing and their interconnectivity have been associated with pathological brain rhythms, such as those appearing in absence epilepsy (Beenhakker and Huguenard, 2009; Huguenard and McCormick, 2007; Makinson et al., 2017; Sohal and Huguenard, 2003; Steriade, 2005). Changes in the incidence and density of spindle oscillations during sleep have been observed in different disorders, such as schizophrenia (Castelnovo et al., 2018; Ferrarelli et al., 2010, 2007; Manoach et al., 2016, 2014), neurodevelopmental disorders (Gruber and Wise, 2016), attention deficit hyperactivity disorder (Saito et al., 2019), Alzheimer’s disease (Weng et al., 2020), among others.

Although the properties of thalamic and reticular neurons have been extensively studied *in vitro* (Connelly et al., 2017; Cox et al., 1996; Jahnsen and Llinás, 1984; Lee et al., 2007; Pinault et al., 1995; Pinault and Deschênes, 1998; Spreafico et al., 1991), there are still significant gaps in our understanding of thalamoreticular circuitry (O’Reilly et al., 2021). Computer models and simulations can facilitate the integration and standardization of different sources of experimental data, highlight key missing experiments, while providing a tool to test hypotheses and explore the structural and functional complexity of neural circuits (Billeh et al., 2020; Einevoll et al., 2019; Markram et al., 2015a). Previous models of small thalamic networks or thalamic slices have focused on investigating specific physiological and pathological aspects of thalamic microcircuits, wth the level of detail and choice of biophysical mechanisms dictated by their hypotheses (Bazhenov et al., 1998; Bús et al., 2018; Destexhe et al., 1996; Golomb et al., 1996; Li et al., 2017; Wang et al., 1995).

In this work, we follow and extend the reconstruction pipeline presented in (Markram et al., 2015a) to develop a digital model of a thalamoreticular microcircuit of a portion of first-order somatosensory thalamus in the adult mouse, including the ventral posterolateral nucleus (VPL) and the corresponding region of the reticular nucleus (Rt). We performed targeted *in vitro* experiments to collect electrophysiological, morphological and synaptic data from the mouse. We then used these measurements, combined with systematic curation of data from the literature and open access datasets, to build biophysically and morphologically-detailed neuron models. We defined the microcircuit geometry and populated it with experimentally-measured neuron densities. Three-dimensional morphological reconstructions constituted the basis to constrain the detailed connectivity between neurons of the thalamus and the reticular nucleus. Synaptic connections comprised chemical synapses with short-term depression and facilitation and electrical synapses (gap junctions). Extra-thalamic inputs, were modeled by including synapses from sensory afferents (medial lemniscus) and corticothalamic feedback.

This approach yielded the first morphologically and biophysically-detailed model of a thalamic microcircuit, demonstrating that the modeling strategy developed for cortical microcircuitry (Markram et al., 2015a) can be applied to other brain regions. Although this model is constrained with only cellular and synaptic experimental data, we found that it reproduces a number of network-level *in vitro* and *in vivo* findings. Where data was missing, we used generalization principles and tested whether the model reproduces experimental findings that were not used during building (validation process). After validating the model at different levels, we studied its dynamics in wakefulness and sleep-like states and rhythm generation (spindle-like oscillations). Supercomputer-based simulations of the model recapitulate multiple independent network-level experimental findings in wakefulness and sleep including spontaneous and evoked activity, stimulus dependent recruitment of thalamic inhibition, thalamic surround inhibition, adaptation of thalamic sensory responses, reticular nucleus-triggered spindle oscillations, and other properties of spindle oscillations during sleep, including cellular and synaptic mechanisms that have been previously implicated. Beyond recapitulating known properties and mechanisms, we found that the inhibitory rebound of thalamic relay cells results in frequency-selective enhancement of thalamic responses during wakefulness, in addition to its role in spindle generation during sleep. The characteristic waxing and waning of spindle oscillations was found to be generated intrathalamically and is not dependent on cortical factors. In addition, differential changes in thalamic and reticular cell excitability resulted in altered spindle frequencies and determined the incidence of spindle occurrence. This last point is particularly relevant to interpreting the presence or absence of spindles in different brain disorders. We provide the experimental data and computational models as a free resource for further hypothesis exploration and model development by the community.

## 2 Methods

### 2.1 Constraining and validating the model with experimental data

The model microcircuit was built by constraining and validating it at multiple levels, using experimental data and algorithmic approaches, based on methods published previously (Markram et al., 2015a). For validation we performed direct comparison of the model properties with experimental measurements that were not used during the model building steps. Before describing the details of the reconstruction, validation and simulations, we provide a list of data used for constraining the model, the validation data and further validations at the network level.

#### 2.1.1 Experimental data used to constrain the model

The following experimental data was used to constrain the model, further details on the experimental procedures and literature references are provided below.

- Three-dimensional reconstructions of neuron morphologies, from *in vitro* and *in vivo* labeling
- Ion channel kinetic parameters
- Electrophysiological data from *in vitro* patch-clamp recordings (current step stimuli)
- Neuron densities
- Fraction of inhibitory and excitatory neurons
- Fraction of electrical types for each morphological type
- Axonal bouton densities (i.e. number of boutons per axonal unit length)
- Volumetric densities of lemniscal boutons (number of boutons per unit volume)
- Ratio of corticothalamic to lemniscal bouton densities and ratio of corticothalamic to thalamocortical bouton densities (volumetric data from the literature)
- Postsynaptic potential amplitudes and their change in response to trains of presynaptic inputs from *in vitro* paired-recordings (short-term plasticity protocols)
- Number of neurons connected through gap junctions
- Synaptic current kinetic parameters

#### 2.1.2 Experimental data used for model validation

The following experimental measurements were not used for constraining the model during the building process but were used for validation:

Electrophysiological data from *in vitro* patch-clamp recordings (current ramps and noise)
Number of synapses per connection between interneurons and thalamocortical neurons (*i.e.* number of synapses between each pair of neurons)
Synaptic convergence onto reticular neurons
Postsynaptic potential amplitudes (different subset of neuron pairs than the ones used to constrain the model)
Coefficient of variation of first postsynaptic potential amplitudes
Distance-dependent gap junction connectivity between reticular neurons
Gap junctions coupling coefficients

#### 2.1.3 Validations at the network level

We identified the following network responses during simulated activity as a general validation of the reconstruction process:

Spontaneous *in vivo*-like activity, characterized by uncorrelated firing and low firing rates in TC and Rt cells (Born et al., 2021; Hartings et al., 2000; Nestvogel and McCormick, 2021).
Evoked activity with simulated sensory input in TC as well Rt cells (Hartings et al., 2000).
Adaptation to repeated sensory stimuli at different frequencies (Manuel A Castro-Alamancos, 2002)
Corticothalamic inputs counterbalance sensory adaptation (Mease et al., 2014)
Increased thalamic bursts after brief stimulation of the reticular nucleus and evoked spindle-like oscillations (Halassa et al., 2011)
Evoked spindle-like oscillations (Halassa et al., 2011)
Initiation of spindle-like oscillations during cortical UP-states (Destexhe et al., 2007; McCormick and Bal, 1997; Steriade et al., 1993)

### 2.2 Reconstructing the morphological diversity of neurons

#### 2.2.1 Reconstruction of morphologies

A subset of 3D reconstructions of biocytin-stained thalamocortical (TC) neurons, reticular thalamic (Rt) neurons and thalamic interneurons (IN) were obtained from *in vitro* patch-clamp experiments from 300 μm slices of P14-35 mice (GAD67-eGFP or C57Bl/6J strains) as previously described (Iavarone et al., 2019; Markram et al., 2015). During the electrophysiological recordings neurons were stained intracellularly with biocytin. *In vitro*-stained neurons were mainly located in primary somatosensory nuclei (VPL, and ventral posteromedial nucleus - VPM) and the somatosensory sector of the reticular nucleus (Clemente-Perez et al., 2017; Lam and Sherman, 2011; Pinault and Deschênes, 1998). Reconstructions used the Neurolucida system (MicroBrightField) and were corrected for shrinkage along the thickness of the slice. Shrinkage along other dimensions was taken into account during the unraveling step (see below). Dendrites were reconstructed with a 100x magnification (oil immersion objective) and axons at 60x (water immersion objective).

*In vivo*-stained TC and Rt morphologies were obtained through different experimental techniques. In some cases, neurons were labeled by injection of replication-defective Sindbis virus particles in the thalamus or Rt nucleus in C57Bl/6J adult mice (Furuta et al., 2001) or electroporation of RNA of the same virus (Porrero et al., 2016). The virus labeled the membrane of the neurons thanks to a palmitoylation signal linked to a green fluorescent protein (GFP). Brains were cut in 50 μm serial sections and immunostained against GFP and enhanced with glucose oxidase-nickel staining (Shu et al., 1988). Neurons were reconstructed from sequentially-ordered slices under bright-field optics using the Neurolucida system (MicroBrightField). The complete method is described elsewhere (Rodriguez-Moreno et al., 2020).

*In vivo*-labelled TC morphologies were obtained from the Janelia Mouselight project, from sparsely-labeled adult C57/BL6 mice brains; the method is described in detail elsewhere (Winnubst et al., 2019) and summarized here. Brains were then delipidated, fluorescence was enhanced by immunolabeling and imaged with a 40x oil-immersion objective. This procedure generated large datasets of high-resolution image stacks. The 3D reconstructions were conducted combining semi-automated segmentation of the neurites, human annotation and quality control. Janelia Mouselight reconstructions lacked diameter variations in their neurites, which is important for accurate electrical modeling of neurons (Jaeger, 2000). For this reason, we only used their axons in order to increase the variability of our axonal reconstructions. We obtained 96 morphologies whose soma was located in the thalamus and we visually inspected their shape along with 3D meshes of the reticular nucleus of the thalamus using the Janelia MouseLight Project (RRID:SCR_016668; http://ml-neuronbrowser.janelia.org/). Since most thalamocortical neurons project to the Rt on their path to the cortex (Clascá et al., 2012; Lam and Sherman, 2011) we selected the 41 morphologies which gave off collaterals in the reticular nucleus. We assumed that neurons without collaterals in the Rt were partially labeled and/or reconstructed, since those collaterals are often very thin (Harris, 1987). Given the limited number of reconstructed morphologies of neurons in VPL and VPM in the Janelia MouseLight dataset, we included 27 axons (with collaterals to Rt) from other thalamic nuclei. To ensure that the connectivity would not be impacted, we analyzed the geometrical properties of the Rt collaterals and found that the difference within the same nucleus was as high as the difference between nuclei.

For *in vivo* labeling of reticular neurons virus injections for sparse labeling of whole brain neuron morphologies were employed in SSt-Cre;Ai139 adult mice (Daigle et al., 2018; Harris et al., 2014). Brains were imaged using fluorescence micro-optical sectioning tomography (fMOST) (Gong et al., 2016). Neurons were manually reconstructed from high resolution image stacks obtained after slicing. Further details of the method are available in related publications (Peng et al., 2021).

#### 2.2.2 Morphology analysis, alignment and visualization

Raw morphological data did not have a common orientation along a principal axis, which is necessary to place them in the microcircuit volume according to biologically-plausible constraints (see below). We thus computed a rotation matrix so that the principal axis of the morphology was parallel to the vertical axis of the microcircuit. The principal axis of TC morphologies was the one connecting the center of the soma and the center of mass of the axon collaterals in the Rt nucleus (see below). For Rt neurons, the principal axis connected the soma and the center of mass of the axonal arborization in the thalamus. After rotating the morphologies, we visually validated the results. Rotation of the INs was not performed, since no orientation information relative to known landmarks was available.

For morphology analysis we used the open-source library NeuroM (https://github.com/BlueBrain/NeuroM). To identify the TC axon collaterals projecting to the Rt we selected the morphological sections which had branch order >= 1 and path distance from the soma < 2,500 μm and visually validated the results. For some morphologies, we selected those having path distance <= 2,000 μm, because some TC neurons have collaterals projecting to other subcortical regions (e.g striatum), see (Clascá et al., 2012).

Raw morphological data were algorithmically corrected for slicing artifacts and processed to generate a large pool of unique morphologies for building the microcircuit and connectivity. Spurious sections, which were accidentally introduced during manual reconstruction, were identified as those having 0 μm diameter and removed. The details are described in Supplemental Experimental Procedure of the neocortical microcircuit model (Markram et al., 2015), and summarized below.

The morphology images in Fig. 4 were created using NeuroMorphoVis (Abdellah et al., 2018).

**Figure 1.**
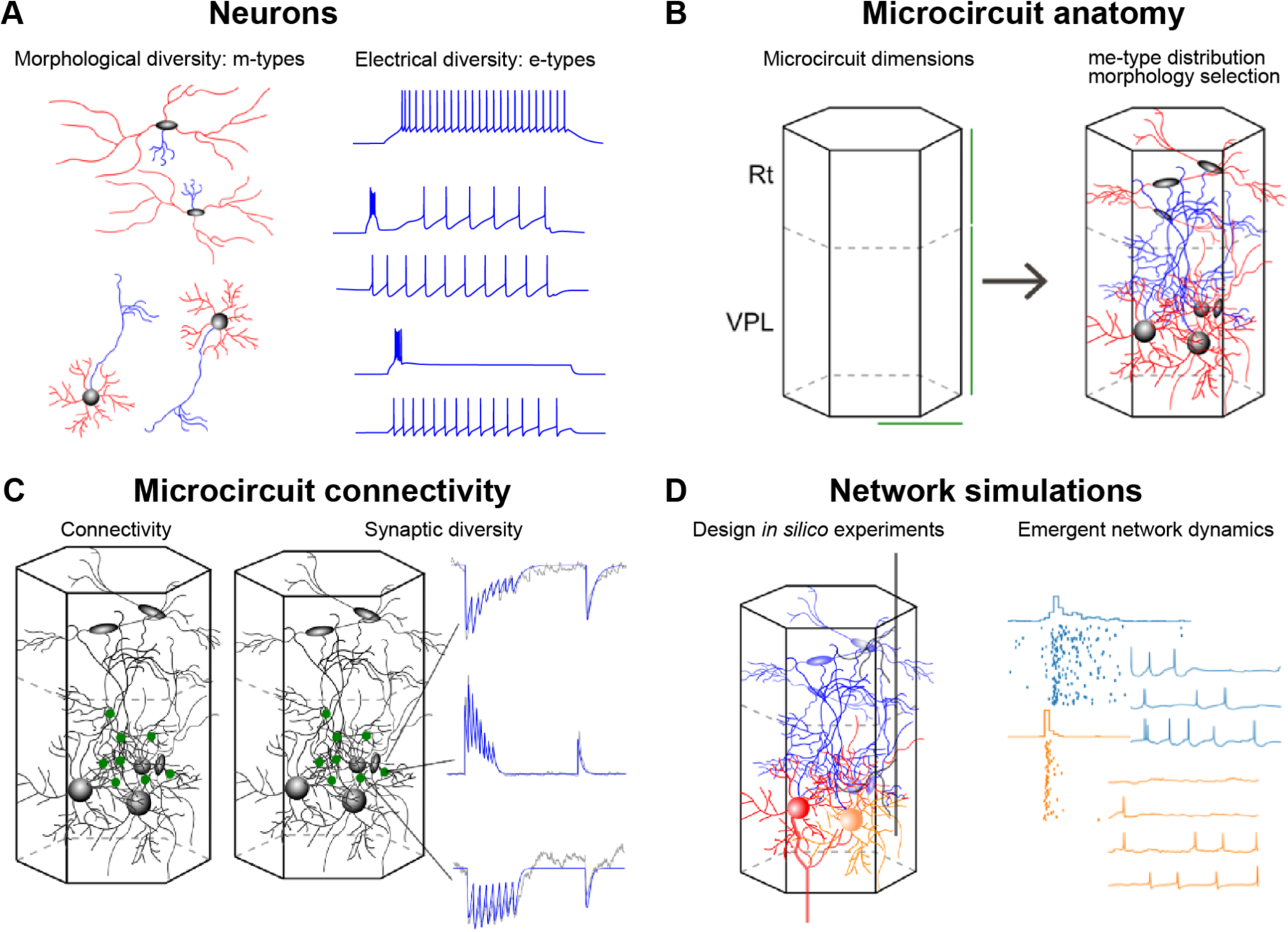
Workflow for the digital reconstruction of thalamoreticular microcircuitry. (**A**) Morphological and electrical diversity of thalamic neurons: identify the different morphological types (m-types) and electrical types (e-types) and build a large number of unique morphologically-detailed neuron models (me-models). (**B**) Define the microcircuit volume, with its horizontal and vertical dimensions. Place the me-models selecting the appropriate exemplar that best fits the anatomical constraints and axonal and dendritic distribution. (**C**) Build the connectivity starting from the morphological appositions and pruning them to match experimental constraints. Map the synaptic diversity of neurons based on the synaptic physiology of characterized pathways. (**D**) Introduce synapses formed by lemniscal and corticothalamic afferents for reproducing experiments *in silico* and studying emergent network dynamics.

**Figure 2.**
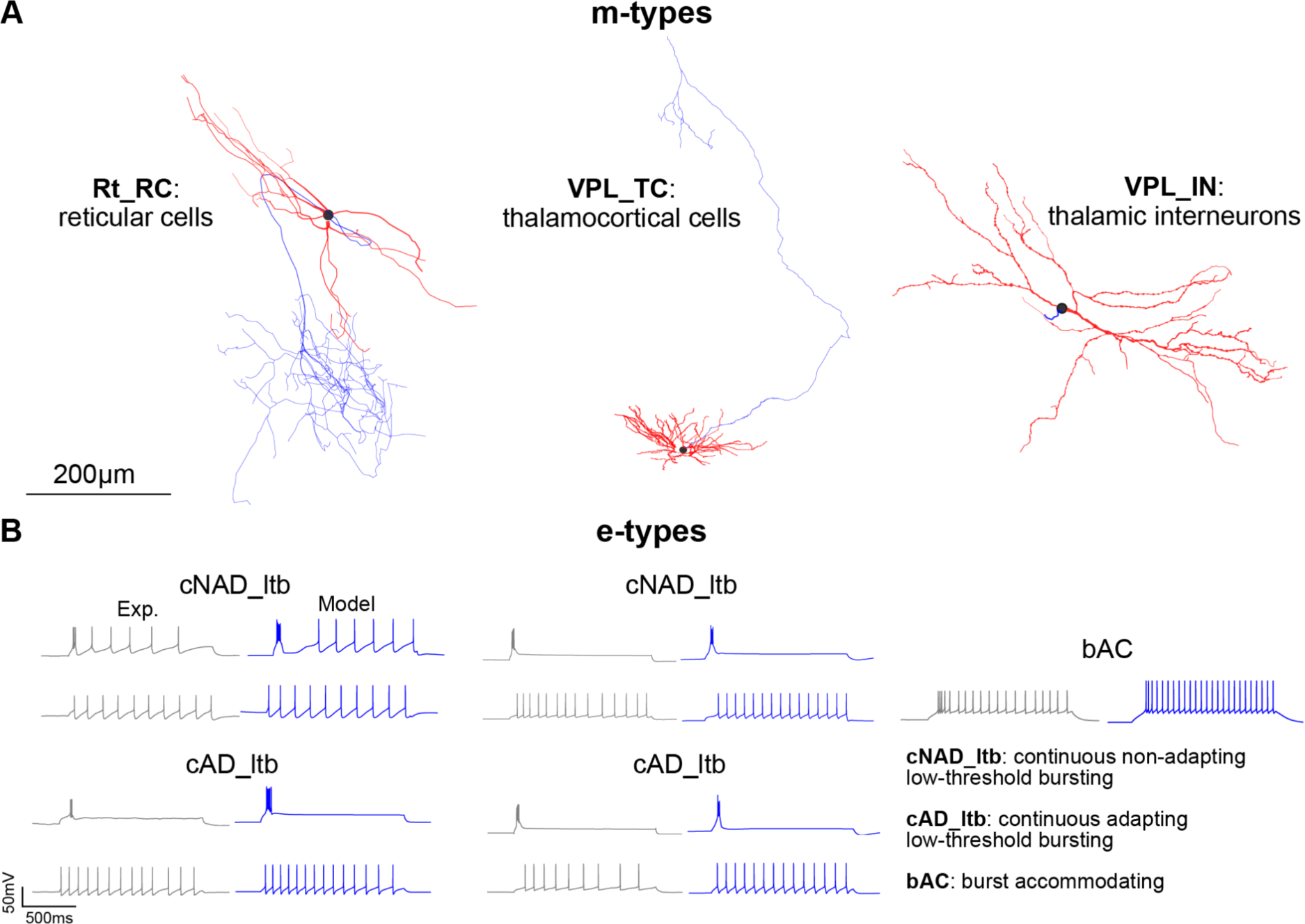
Single cell morphological and electrophysiological data and models. (**A**) Exemplar 3D reconstructions of three thalamic and reticular morphological types (m-types) from the mouse. Axon in blue, dendrites in red, soma in black. For the VPL_TC m-type, the axonal projection to the neocortex is not shown. All reconstructions are shown on the same scale. (**B**) Electrical types (e-types) and corresponding electrical models. From left to right: exemplar recordings (gray) and models (blue) corresponding to Rt_RC, VPL_TC and VPL_IN m-types. For cNAD_ltb and cAD_ltb e-types two distinct firing modes are shown: low threshold-bursting (first row) and tonic firing (second row).

**Figure 3.**
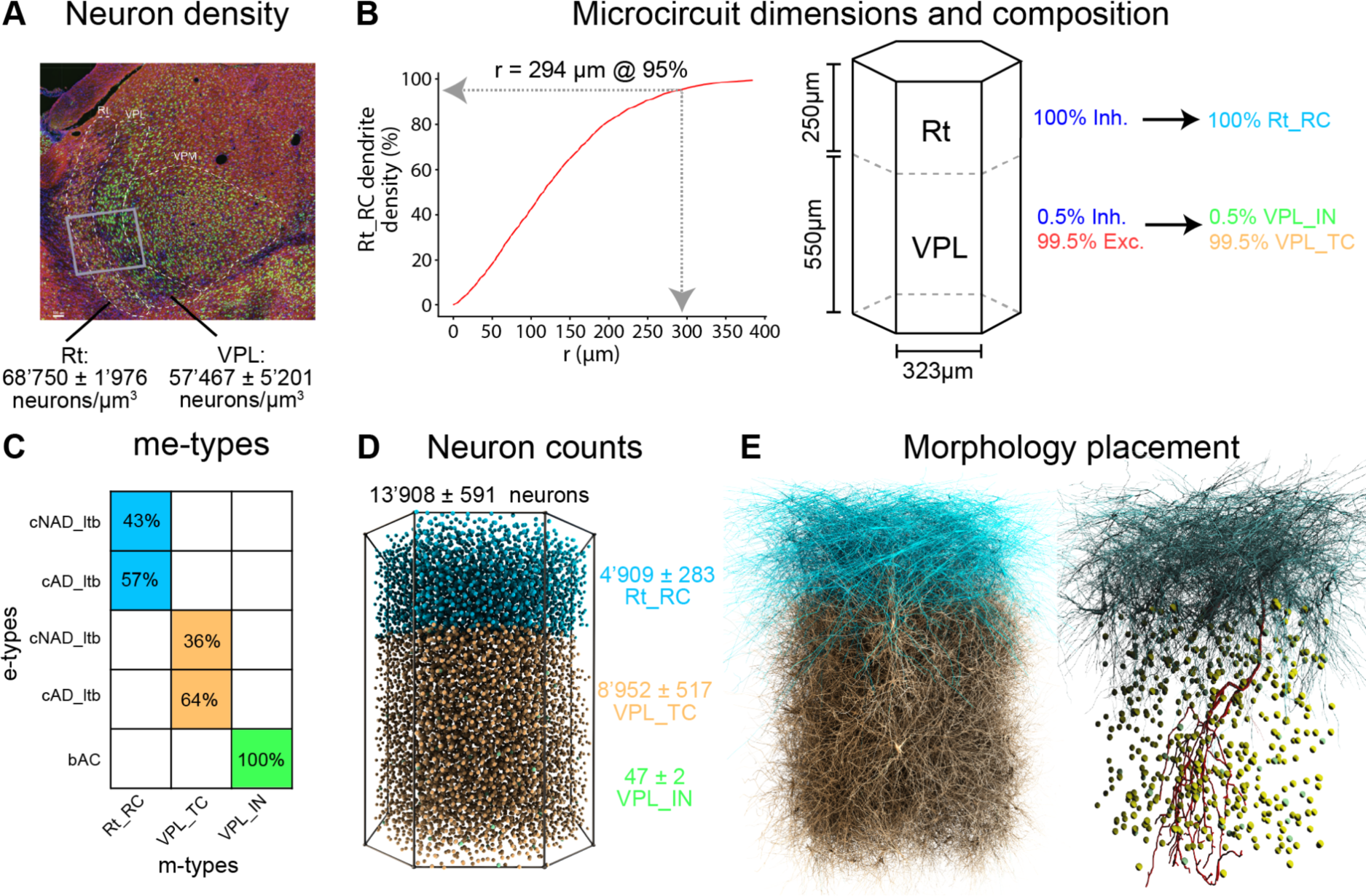
Reconstructing neuron densities, microcircuit dimensions and composition, neuron counts and morphology placement. (**A**) Mean neuron densities in the Rt and VPL nucleus of the thalamus. Confocal imaging of an exemplar slice after staining with anti-GABA (red), anti-NeuN (green) and DAPI (blue). Brain regions outlines were drawn after the alignment of the slice with the Allen Reference Atlas. The gray box represents a thalamic microcircuit. (**B**) Microcircuit dimensions (lateral and vertical dimensions). Left: the lateral dimension was the smallest circle for obtaining saturated Rt_RC dendritic density at the center of the microcircuit. The cut-off radius at 95% of the plateau density was 294 μm. Middle: hexagonal boundaries were adopted for tiling and vertical dimensions of the Rt and VPL regions were calculated from the Allen Reference Atlas (see Methods). Right: excitatory/inhibitory fractions and m-types composition. Inhibitory fractions as reported in the Mouse Cell Atlas (Erö et al., 2018). (**C**) Fraction of e-types corresponding to each m-type as found in our single cell recordings. (**D**) Predicted neuron numbers and soma positions in the microcircuits (mean and std of the five microcircuit instances). (**E**) Morphologies placed in the microcircuit, only ∼10% of the neurons are shown (left) and axons are omitted for clarity. Right: one exemplar Rt_RC axon (red) is shown innervating the VPL.

**Figure 4.**
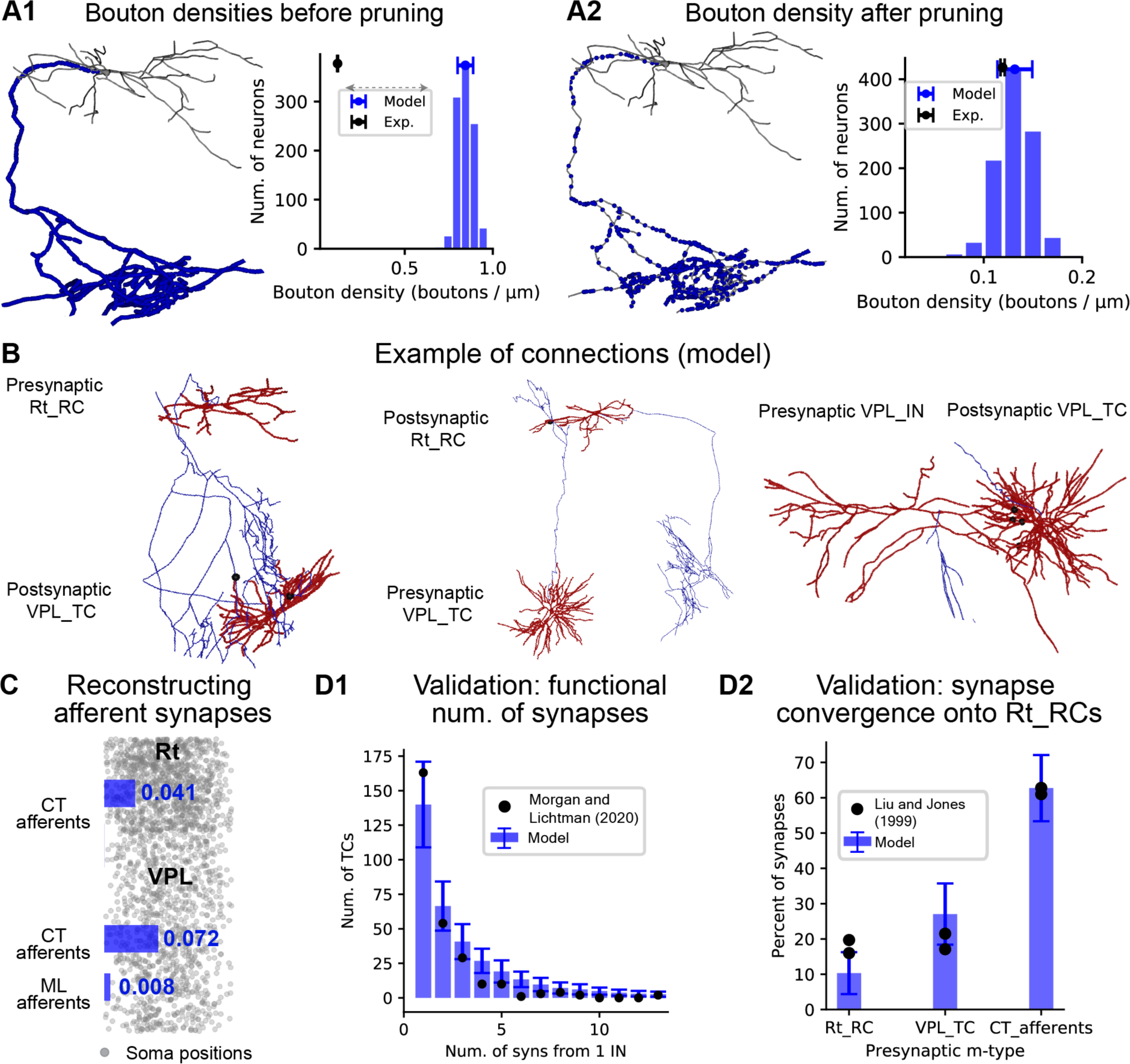
Reconstructing and validating intrathalamic and thalamic afferent connectivity. (**A1-2**) Constraining intrathalamic connectivity using neuron morphologies and bouton densities. (**A1**) As a first step, axodendritic appositions are used as location of putative synapses. The connectivity based on these appositions is characterized by high numbers of bouton densities (number of boutons / axonal length). Left: the location of putative synapses is shown for an exemplar Rt_RC neuron (red: dendrites, blue: axon, black: soma). Right: distribution of bouton density for 1,000 Rt_RC morphologies in the model and mean and std in the experiment (n=2 Rt_RC morphologies). (**A2**) The experimental bouton densities are used as a constraint to remove a fraction of axodendritic appositions (see Methods for details). (**B**) Example of the resulting mono- and multi-synapse connections between pairs of neurons in the model. Black dots represent the location of functional synapses. (**C**) Volumetric bouton densities (numbers reported in boutons/μm^3^) was used as a constraint to add synapses from medial lemniscal (ML) and corticothalamic (CT) afferents. (**D1**) Validation of the distribution of synapses per connection. The model was compared to findings from an EM reconstruction of 1 interneuron in the mouse (Morgan & Lichtman, 2020), 47 VPL_IN in the model (columns are the mean and bars the std). (**D2**) Validation of synapse convergence onto Rt_RC neurons in the model was validated against electron microscope (EM) experiments in the rat (N=2, Liu & Jones (1999), black dots). Bars and vertical lines show mean and standard deviation for all Rt_RCs in the model (N=4,909).

#### 2.2.3 Unraveling morphologies

Since we found that 3D reconstructions from *in vitro*-stained neurons had increased tortuosity in their dendrites as a result of tissue shrinkage, we unraveled them using an existing algorithm (Markram et al., 2015). This process resulted in an increase of the reach of the morphologies, while preserving the original length of the branches. Briefly, unraveling was performed by sections and for each section a sliding window composed of a given number of successive points was created. The number of points in the sliding window (N) was the only parameter of the algorithm and we found that N=5 previously used performed well on thalamic morphologies. The general direction of the points in the window was computed using principal component analysis (PCA). The segment at the middle of the window was then aligned along this direction. It meant that its direction was set to the one of the sliding windows but it retained its original length. The sliding window was moved over all points of the section and the algorithm was applied to all sections.

#### 2.2.4 Repairing morphologies

Most of the *in vitro*-stained morphologies were truncated at slice edges and in the case of some TC morphologies, which have very dense dendritic arborization, this resulted in a significant decrease in dendritic mass. We applied an existing algorithm (Anwar et al., 2009; Markram et al., 2015) to repair missing dendritic branches. First, the algorithm detects cut points on the XY plane, i.e. the plane parallel to the slice, along the Z direction (parallel to the slice thickness). The 3D coordinate system was centered on the morphology soma. Although the algorithm was designed to detect cut points on two planes, we found that our morphologies were truncated on the top plane. We improved the algorithm by searching the cut points before unraveling the morphologies and updated their position during the unraveling step. Cut detection required a tolerance parameter to detect terminal points within a certain distance from maximum Z extents. We found that 15 μm gave the most accurate results by visual inspection of the morphology. Some terminal points were then tagged cut points and dendrites were repaired.

The dendrite repair process created new dendritic sections starting at the identified cut points. Dendrite repair did not aim to recover the initial morphology, but rather recreated it in a statistical manner, under the assumption of statistical symmetry of the morphology. This method analyzed the behavior of intact branches as a function of branch order and euclidean distance from the soma. For each branch order, probability density clouds of branch continuation, bifurcation or termination were calculated in a series of concentric spheres (Sholl, 1953). At each cut point, the behavior of the branch was sampled according to the calculated probabilities. The factor governing the direction of the re-grown branches was adjusted to achieve final branches tortuosity comparable with our experimental data. To address neurite swelling artifacts at cut points, the diameters of the re-grown branches were set to the average diameter of the last section.

#### 2.2.5 Morphology diversification

We increased the variability of the reconstructed and repaired morphologies to ensure robust and invariant connectivity patterns (S. Hill et al., 2012; Ramaswamy et al., 2012). We followed a previously published method (Markram et al., 2015a) to generate a unique branching pattern for each morphology, while maintaining the general morphological and electrical structure for each m-type. In summary, branch lengths and rotations at each bifurcation point were varied according to random numbers drawn from Gaussian distributions with mean 0% and standard deviation 20% for branch lengths and mean 0° and standard deviation 20° for branch rotations. A sample of the resulting morphologies was visually validated, and we did not find significant alterations of their structure for any of the m-types (Figure S1).

We then applied a mix-and-match procedure to maximize the utilization of good morphological reconstruction data. This procedure divided dendrites from axons and allowed us to combine good dendritic reconstructions of TC and Rt dendrites from *in vitro* and *in vivo*-stained neurons and good axonal reconstructions from *in vivo*-stained neurons. *In vitro*-stained neurons typically lacked reconstruction of the full axon due to the slicing procedure and/or poor labeling. For each morphology, we manually annotated which dendrites and axons were to be kept. The decision in most cases depended on the labeling method (*in vitro* vs. *in vivo*).

To increase the probability that *in vivo*-stained morphologies and in particular the axons of TC and Rt morphologies were compatible with the microcircuit dimensions (see below) we duplicated and scaled the morphologies along their principal axis (Y-axis) by ± 2.5%.

### 2.3 Reconstructing the electrical diversity of neurons

#### 2.3.1 Electrophysiological data

The firing patterns of TC, Rt neurons and interneurons (INs) were characterized *in vitro* from brain slices of P14-35 GAD67-eGFP or C57Bl/6J mice and expert-classified into five electrical types (see Results). The detailed electrophysiological protocol has been published elsewhere (Iavarone et al., 2019a). Neurons were sampled from the ventrobasal complex of the thalamus (VPL and VPM nuclei) and the somatosensory sector of the reticular nucleus (Clemente-Perez et al., 2017; Lam and Sherman, 2011).

We used responses to step-like currents to build electrical models, ramp and noise currents to validate them (Iavarone et al., 2019a), along with excitatory postsynaptic-like currents (EPSC) injected into the dendrites. All the recordings were corrected for liquid junction potential by subtracting 14 mV from the recorded voltage.

#### 2.3.2 Neuron models

Multicompartmental conductance-based models employed 3D morphological reconstructions. Active ion currents and a simple intracellular calcium dynamics model were distributed in the somatic, dendritic and axonal compartments. Only the axonal initial segment (AIS) – not the complete axon – was modeled (Markram et al., 2015). The axons were substituted by a 60 μm stub constituted by two sections, five segments each. For each segment, the diameter was extracted from the original axon in order to preserve its tapering. Morphologies were divided into compartments of 40 μm maximal length. Specific membrane capacitance was set to 1 μF/cm^2^ and specific intracellular resistivity to 100 Ωcm.

#### 2.3.3 Ion channel models

We included ion current models whose kinetics were obtained from previously published ion current models or published experimental data. All ion channel models were corrected for liquid junction potential and for simulation at different temperatures whenever possible. Simulation temperature was always set to 34° C.

The details of the ion channel kinetics and calcium dynamics used for low-threshold bursting neurons (TC and Rt) have been described elsewhere (Iavarone et al., 2019a) and are summarized here. The type of ionic currents present in TC and Rt were: transient sodium current, delayed potassium current and low-threshold calcium from a previous model of TC neurons (Alain Destexhe et al., 1998); h-current model was built from published data (Budde et al., 1997; Iavarone et al., 2019a; McCormick and Huguenard, 1992; McCormick and Pape, 1990); persistent sodium, based on an existing models (Amarillo et al., 2014; Hay et al., 2011a) and published data (Parri and Crunelli, 1998), A-type transient potassium was taken from an existing model (Amarillo et al., 2014), based on published data (Huguenard and McCormick, 1992); high-threshold calcium was based on published models and data (Amarillo et al., 2014; McCormick and Huguenard, 1992), SK-type calcium-activated potassium, was taken from previous models and published data (Hay et al., 2011a; Köhler et al., 1996). Intracellular calcium dynamics was modeled with an exponential decay mechanism that linked low-threshold and high-threshold calcium currents to the calcium-activated potassium.

Since interneurons had firing patterns similar to cortical ones, we used the same ion channel models of the cortical microcircuit model (Markram et al., 2015a), which were based on existing models or published data. The type of ionic currents were transient sodium (Colbert and Pan, 2002), low-threshold calcium (Avery and Johnston, 1996), h-current (Kole et al., 2006), persistent sodium (Magistretti and Alonso, 1999), transient potassium (Korngreen and Sakmann, 2000), high-threshold calcium (Reuveni et al., 1993) and potassium Kv3.1 (Rettig et al., 1992). The reversal potential of sodium, potassium and h-current were set to 50 mV, −90 mV and −43 mV, respectively.

Ion channel models were distributed uniformly and with different peak conductance values for somatic, dendritic and axonal compartments, except for the h-current in the interneurons, whose distribution increased exponentially from the soma to the dendrites (Markram et al., 2015a).

#### 2.3.4 Optimization of neuron models

Five electrical models (e-models), corresponding to each electrical-type (e-type), were fitted using a multiobjective optimization algorithm using the Python library BluePyOpt (Iavarone et al., 2019a; Van Geit et al., 2016). The free parameters of the model were the peak conductances of the different mechanisms, parameters of the intracellular calcium dynamics (time constant of decay and percent of free calcium, *gamma*) and the reversal potential of the passive mechanism that contributes to the resting membrane potential. Each e-model was fitted with an exemplar morphology.

The optimization objectives were the electrical features extracted from the electrophysiological recordings. The detailed experimental protocol and the type of current stimuli and features are described elsewhere (Iavarone et al., 2019a), and summarized here. For all the e-types, two hyperpolarizing steps (−20/−40% and −120/−140% of the threshold current) were used to constrain passive properties (input resistance, resting membrane potential) and current activated by hyperpolarization, e.g. h-current (sag amplitude). Three levels of depolarizing steps (150%, 200%, 250% of the threshold current) were used to constrain firing pattern (adaptation index or inverse of the first and last interspike intervals, spike count, mean frequency) and spike shape-related features (action potential amplitude, depth of the after-hyperpolarization, action potential duration). All these protocols were applied in combination with a hyperpolarizing holding current (to reach stable membrane potential of −84 mV, after liquid junction potential correction).

When low-threshold bursting cells are hyperpolarized compared to their resting membrane potential and then stimulated, they fire stereotypical low-threshold bursts. One step (200% of firing threshold) on top of a hyperpolarizing current was therefore used to constrain the bursting response, while three depolarizing steps on top of a depolarizing holding current (to reach −64 mV) were used to constrain the tonic firing responses, as explained above. For reticular neurons, a new feature (*initburst_sahp*) was added for the afterhyperpolarization after the burst. For Rt and TC cells, two additional protocols without any current injection or only holding currents were used to ensure that the e-models were not firing without stimulus or with the holding current only.

Electrical features from the experimental recordings and model traces were extracted using the open source library eFEL (https://github.com/BlueBrain/eFEL).

We considered a model a good fit to the experimental data if all the feature errors (i.e. the Z-scores) were below 3.

#### 2.3.5 Quality assurance of morpho-electrical models

After fitting the five e-models, they were combined with the 92,970 morphologies generated as output of the morphology diversification step. An automated pipeline tested the e-models in combination with the different morphologies (me-models) and filtered out those that deviated significantly from the experimental electrical features. To decide which me-model was to be accepted, we used the repaired exemplar morphology (i.e. the morphology used during the optimization, after being repaired) as a benchmark: a me-model passed if it had all the feature errors were below 5 standard deviations of the repaired exemplar (Markram et al., 2015a). To account for the input resistance given by the different morphologies, we devised an algorithm, based on binary search, to find the appropriate holding and threshold current for each me-model (Iavarone et al., 2019a).

In addition, we first ran this pipeline on a small subset of the morphologies generated after morphology repair. In this way, we could visually inspect if the accepted me-models were generating biologically plausible firing behavior and the reasons why other me-models had high feature errors. In some cases, after inspecting the me-model voltage responses, we set less stringent criteria on some features, to ensure that we had enough different me-models for building the microcircuit. At the same time, we set more stringent criteria to reject me-models that were active without any input, since we did not find neurons that were spontaneously active in our experimental recordings.

### 2.4 Measuring neuron density

#### 2.4.1 Immunohistochemistry of Rt and VPL for cell counting

We complemented the neuron densities values from the Blue Brain Cell Atlas (Erö et al., 2018; RRID:SCR_019266) by counting neurons in adult mouse brain slices. The brain was cryosliced at 50 μm on the sagittal plane and stained following standard immunohistological procedures with antibodies anti-GABA (for inhibitory neurons), anti-NeuN (for neurons) and DAPI (for all cells), using an existing protocol (Markram et al., 2015). The slices were imaged with a confocal microscope (Zeiss, 710). The immunohistology and imaging of the region of interest (ROI) was completed for one P21 C57B1/6J mouse.

#### 2.4.2 Semi-automated cell counting and cell densities

The images were aligned to the Allen Reference Atlas to create proper boundaries for the Rt and VPL. We used Imaris® software (Bitmap) to create the ROI, for counting the neurons and to estimate the volume for density calculation. For a chosen ROI, the software detected the difference of signal intensity, created a 3D shape around the detected cells and extracted statistics (e.g. count, positions) following given parameters. These parameters were defined by running multiple trials so that the results from semi-automated cell counting were as close as possible to those from manual cell counting. The semi-automated counting method results in very low error rates compared to manual counting (2.25%) and is less time consuming. A 3D shape of the entire ROI was created in order to extract the volume for density calculation. Neuron densities were calculated as the ratio between neuron counts in a ROI and the volume as calculated in Imaris for each slice. For modeling we used the average cell densities for Rt and VPL neurons.

### 2.5 Reconstructing the dimensions and structure of a thalamoreticular microcircuit

Since the thalamus does not have a clear laminar structure, we approximated a thalamic microcircuit as a cylindrical volume having its base parallel to a portion of the Rt and its vertical dimension (y-axis) running through the VPL and Rt (see Fig. 3A).

The horizontal dimensions of the microcircuit were calculated from the density of dendritic fibers at the center of the circuit, following an approach published previously (Markram et al., 2015a). For each m-type, we began by considering all the morphologies (after repairing them) that had their somata located within 25 μm from the circuit center on the horizontal plane (XZ). We then increased the maximal distance in steps of 25 μm which resulted in an increase of dendritic densities at the center. The microcircuit horizontal dimension (radius) resulting from this process was 294 μm, corresponding to the distance where 95% of the asymptotical maximal density of reticular neuron dendrites was reached. As a comparison, considering only thalamocortical cell morphologies would have resulted in a circuit with radius 125 μm, while considering only interneurons the radius would have been 279 μm.

We used hexagonal boundaries with the same area as the resulting circle to facilitate tiling of multiple microcircuits, while keeping asymmetrical edge effects minimal. The resulting side of the hexagon was 323 μm and the longest diagonal (vertex-to-vertex) measured 646 μm.

To calculate the vertical dimension of the microcircuit, we extracted a 3D subvolume within the VPL and the Rt. We started from the thalamus parcellation of the Allen Brain Atlas version 3 (25 μm resolution) (Goldowitz, 2010). A spherical coordinate system was fitted to the volume of the Rt, which can be approximated by a spheroidal surface. We chose a ROI located approximately in the middle of the VPL nucleus and computed the probability distribution of widths in the ROI for the VPL and Rt. The widths were calculated along the radius of the spherical coordinate system. The resulting thickness corresponds to the median of the distributions, which was 550 μm for the VPL and 250 μm for the Rt.

#### 2.5.1 Soma positions and me-type model assignment

The horizontal and vertical extents resulted in a microcircuit having the shape of a hexagonal prism, that was 646 μm wide (at the widest point) and 800 μm high; 69% of the volume was occupied by the VPL and 31% by the Rt. This volume was then populated by defining somata positions according to the experimentally measured neuron densities in the Rt and VPL. The positions were distributed according to an algorithm based on Poisson disc sampling (Bridson, 2007; Tulleken, 2009). This algorithm avoids clustering normally obtained with sampling according to uniform distributions, by using a parameter for the minimum distance between points. To calculate the minimum distance, we used the cell densities to calculate the expected number of cell positions per voxel. Each soma position was assigned an m-type according to the excitatory/inhibitory fractions and an electrical model in agreement with the me-types composition (Fig. 3). Moreover, each position was associated with a random rotation around the y-axis to be applied to each morphology.

#### 2.5.2 Morphology placement

Our pool of experimental morphologies and the ones derived from the morphology diversification process contained morphologies with different sizes and shapes. Moreover, it contained TC morphologies whose somata was not located in the VPL nucleus and Rt morphologies whose axons were not arborizing in the VPL nucleus. We adapted a placement scoring algorithm (Markram et al., 2015) to ensure that each position was assigned a suitable morphology considering its geometrical properties and the microcircuit vertical dimension.

We thus defined placement rules that took into account the known properties of Rt and TC neurons’ arborizations relative to the anatomical boundaries of thalamic nuclei (Harris, 1986; Pinault et al., 1995). Each reconstruction of TC and Rt neuron morphologies was manually annotated according to the placement rules. For TC cells, we identified the axonal arborization projecting to the Rt (and that should be located in the Rt part of the model). For Rt cells, the densest part of axonal arborization was annotated, which should be located in the VPL. For IN, the only constraint is that the full morphology should be contained within the VPL and not crossing into the Rt (Morgan and Lichtman, 2020). Each annotation was automatically carried over during the unraveling, repairing and diversification steps. Moreover, we included a stricter rule to avoid that Rt morphologies were located outside the top of the circuit boundary, with a 30 μm tolerance.

Given the placement rules, each morphology was assigned a score based on the microcircuit position and the constraints set by the placement rules (Markram et al., 2015a).

#### 2.5.3 Generating different microcircuit instances

We created five different microcircuit instances to assess the model robustness to different input parameters. The experimentally-measured cell densities were jittered by +/-5%, resulting in microcircuits with different total number of neurons and number of neurons for each m-type (see Fig. 4).

### 2.6 Reconstructing the synaptic connectivity of a thalamoreticular microcircuit

#### 2.6.1 Connectivity based on morphological appositions

After placing the morphologies in the 3D microcircuit volume we generate the first version of the connectivity by detecting zones of geometrical overlap (“touches”) using an existing touch detection algorithm (Kozloski et al., 2008; Markram et al., 2015a). Briefly, this algorithm sub-divided the circuit 3D space into sub-volumes ensuring that each sub-volume contained the same amount of data, i.e. the same number of morphological segments. Each sub-volume was processed in parallel on different cores and written in parallel to disk. All geometrical overlaps were considered as touches if their distance was smaller or equal to 1 μm (“touch distance”).

Touches were then filtered according to biological rules: touches were allowed between all m-types, except between interneurons and reticular cells, because interneurons are only located in the thalamus and are not expected to have neurites extending into the reticular nucleus (Morgan and Lichtman, 2020). Touches between VPL neurons were removed, in agreement with experimental findings showing that excitatory connection between TC neurons disappear during development (Lee et al., 2010).

Interneurons also form axonal and dendritic inhibitory synapses (Acuna-Goycolea et al., 2008; Cox and Sherman, 2000; Zhu and Heggelund, 2001). For all other m-type combinations, touches formed between presynaptic axons, postsynaptic dendrites and somata.

The same algorithm was used to detect touches between Rt_RC dendrites, i.e. the locations of putative gap junctions. Since gap junctions are established with close appositions of cell membranes, we used a touch distance of 0 μm in this case.

At the end of this process the resulting contacts (or “appositions”) are normally higher compared to experimental findings and are pruned further to arrive at the final functional synapses (Reimann et al., 2015).

#### 2.6.2 Determining functional synapse positions

We employed an existing algorithm to decide which appositions were to be pruned according to biological constraints (Reimann et al., 2015). The main constraints were the experimental bouton densities (number of boutons / axonal length) from 3D neuron reconstructions (n=9 TC axons and n=2 Rt axons) and the coefficient of variation of number of synapses per connections (i.e. the number of functional synapses, between a pair of neurons) from presynaptic INs and post-synaptic INs and TCs (Morgan and Lichtman, 2020).

In the first two steps, the algorithm tried to match the distribution of synapses per connection, using the coefficient of variation of appositions per connections and the coefficient of variation of synapses per connection. Then, in step 3, it compared the current bouton density to the target value and removed multi-synaptic connections until the target value was matched. The number of synapses per connections, *N_func_*, was predicted from the number of appositions per connections (*N_app_*) resulting from the previous steps, similarly to uncharacterized pathways in cortical microcircuitry (Reimann et al., 2015). *N_func_* was predicted from *N_app_* according to a simple formula (*N_func_* = 1 ⋅ *N_app_*) for each m-type to m-type connection. We used a generalized coefficient of variation for *N_func_* of 0.9 for all connections, c extracted from published data (Morgan and Lichtman, 2020, Fig. 3B). The coefficient of variation was combined with the predicted *N_func_* to calculate its standard deviation, as detailed in (Reimann et al., 2015). At the end of this pruning process, we verified that the bouton densities in the model matched the experimental ones (see Fig. 4A). The shape of a geometric distribution for *N_func_* was a prediction from our touch detection process.

#### 2.6.3 Connections from lemniscal and corticothalamic afferents

We followed an approach similar to the generation of thalamic input to the cortical microcircuit model (Markram et al., 2015) to model afferent synapses in the thalamus from the sensory periphery (medial lemniscus) and from the cortex. The algorithm uses volumetric bouton densities and the morphologies already placed in a circuit to map synapses from afferent “virtual” fibers to postsynaptic morphologies.

We built medial lemniscus (ML) and corticothalamic (CT) afferents separately for one microcircuit. Since data for lemniscal innervation in the mouse VPL was not available we calculated volumetric bouton density from data of mouse VPM (Takeuchi et al., 2017), see Fig. 4C for the exact values. Volumetric bouton densities for the CT pathway were derived from known proportions between CT synapses and other synapses onto TC and Rt neurons, as found in electron microscope investigations (Çavdar et al., 2011; Mineff and Weinberg, 2000) (see Fig. 4C).

Each synapse was assigned a virtual ML or CT fiber. We estimated a number of 2,601 ML fibers; this number took into account the ratio between the putative number of neurons from the dorsal column nuclei projecting to the thalamus (Shishido and Toda, 2017) and the number of neurons in the VPL (see (Jones, 2007) for a similar calculation). The number of CT fibers was 75,325, about ten times the number of thalamocortical fibers in a microcircuit (Crandall et al., 2015; Monconduit et al., 2006; Sherman and Koch, 1986).

To take into account the correlation between synaptic inputs onto postsynaptic neurons innervated from the same afferent fiber, the mapping between postsynaptic synapses and fibers took into account their respective positions, i.e. synapses that were closer together were more likely to be innervated by the same presynaptic fiber. As in the neocortical microcircuit model (Markram et al., 2015), the probability (*P*) that a synapse was assigned to a fiber depended on the distance between the synapse and the fiber:

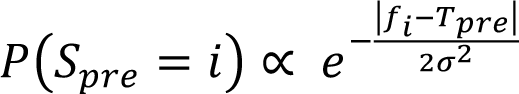

where *S_pre_* represents the mapping of a synapse *S* to the presynaptic fiber *i*, *T_pre_*is its spatial location, *f_i_* the spatial location of fiber *i* and ⌠ denoted the degree of spatial mapping, that was set to 25 μm.

### 2.7 Modeling synapse physiology

#### 2.7.1 Stochastic synaptic transmission and short-term plasticity

We used existing models of stochastic transmission at excitatory and inhibitory synapses (Markram et al., 2015). They consisted of a 2-state Markov process, with recovered and unrecovered states. When a pre-synaptic event occurs (pre-synaptic spike or spontaneous release) the synapse will release if it is the recovered state. If there is release, the synapse will transition to the unrecovered state. The ensemble average response is equivalent to the phenomenological Tsodyks-Markram model (Fuhrmann et al., 2002; Tsodyks and Markram, 1997). The underlying assumptions were derived from the classical model of quantal synaptic release, in which each synapse is assumed to have *N* independent release sites, each has a probability *p* of releasing a single quantum *q* (del Castillo and Katz, 1954; Korn and Faber, 1991). The number of release sites was assumed to be equivalent to the number of synapses per connection (Markram et al., 2015). The detailed implementation of the synapse models can be downloaded from the neuron model packages in the Neocortical Microcircuit Portal (rrid:SCR_022032; (Ramaswamy et al., 2015)).

We modeled short-term synapse plasticity with depressing (E2 and I2) and facilitating synapses (E1), see Fig. 5. In our experimental recordings, in agreement with experimental findings, all existing intrathalamic (between TC, Rt neurons and INs) and lemniscal connections were depressing (Cox et al., 1997; Gentet and Ulrich, 2003; Miyata, 2007; Mo et al., 2017; Simko and Markram, 2021), while corticothalamic ones were facilitating (Crandall et al., 2015; Jurgens et al., 2012; Landisman and Connors, 2007; Miyata, 2007; Reichova and Sherman, 2004). When sufficient experimental paired recordings data were available, the parameters of the Tsodyks-Markram model of short-term synaptic plasticity were fitted (see above). The data used for fitting were the excitatory postsynaptic potentials (EPSPs) or inhibitory postsynaptic potentials (IPSPs) peaks amplitudes (or EPSCs/IPSCs in the case of voltage-clamp recordings), evoked by stimulating the presynaptic cell with a train of eight pulses followed by a recovery pulse (see Fig. 5A). The parameters were: U - release probability, D - time constant of recovery from depression, and F - time constant of recovery from facilitation. Postsynaptic data were filtered and deconvolved for easier automatic identification of the peaks (Barros-Zulaica et al., 2019). A multi-objective optimization algorithm was used to find the values for U, D and F (Van Geit et al., 2016).

**Figure 5.**
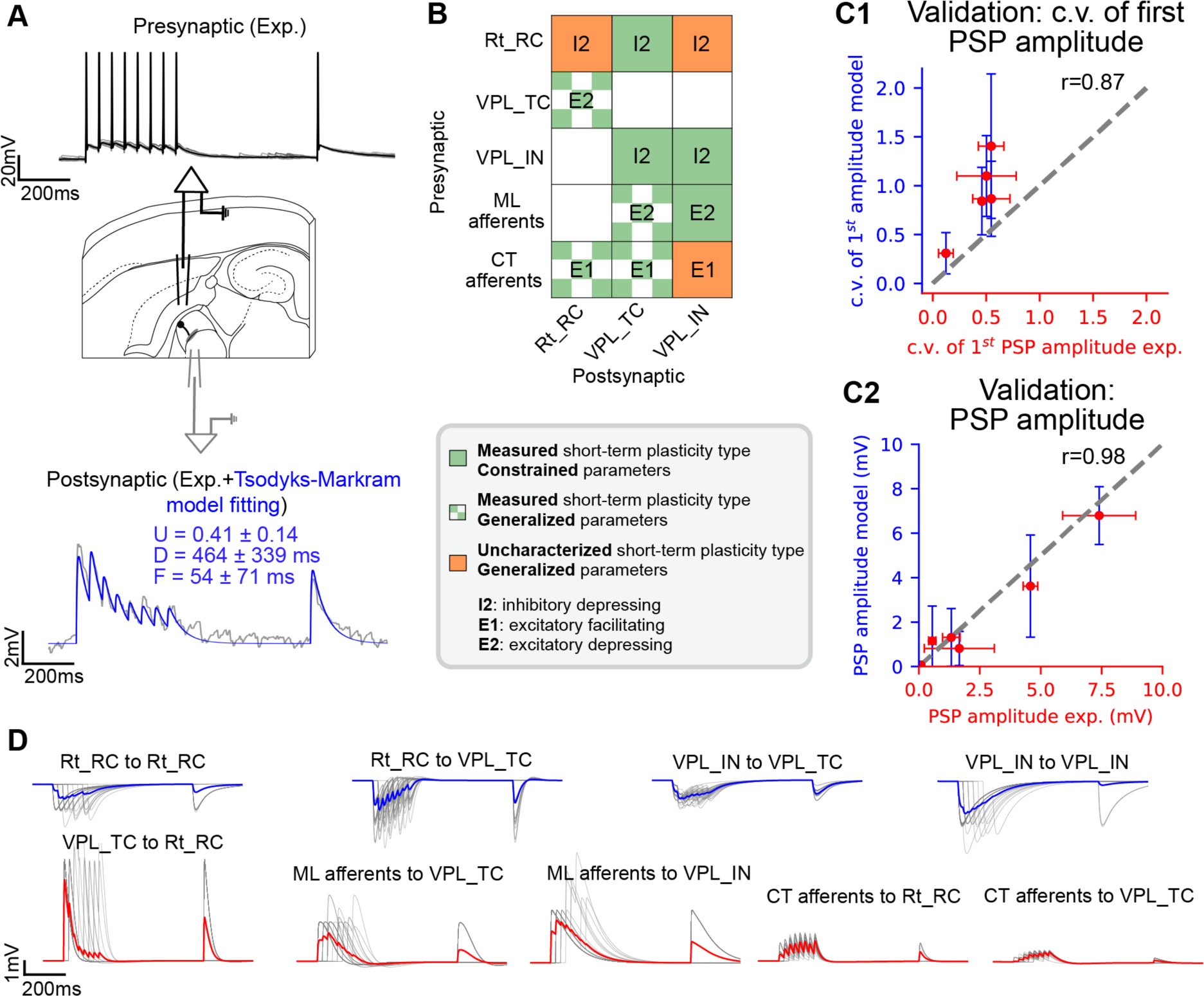
Reconstructing and validating short-term synaptic plasticity and postsynaptic potential (PSP) amplitudes. (**A**) Top: illustration of *in vitro* paired recordings, used to constrain the parameters of the Tsodyks-Markram model of short-term plasticity. A presynaptic neuron (black pipette) was stimulated with 8 pulses at 40 Hz followed by a recovery stimulus. The response in a postsynaptic neuron (gray pipette) was recorded and used to constrain the model parameters (U, D, F). (**B**) Map of short-term plasticity types in the model. Green: in-house experimentally-characterized pathways (as in A), green checked: pathways for which the synapse type was derived from literature and parameters were generalized from similar pathways (see Methods for details), orange: pathways for which paired recording data was not available. (**C1**) Validation of the coefficient of variation (c.v.) of first PSP amplitudes, quantifying the trial-to-trial variability for 5 *in vitro* characterized pathways (see Table 4.3). (**C2**) Comparison of PSP amplitudes in the model for 7 characterized pathways in-house or in the literature (see Table 4.2). Dots and error bars show mean and standard deviation, dashed line shows the regression fit. (**D**) Example of *in silico* paired recordings for different pathways. For each pathway, the somatic membrane potential of the postsynaptic neuron is shown (gray: trials; blue and red: mean traces for inhibitory or excitatory connections). All recordings are shown on the same scale for easy comparison of PSP amplitudes in different pathways.

Data to fit the UDF parameters was available for some of the pathways: Rt neurons to TCs, IN to TCs, INs to INs and ML to INs connections; for all the other pathways we followed these generalization rules:

TC to Rt synapses were shown to be strong, reliable and depressing (Gentet and Ulrich, 2003). We used parameters for L4Exc to L4Exc connections from the neocortical microcircuit model (Markram et al., 2015) as they had the highest release probability (analogous to the U value in the case of depressing synapses (Ecker et al., 2020).
All uncharacterized inhibitory to inhibitory synapses (i.e. Rt to Rt and Rt to IN) had the same dynamics of an inhibitory-inhibitory characterized pathway (i.e. IN to IN).
CT synapses onto first order thalamic nuclei (e.g. VPL, VPM, dorsal part of the lateral geniculate complex) have been consistently reported to be facilitating. As we did not have paired recordings to estimate synapse parameters for CT to TCs, CT to INs and CT to Rt pathways, we took parameters from excitatory facilitating synapses (E1: L5TTPC-L5MC, Markram et al., 2015).
ML inputs to first order sensory thalamic nuclei (e.g. VPM) were shown to be depressing (e.g. see Mo et al., 2017; Reichova and Sherman, Castro-Alamancos, 2002; Miyata, 2007), as shown in our ML to INs recordings. We thus extrapolated the parameters for ML to TC connections from ML to INs ones (for which data was available).

Synapse dynamic parameters in the model were different for each synapse and drawn for truncated Gaussian distributions.

Spontaneous miniature potentials were modeled as independent Poisson processes at each synapse that triggered release at low rates (0.01 Hz).

#### 2.7.2 Synapse models

Excitatory synaptic transmission was modeled with AMPA and NMDA receptor kinetics, and GABA_A_ receptors were used for inhibitory connections. The rise and decay phases of the currents were described using mono-exponential functions. We used time constants from thalamic experiments performed at 34-35 degrees C, when available, or from cortical synapses models when thalamic-specific ones were missing. The rise time and decay time constants for AMPA receptors were 0.2 ms and 1.74 ms, respectively (Häusser and Roth, 1997). For TC to Rt connections the AMPA decay time constant was 1.58 ms and CT afferents to Rt was 2.74 ms (Deleuze and Huguenard, 2016). The rise and decay time constants of the NMDA component were 0.29 and 43 ms (Sarid et al., 2007). The magnesium concentration was set to 1 mM (Jahr and Stevens, 1990) and the reversal potential of the AMPA and NMDA currents was 0 mV. Experimentally measured ratios of NMDA and AMPA conductances were gathered from the literature and are summarized in Table 4.1 (Arsenault and Zhang, 2006; Deleuze and Huguenard, 2016; Miyata and Imoto, 2006).

Inhibitory synaptic transmission was modeled with GABA_A_ receptor kinetics. The rise and decay time constants were 0.2 ms and 8.3 ms, respectively (Markram et al., 2015a). The reversal potential of GABA_A_ current was set to −82 mV for all inhibitory pathways, except for connections onto postsynaptic TC neurons, where it was −94 mV, consistent with lower chloride reversal potentials in TC compared to Rt neurons (Ulrich and Huguenard, 1997).

#### 2.7.3 Constraining synapse conductance values

Synaptic conductance values were optimized by performing *in silico* paired recordings to match the postsynaptic potential (PSP) amplitudes measured experimentally whenever data was available, similarly to other morphologically detailed models (Ecker et al., 2020; Markram et al., 2015a). For each pathway, 50 neuron pairs were simulated, and each pair was recorded for 30 trials. Experimentally characterized values in rodents are summarized in Table 2. For all other pathways, we extrapolated the quantal synapse conductances from similar pathways, according to the same generalization principles applied for short-term plasticity parameters (see Table 1).

**Table 1.** Synapse kinetics and short-term plasticity parameters. Synaptic parameters for all pathways in the model. Quantal synaptic conductance *g_syn_*(in nanosiemens nS), *Ι_d_* is the decay time constant of AMPA and GABA_A_ currents for excitatory and inhibitory connections. *U_SE_* (utilisation of synaptic efficacy, analogous to release probability), *D* (time constant of recovery from depression), *F* (time constant of recovery from facilitation) are the short-term plasticity parameters. Values are expressed as mean ± standard deviation. All the parameters were fitted to in-house paired-recordings or generalized from similar pathways (Fig. 5 and Methods for details).

| Pathway | Synapse type | $g_{\text{syn}}$ (nS) | $\tau_d$ (ms) | NMDA/AMPA ratio | $U_{\text{SE}}$ | D | F |
| --- | --- | --- | --- | --- | --- | --- | --- |
| Rt_RC to Rt_RC | Inh. Dep. | $0.9 \pm 0.23$ | $8.3 \pm 2.2$ | NA | $0.41 \pm 0.14$ | $464 \pm 339$ | $54 \pm 71$ |
| Rt_RC to VPL_TC | Inh. Dep. | $1.1 \pm 0.4$ | $8.3 \pm 2.2$ | NA | $0.32 \pm 0.18$ | $352 \pm 46$ | $2 \pm 209$ |
| Rt_RC to VPL_IN | Inh. Dep. | $0.9 \pm 0.23$ | $8.3 \pm 2.2$ | NA | $0.41 \pm 0.14$ | $464 \pm 339$ | $54 \pm 71$ |
| VPL_TC to Rt_RC | Exc. Dep. | $2.8 \pm 0.1$ | $1.58 \pm 0.26$ | 0.57 | $0.86 \pm 0.09$ | $671 \pm 17$ | $17 \pm 5$ |
| VPL_IN to VPL_TC | Inh. Dep. | $0.4 \pm 0.4$ | $8.3 \pm 2.2$ | NA | $0.47 \pm 0.18$ | $137 \pm 46$ | $239 \pm 209$ |
| VPL_IN to VPL_IN | Inh. Dep. | $2.7 \pm 0.4$ | $8.3 \pm 2.2$ | NA | $0.41 \pm 0.14$ | $464 \pm 339$ | $54 \pm 71$ |
| ML to VPL_TC | Exc. Dep. | $1.15 \pm 0.12$ | $1.74 \pm 0.18$ | 0.41 | $0.3 \pm 0.21$ | $2350 \pm 315$ | $1 \pm 2$ |
| ML to VPL_IN | Exc. Dep. | $1.15 \pm 0.12$ | $1.74 \pm 0.18$ | 0.41 | $0.48 \pm 0.21$ | $690 \pm 315$ | $57 \pm 53$ |
| CT to Rt_RC | Exc. Fac. | $0.16 \pm 0.016$ | $2.74 \pm 0.25$ | 0.99 | $0.09 \pm 0.12$ | $138 \pm 211$ | $670 \pm 830$ |
| **CT to VPL_TC** | Exc. Fac. | 0.16±0.01<br>6 | 1.74±0.18 | 1.91 | 0.09±0.12 | 138±211 | 670±830 |
| **CT to VPL_IN** | Exc. Fac. | 0.16±0.01<br>6 | 1.74±0.18 | 0.99 | 0.09±0.12 | 138±211 | 670±830 |

**Table 2.** Coefficient of variation (CV) of first PSP amplitudes. CV of first PSP amplitudes values as characterized experimentally through *in vitro* paired recordings. Values are reported as mean ± standard deviation (of multiple pairs). (Related to Fig. 5C1).

| Presynaptic | Postsynaptic | CV 1<sup>st</sup> PSP amplitude, experiment (mV) | CV 1<sup>st</sup> PSP amplitude, model (mV) | Data source |
| --- | --- | --- | --- | --- |
| **Rt_RC** | **VPL_TC** | 0.4600 (n=1) | 0.8424 ± 0.3450 (n=47) | In-house |
| **VPL_TC** | **Rt_RC** | 0.1232 ± 0.0686 (n=11) | 0.3089 ± 0.2112 (n=43) | (Gentet and Ulrich, 2003) |
| **VPL_IN** | **VPL_TC** | 0.5479 ± 0.1744 (n=4) | 0.8663 ± 0.386 (n=22) | In-house |
| **VPL_IN** | **VPL_IN** | 0.5028 ± 0.2783 (n=10) | 1.0993 ± 0.4132 (n=49) | In-house |
| **ML** | **VPL_IN** | 0.5466 ± 0.1195 (n=1) | 1.4047 ± 0.7389 (n=49) | In-house |

### 2.8 Modeling gap junctions

Along with excitatory and inhibitory chemical synapses, the microcircuit included detailed gap junction (GJ) connectivity established between the dendrites of Rt neurons. We used the same touch detection algorithm described above to find appositions between Rt neuron dendrites and somata. Since we did not have any experimental data on the number of GJs between connected neurons or the density of GJs (number of GJs per unit length of dendrite or volume), in this first draft we randomly removed a certain fraction of GJs until we matched data on neuron divergence (Fig. 6A). To analyze the number of coupled neurons and their spatial properties (Fig. 6), we reproduced the experimental protocol (S.-C. Lee et al., 2014), by analyzing a sample of 33 Rt neurons in a 90 μm vertical slice located at the center of the microcircuit.

**Figure 6.**
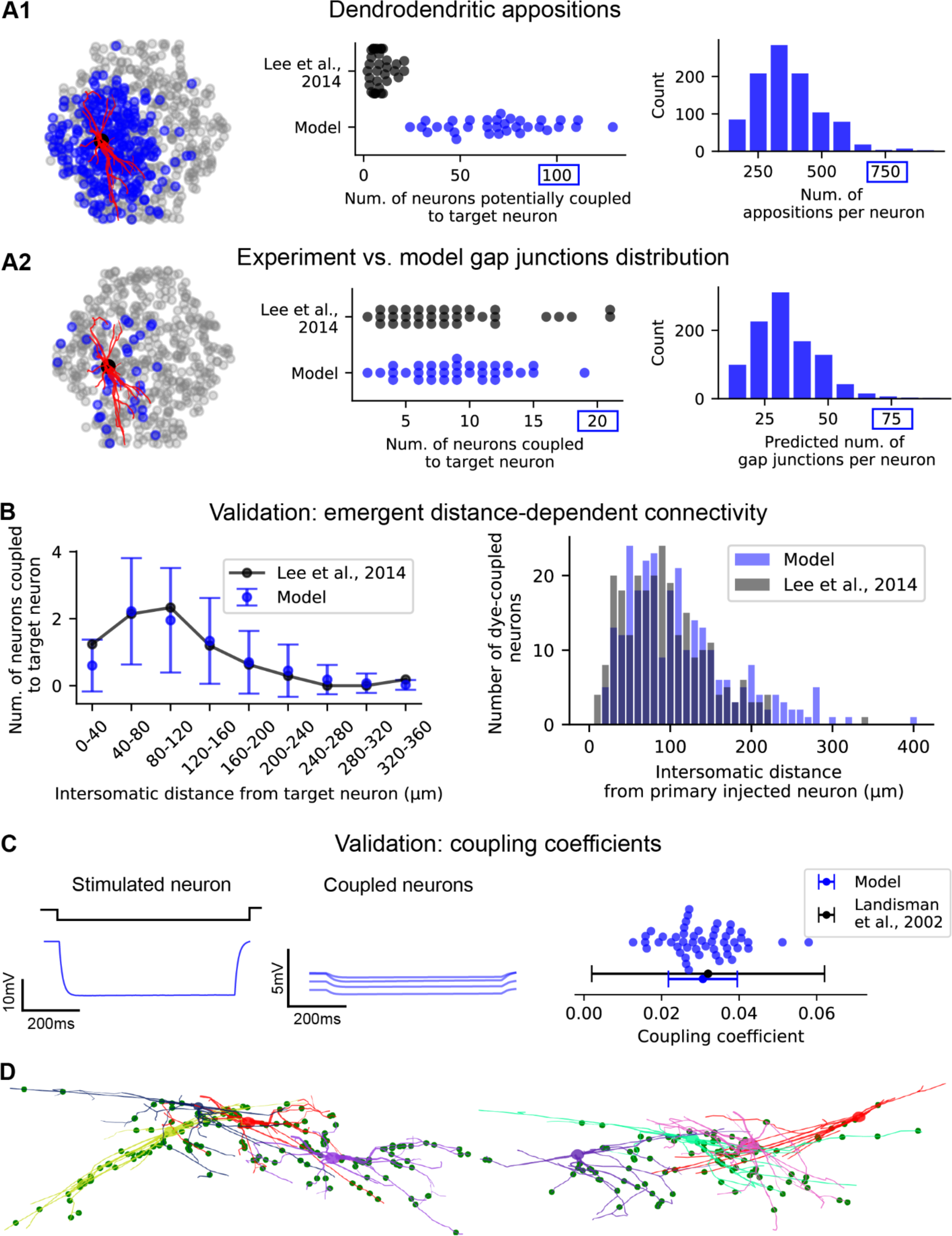
Reticular nucleus gap junction connectivity is predicted by dendrodendritic overlap. (**A1**) Potential connectivity based on dendrodendritic (and somatic) appositions between Rt_RC dendrites. Left: view of the microcircuit from the Rt side showing the location of a sample of 500 Rt_RC neurons (gray dots), a “target” Rt_RC morphology (2D projection, dendrites in red) and the location of Rt_RC neurons connected to the target Rt_RC (blue dots). Middle: neuron divergence (number of postsynaptic neurons) in the model and literature (N=33 for both experiment and model). Each dot represents one target neuron. Right: distribution of potential connectivity divergence (number of appositions per neuron, for a sample of 1000 Rt_RC neurons in the model). (**A2**) Predicted gap junctions (GJ) after randomly removing dendrodendritic appositions to match average GJ divergence. Left: as in A1. Middle: neuron divergence in the model matches experimental findings. Right: the resulting GJ divergence is reduced by an order of magnitude. Note different maximal values in A1 and A2. (**B**) Validations of distance dependent GJs connectivity. Right: *In silico* dye-injections were performed in the model to reproduce dye-coupling experiments (n=500 neurons in the model, n=33 experiment, mean and standard deviation are shown). (**C**) Validation of GJs functional properties. Left: example of *in silico* paired recordings, where a Rt_RC is stimulated with a hyperpolarizing current step, its somatic potential is recorded, along with the somatic potential of all coupled neurons (only a sample is shown). The ratio of the voltage response between a coupled neuron and the stimulated neuron is the coupling coefficient (CC). Right: comparison of CC values in the model (n = 50 pairs, each one represented by a dot) with paired recordings from the literature. Dots: mean, error bars: standard deviation. (**D**) Resulting GJ connectivity. Example of clusters of 4 Rt_RC neurons coupled by GJs and GJ locations. Each neuron morphology is represented by a different color, axons are omitted for clarity. Green dots show the detailed morphological location of GJs that each of the neurons receive from the 3 others and from other Rt_RC neurons not shown here.

**Figure 7.**
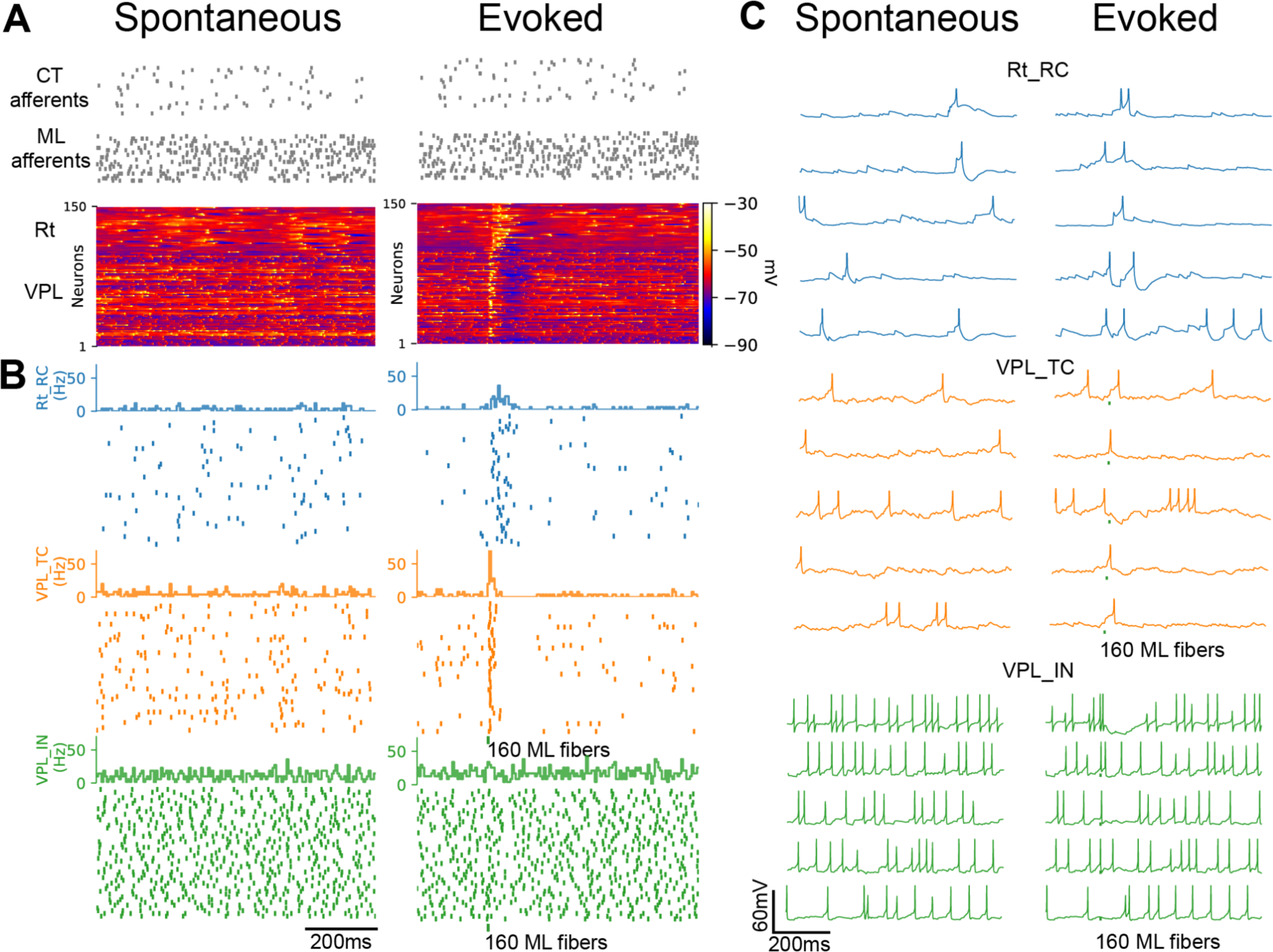
Spontaneous and sensory-evoked activity in the simulated thalamoreticular microcircuit (*in vivo* wakefulness-like condition) Simulating wakefulness-like spontaneous and evoked activity. (**A**) Population voltage raster showing the membrane potential of a sample of 50 active neurons per m-type. Each row represents the activity of a single neuron and is sorted according to microcircuit depth. Note the uncorrelated membrane fluctuations along the depth of the microcircuit and the increased responses with the sensory stimulus in both Rt and VPL and visible hyperpolarization in the VPL after the stimulus. (**B**) Firing rate histograms and spike rasters showing uncorrelated spiking activity in all m-types. VPL_IN have higher firing rates due to their lower spiking threshold. Note that Rt_RCs show increased activity for a longer time after the stimulus compared to VPL neurons. (**C**) Exemplar single cell recordings for 5 neurons for each m-type. Note the variability in spiking and subthreshold activity between different cells of the same m-type. For instance, VPL neurons respond to the stimulus with spikes, spikes followed by hyperpolarization or hyperpolarization alone. In this and following figures spikes are truncated at −25 mV.

Functionally, GJ were modeled as conductances that coupled the membrane potential of the adjacent morphological compartments (simple resistors). We predicted the value of gap junction conductance for all gap junctions and validated their functional properties by comparing coupling coefficient values with experiments (Haas et al., 2011; Landisman et al., 2002; S.-C. Lee et al., 2014; Long et al., 2004).

Once the structural properties of gap junctions-coupled neurons were validated, we performed *in silico* paired recordings and measured the coupling coefficients for each pair of neurons. We found that the mean coupling coefficients in the model compared well with the experiments for gap junction conductance values of 0.2 nanosiemens (nS).

After adding gap junctions to the circuit, the input resistance of the neurons changed. To guarantee that the electrical properties of the neurons did not change, thus changing the responses to synaptic inputs, we devised an algorithm to compensate for the change in input resistance (Amsalem et al., 2016). The algorithm changed the conductance of the leak current (*g_pas_*) to restore the input resistance of the neuron before adding gap junctions. This compensation resulted in a different *g_pas_*value for each neuron.

### 2.9 Simulation methods and conditions

#### 2.9.1 Simulation software and high-performance computing resources

The reconstructed microcircuit was simulated using software based on the NEURON simulation package ((Hines and Carnevale, 1997); RRID:SCR_005393). A collection of tools and templates were written in order to handle simulation configuration, *in silico* network experiments and to save the results. We used the CoreNEURON simulator engine (Kumbhar et al., 2019), which has been optimized for efficient large-scale simulations. A typical simulation run of a microcircuit for 3,500 ms of simulation time took ∼45 minutes on 16 Intel Xeon 6140 CPUs (288 cores, with HyperThreading enabled).

#### 2.9.2 Simulating in vivo-like conditions

To simulate spontaneous activity in *in vivo* wakefulness-like states, we activated lemniscal and CT fibers with Poisson spike trains at 25 and 4 Hz, respectively. We lowered the extracellular calcium concentration from 2 mM (*in vitro*-like conditions) to 1.2 mM, with the effect of reducing synapse release probabilities and PSPs amplitudes (Markram et al., 2015a). PSPs were dependent on calcium concentration in the same way as uncharacterized pathways as previously published (Markram et al., 2015, Fig. S11 – Intermediate [Ca^2+^]_o_ dependence). This condition was used in all simulations of *in vivo* wakefulness-like activity, if not stated otherwise.

To simulate cortical UP and DOWN states, we removed the background activity from the CT afferents (Figg. 11, 15).

#### 2.9.3 Simulating lightly-anesthetized in vivo-like conditions

To simulate lightly anesthetized *in vivo-*like states, we followed the same methodology as for *in vivo*-like conditions, however the spontaneous firing induced at the thalamus through the ML afferents was reduced from 25 Hz to 10 Hz to reflect the presumed hyperpolarizing influence of the anesthetic (Fig. 9).

**Figure 8.**
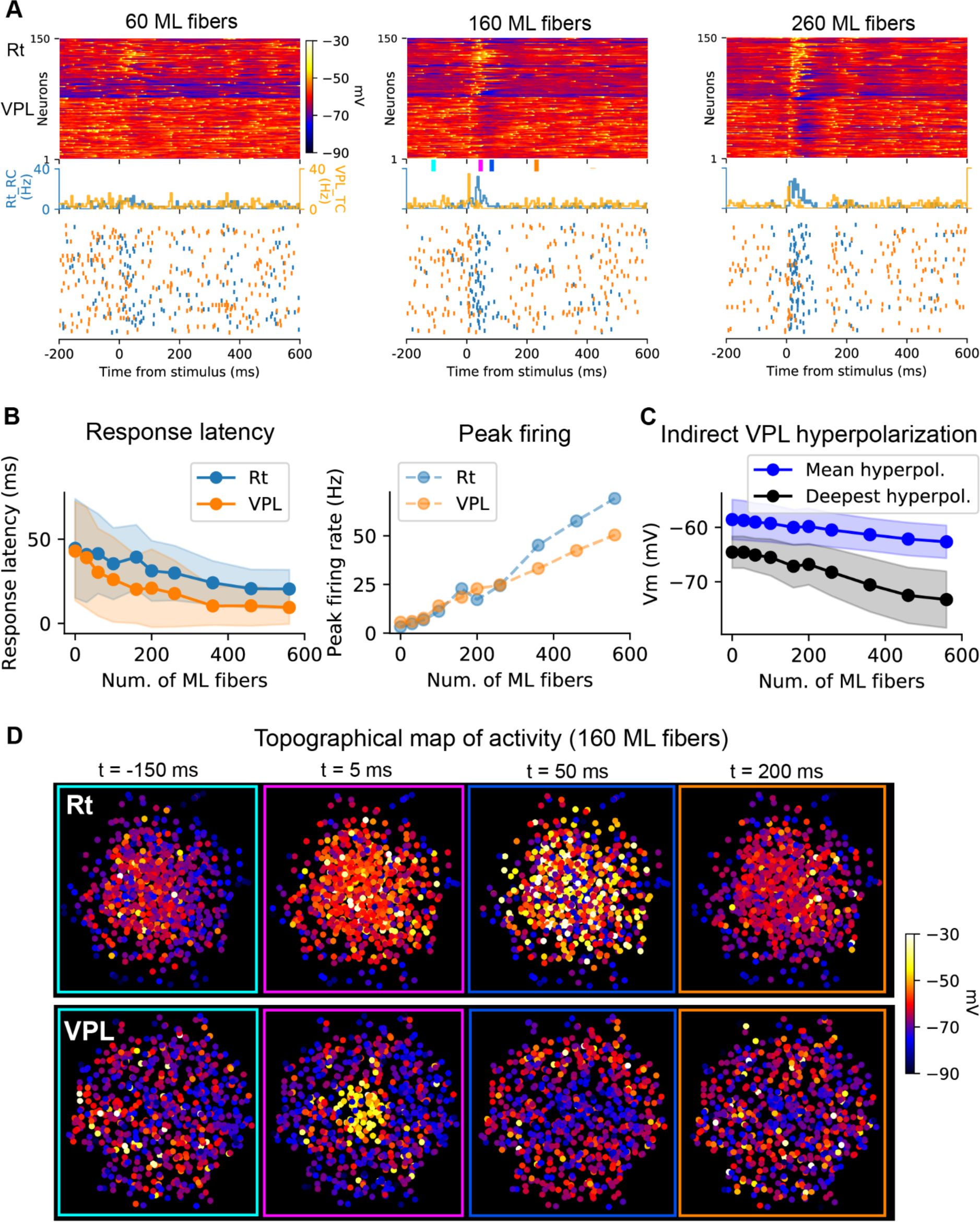
Stimulus-dependent recruitment of reticular nucleus neurons and surround inhibition of thalamic neurons (*in vivo* wakefulness-like condition). (**A**) Simulated sensory inputs with brief activation of increasing numbers of medial lemniscal fibers (ML). Top: voltage rasters show Rt and VPL responses of a sample of 150 active neurons, sorted by their vertical position in the microcircuit. Bottom: spiking responses (firing rate histograms and spike rasters). Note that stimulus-evoked responses in the VPL as well as the following hyperpolarization increase with increasing stimulus size. (**B**) Stimulus-response curves. Response latency decreases with stimulus size (left), while peak firing rates increase the VPL, as well as in the Rt (right). Mean (lines) and standard deviation (shaded areas) are shown. The peak firing rate is calculated in the 100 ms following the stimulus and response latency as the time to first spike after the stimulus (n=1,000 neurons). (**C**) Stimulus-dependent hyperpolarization in the VPL. The mean and deepest hyperpolarization in VPL cells are shown (lines: mean, shaded areas: standard deviation). Note that with increasing stimulus size more VPL cells are inhibited by Rt neurons and the hyperpolarization becomes stronger. The hyperpolarization is calculated in a time window of 40-200 ms after the stimulus (same sample as in B, n=1,000). (**D**) Topographical activity in a slice through the VPL and Rt showing the average membrane potentials at different time windows before, during and after the stimulus, as indicated by colored ticks in A (middle panel), 160 fibers were activated. Time windows of 10 ms starting at the time indicated were used for the average activity. Note that increased activity is confined to the central part of the VPL in response to the stimulus (t=5 ms), which triggers spiking activity in the Rt (t=50 ms) in central as well as in peripheral neurons. This result suggests that the Rt has larger receptive fields compared to the VPL. Consequently (t=50ms), the central part and the surround in the VPL is inhibited (blue points at t=50ms).

**Figure 9.**
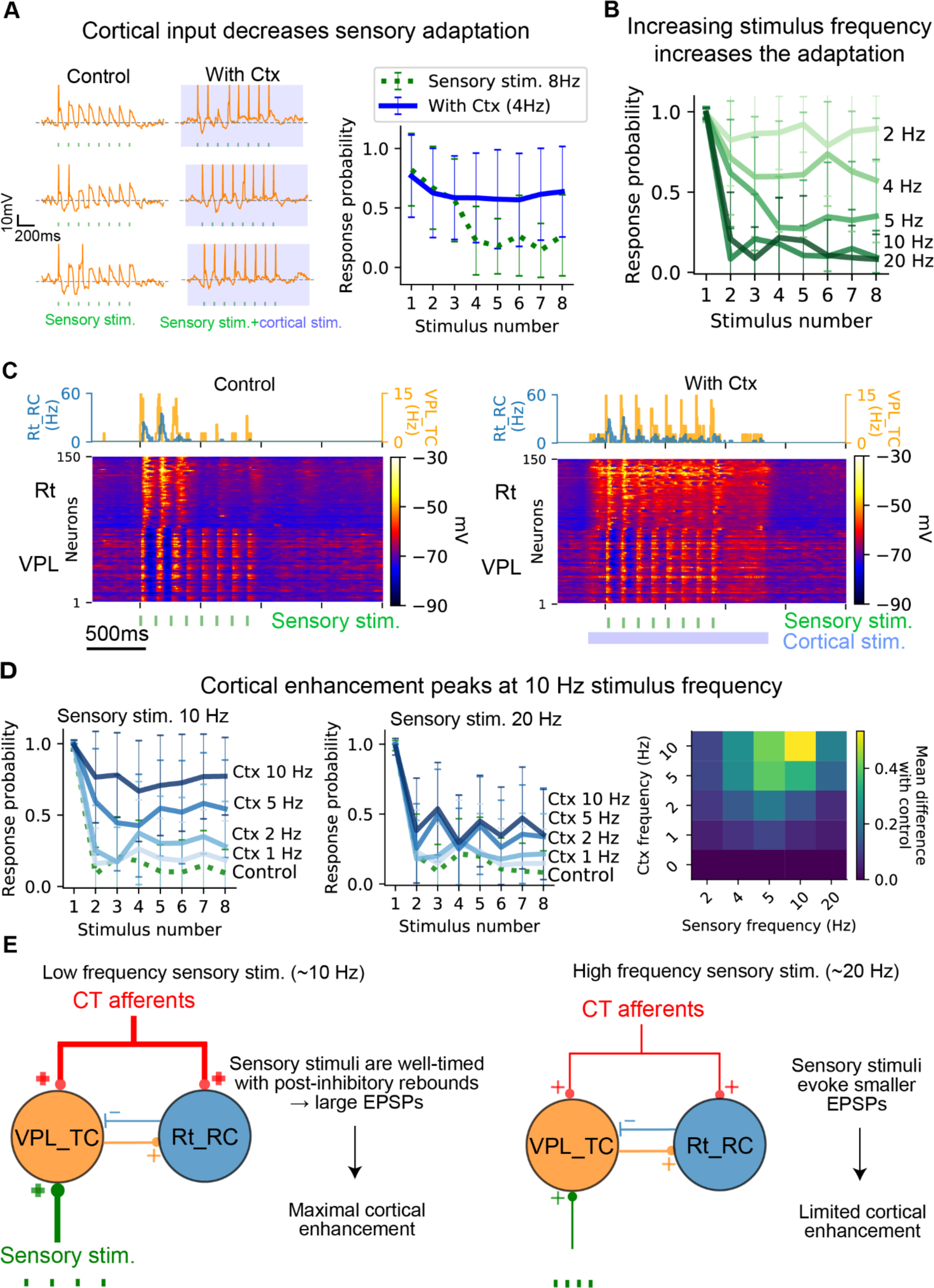
Frequency-dependent sensory adaptation and cortical enhancement of sensory responses Control condition (lightly-anesthetized *in vivo*-like state, see Methods) (cfr. Mease et al., 2014) (**A**) Left: example of a VPL_TC cell response (3 out of 25 repetitions) to a train of sensory stimuli (8 stimuli at 8 Hz, green). Note the adapting response: the cell responded only to the first stimuli in the train. Cortical activation at a mean firing rate of 4 Hz (blue rectangle) decreased the sensory adaptation by decreasing the distance from firing threshold. Right: comparison of VPL_TCs firing probability in response to the sensory stimulus and to the sensory stimulus with cortical activation (n=50 cells, 25 repetitions each). Note the increased firing probability with cortical activation (blue line). The markers show mean probability in response to each stimulus, vertical line the standard deviation. (**B**) The adaptation in the VPL to sensory responses increases with increasing frequency of sensory stimulus as shown by the response probability (each line represents the mean and error bars the standard deviation of 50 cells, 25 repetitions each). (**C**) Population voltage rasters in control condition and with cortical activation. Note that in control condition as well as with cortical activation the Rt responds to the first 2-3 stimuli in the train, with visible hyperpolarization in the VPL, which decreases from the 4^th^ stimulus. (**D**) Left: effect of different mean firing rate of cortical input to response probabilities for sensory stimuli at 10 and 20 Hz. Increasing cortical mean firing rates are represented by darker blue tones. Right: map showing the efficacy of cortical input in counterbalancing sensory adaptation for different sensory frequency and cortical mean firing rates. The efficacy was measured as the mean difference between the curves on the right. (**E**) Schematics explaining why cortical enhancement is greater for sensory stimuli ∼10 Hz than higher frequencies (∼20 Hz). Sensory stimuli around 10 Hz are well-timed with post-inhibitory rebounds and activation of low-threshold Ca^2+^ and cause larger EPSPs that can reach firing threshold with cortical activation. For higher stimulus frequency, EPSPs decrease in amplitude due to synaptic depression and cortical inputs are no longer sufficient to reach the firing threshold and counterbalance the adaptation (see also Fig. S2F-G).

#### 2.9.4 Simulating in vitro-like conditions

To simulate *in vitro-*like states, all neurons were left at their resting potentials (which ranged between −75 and −70 mV) and the only source of input was the spontaneous synaptic release from intrathalamic, medial lemniscus and corticothalamic synapses (at a rate of 0.01 Hz). The extracellular calcium concentration was set to 2 mM.

#### 2.9.5 Simulating depolarization levels

As a first approximation of the action of neuromodulators in the VPL and Rt, we applied constant current injections to the soma of each neuron. All neurons in the VPL or Rt regions were depolarized to the same target baseline membrane potential. The amplitude of the current was different for each neuron, to take into account the different input resistance of each morpho-electrical model.

#### 2.9.6 Simulation analysis

The spectrogram in Fig. 10D was calculated using the function *scipy.signal.spectrogram*, with inputs the sampling frequency of simulated membrane potential (10 kHz), *interval* = 5000, *overlap* = 0.99, and the other parameters with the default values.

**Figure 10.**
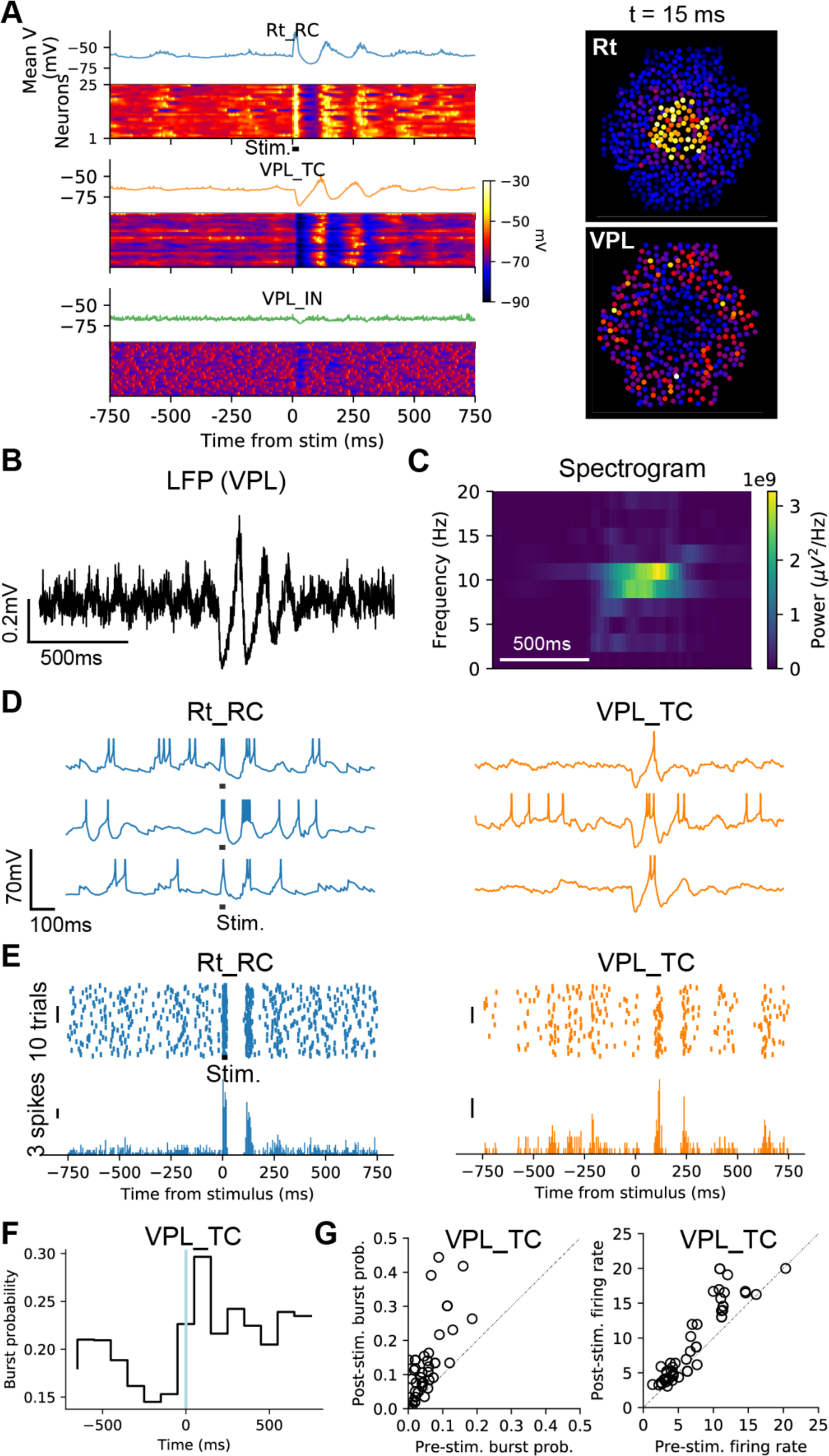
Activating the reticular nucleus increases thalamic bursts and initiates spindle-like oscillations (*in vivo* wakefulness-like condition). Spindle-like oscillations are evoked by localized pulse (20 ms) activation of 750 Rt_RC cells located at the center of the microcircuit. **(A**) Left: voltage rasters showing spindle-like activity. A sample of neighboring 25 neurons per each m-type is shown and color-coded according to their membrane potential. Right: topographical map of activity showing average membrane potential of Rt and VPL neurons in a 10 ms time window starting at the time indicated. (**B**) LFP recording from a central site in the VPL. Note the increased oscillatory activity after the stimulus applied to the Rt (not shown here). (**C**) Frequency-time analysis of the LFP in C showing increased power in the 8-10 Hz frequency range. (**D**) Example of single cell recordings for 3 cells per m-type. Note the burst responses in Rt_RCs and the IPSPs-rebound sequences in VPL_TCs. (**E**) Left: spike rasters and PSTHs showing the activity of one exemplar Rt_RC neuron (50 trials). Note the increased activity in response to the stimulus (black dot, 20 ms pulse) and a second peak, generated by network interactions. Right: Same as in A, for one example VPL_TC, note the post-inhibitory rebound response ∼100 ms after the Rt stimulation. (**F**) Histogram showing increased burst probability following the stimulus in VPL_TC (n=100 VPL_TC), as shown in experiments (Halassa et al., 2011). (**G**) Left: burst probability in VPL_TCs increases as a result of Rt_RC stimulus (each dot corresponds to one cell, the same sample as in F, n=100). Right: analysis of firing rates of VPL_TCs before and after the stimulus, as shown in experiments (Halassa et al., 2011). Pre-stim./post-stim. data were calculated in the 1s preceding/following the stimulus.

Burst probabilities (Fig. 10) were calculated as the ratio between the numbers of spikes belonging to a burst and the overall number of spikes. We considered a spike belonging to a burst when the interspike intervals were <= 15 ms and the first spike in the burst was preceded by a pause >= 50ms. For this analysis, we considered neurons that had a baseline activity between 1 and 20 Hz, as shown in corresponding publication (Halassa et al., 2011), Fig. S3.

To analyze the percentage of neurons firing for each m-type during each cycle of the oscillation (Fig. 12) we started by finding the oscillation peaks. The peaks were extracted from the firing rate histograms as input, using the *scipy.signal.find_peaks* function. We then added a peak corresponding to the time of the stimulus injected into Rt neurons (cycle 0). Spikes for each m-type were then assigned to the different cycles if they occurred within 30 ms of the oscillation peak.

**Figure 11.**
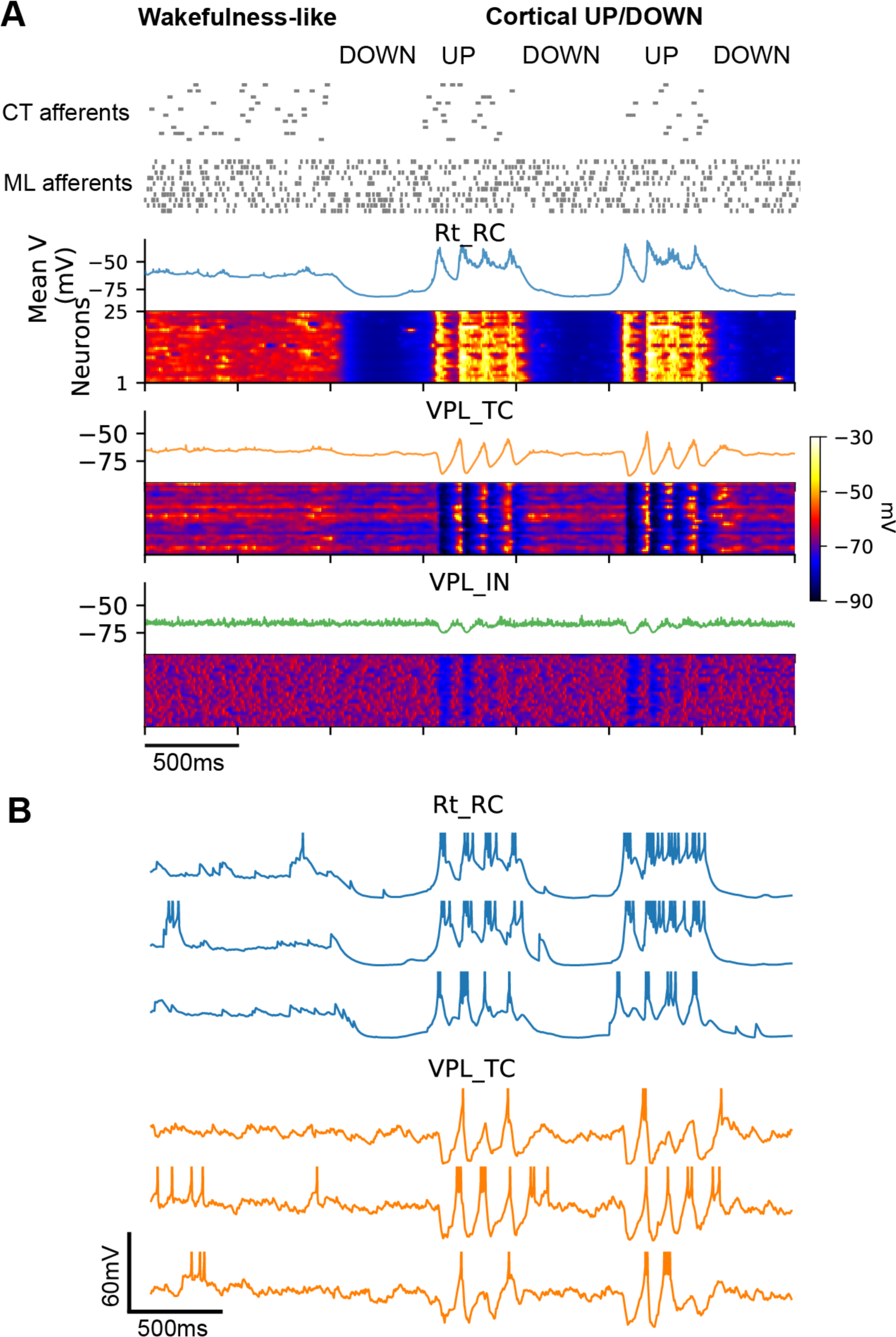
Spindle-like oscillations emerge in response to simulated cortical UP and DOWN states. The network is simulated in an *in vivo*-like condition (see Methods) for the first 1000ms. Afterwards, the background activity from CT afferents is removed (for 500ms), to approximate a cortical DOWN state and then re-activated (for 500 ms), to simulate an UP state. (**A**) Voltage rasters. A sample of 25 neurons per each m-type is shown and color-coded according to its membrane potential. The DOWN state results in a marked hyperpolarization in the Rt, while spindle-like oscillations emerge during the UP states. (**B**) Sample of single cell recordings from the neurons shown in A. Note the change in firing mode during the NREM-like phase, where Rt_RC and VPL_TC fire mainly low-threshold bursts. Spikes are truncated at −25 mV.

**Figure 12.**
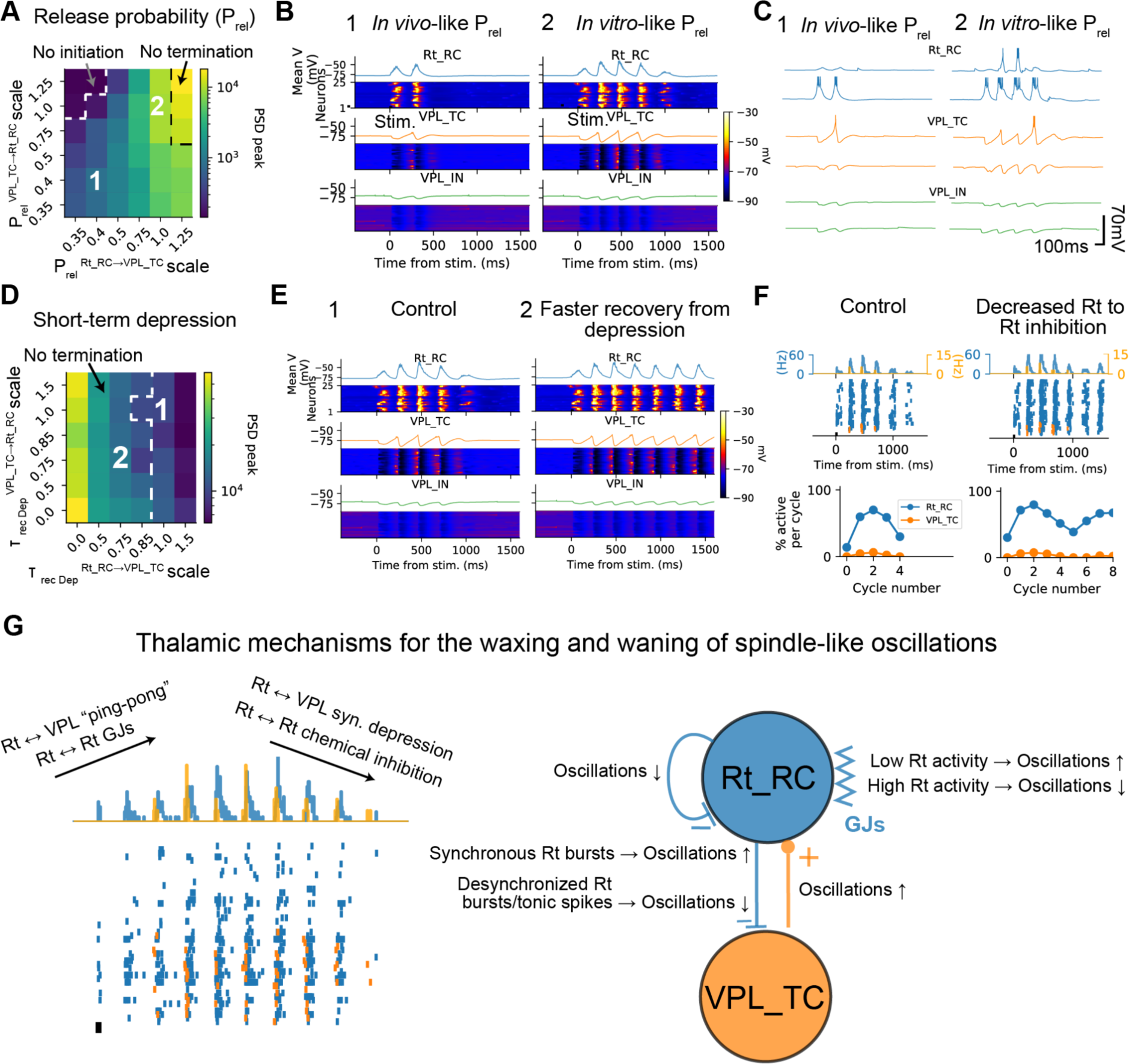
Waxing and waning of spindle-like oscillations emerges from intrinsic cellular and synaptic dynamics (*in vitro*-like condition) In this figure, the circuit is in an *in vitro*-like condition, thus membrane potentials are relatively hyperpolarized and synaptic interactions are stronger, as shown in A. A subset of 1000 Rt_RC cells located at the center of the microcircuit are stimulated with a 20 ms current pulse. (**A**) Parameter map showing the effect of synapse release probability between Rt_RC and VPL_TC cells on oscillation strength. Note that release probability in Rt_RC to VPL_TC connections is key for generating oscillations, which fails to terminate if its value is too high or cannot be generated if too low. (**B**) Right: voltage rasters showing spindle-like activity with *in vitro*-like synapse release probability, corresponding to parameter set 2 in panel A. A sample of 25 neurons per each m-type is shown and color-coded according to its membrane potential. This is the control condition used for this and the following figures. (**C**) Single cell recordings of a sample of 2 cells for each m-type (**D**) Parameter map showing the effect of short-term synaptic depression on evoked spindle-like oscillations. Smaller scale values mean faster recovery from short-term depression (τ=0 indicates no depression). Note that for faster recovery time constants in Rt_RC to VPL_TC synapses the oscillations fail to terminate (region of the parameter space at the left of the dashed line). (**E**) Voltage rasters showing oscillations in the control condition (1) and oscillations that fail to terminate (2), for the corresponding parameter sets shown in D. (**F**) The strength of inhibitory connections between Rt_RC cells plays a role in the termination, as shown by the PSTHs and spike rasters. The conductance of Rt_RC to Rt_RC synapses was reduced to 75% of the original value. This connection decreases the spiking activity in response to the stimulus (as shown by the cycle-by-cycle analysis of active Rt_RC cells, cycle number 0) and the minimum number of active cells to sustain the oscillatory ping-pong between Rt_RCs and VPL_TCs. (**G**) Left: schematic of the cellular and synaptic mechanisms underlying the waxing-and-waning of spindle-like oscillations observed in the model. The ping-pong of activity between the Rt and the VPL and GJs contribute to the waxing, while short-term synaptic depression (between Rt and VPL and vice-versa) and GABAergic inhibition between Rt cells contribute to the waning. Right: each connection can have a positive (↑) or negative (↓) effect on the oscillation or both, depending on network and single cell activity.

Oscillation strength (Fig. 12 and 14) was calculated as the maximal value of the power spectral density (PSD). The PSD was obtained using the function *scipy.signal.periodogram*.

**Figure 13.**
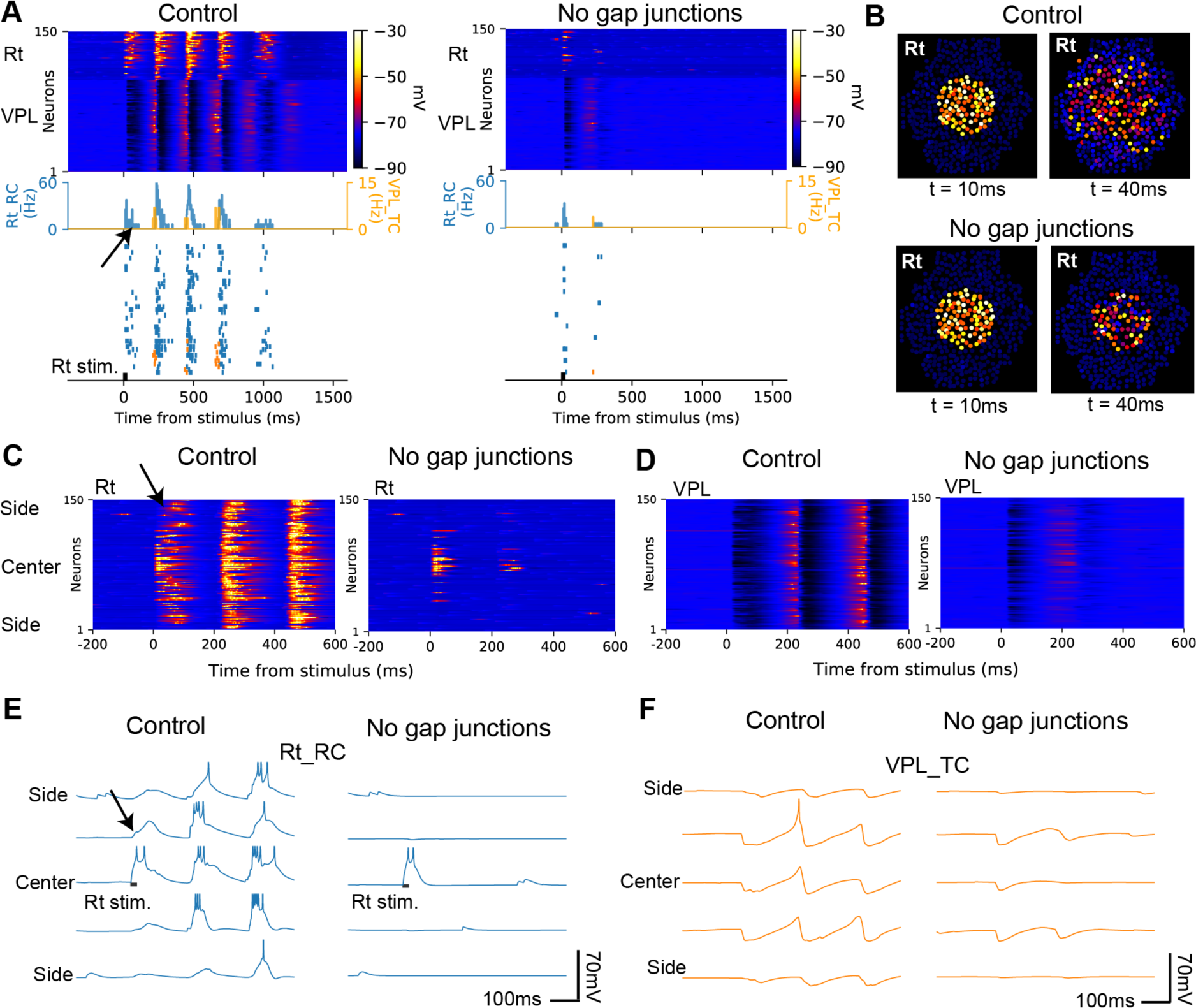
Gap junctions enhance spindle-like oscillations by spatial recruitment of low-threshold spikes in reticular cells. In this figure, the circuit is in an *in vitro*-like condition. A subset of 1000 Rt_RC cells located at the center of the microcircuit are stimulated with a 20 ms current pulse. (**A**) Spindle-like oscillation in control conditions (left) and with gap junctions (GJs) between Rt_RCs removed (right). Population responses are shown (from top to bottom) with voltage rasters, firing rate histograms and spike rasters. Note that when GJs are removed Rt_RCs fire only in a short time window after the stimulus (arrows). (**B**) Topographical activity maps in the Rt showing the membrane potential averages in 10 ms windows at 10 ms and 40 ms after the stimulus. Note that when GJs are removed, the activity at 40 ms is confined to the central part of the circuit. (**C**) Membrane potential along the lateral extent of the microcircuit for the simulations shown in A and B. Each row represents the membrane potentials of Rt and VPL neurons sorted by the distance from the center of the circuit. Note that with GJs, Rt_RC cells are recruited in a distance dependent-manner in the 40-100ms following the stimulus (arrow). (**D**) Same as in C, for VPL_TC cells. Note that without GJs the initial hyperpolarization is shorter in time and a smaller number of cells is hyperpolarized. (**E**) Single cell recordings of Rt_RC from the center and sides of the microcircuit are shown. Note that low-threshold spikes (arrows) are effectively transmitted through GJs and this is the mechanism contributing to the spatial recruitment of Rt_RCs. (**F**) Same as in E for VPL_TCs. Note that with GJs (control) the hyperpolarization is deep and slow enough for generating low threshold spikes.

**Figure 14.**
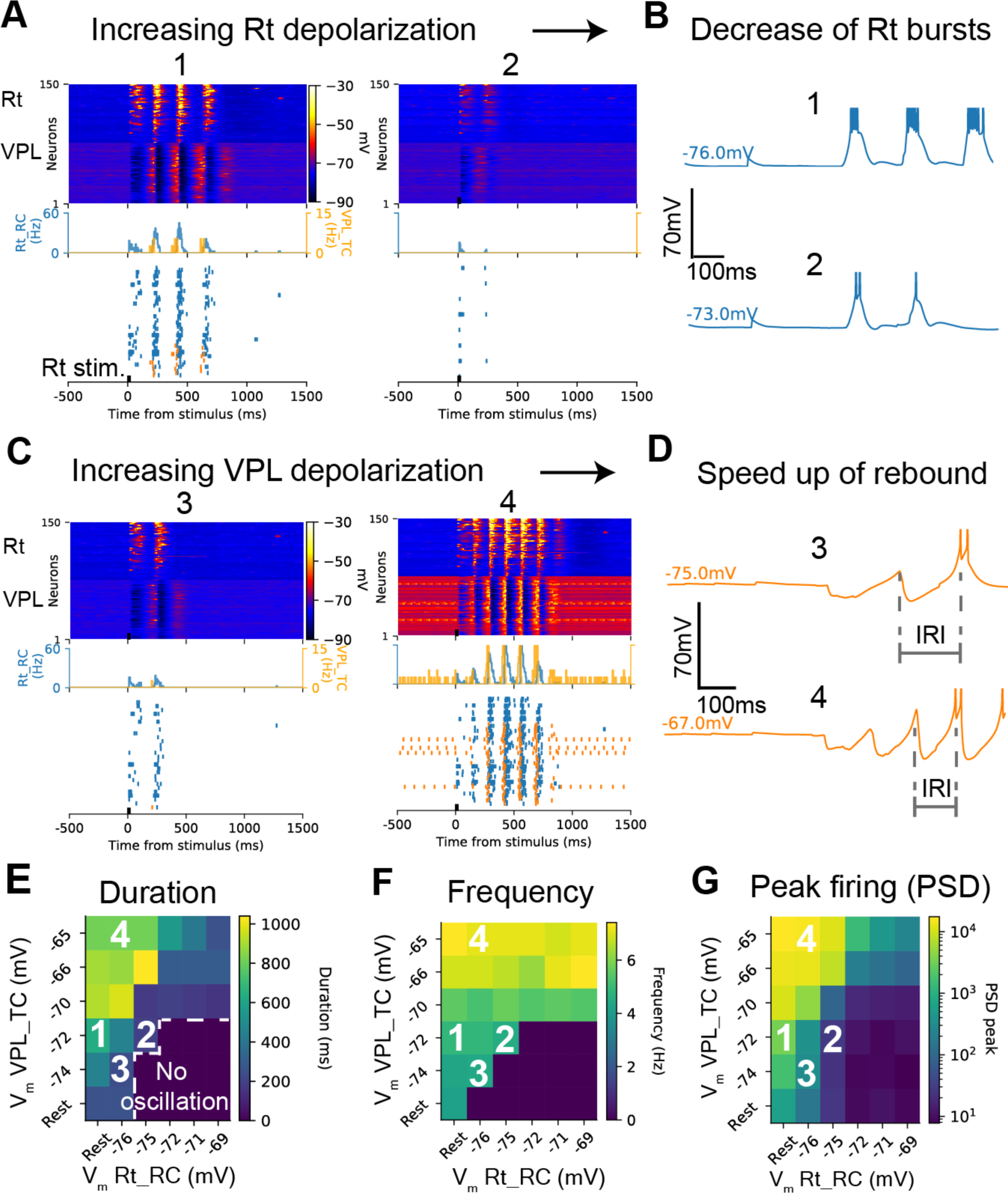
Depolarization levels modulate the properties of spindle-like oscillations. (**A**) Population voltage rasters, spike raster and firing rate histograms showing the effect of increasing Rt_RCs depolarization, while VPL_TCs are mildly depolarized. Note the decrease in oscillation duration with increasing Rt depolarization. (**B**) Single cell recordings of an exemplar Rt_RC showing that depolarizing the Rt resulted in fewer spikes per burst. (**C**) Same as in (A) for increasing depolarization of VPL_TCs. Note the increase of oscillation duration and frequency. (**D**) Same as in B, for an exemplar VPL_TC. Depolarization of the VPL resulted in higher probability of rebound responses (as a consequence of deeper IPSPs) and faster responses (shorter inter-rebound intervals, IRI), which resulted in increased oscillation frequency. (**E**) Parameter maps showing the effect of depolarizing the VPL and Rt on oscillation duration. The membrane potentials explored went from rest to close to firing threshold. Oscillation duration increased with VPL depolarization and decreased with Rt depolarization. (**F**) Same as in E, for oscillation frequency. Depolarizing the VPL has a stronger effect on the frequency than depolarizing the Rt. (**G**) Same as in E for the peak firing rate and its power spectrum density (PSD). Depolarizing the VPL increased the peak of the PSD, while depolarizing the Rt decreased it.

**Figure 15.**
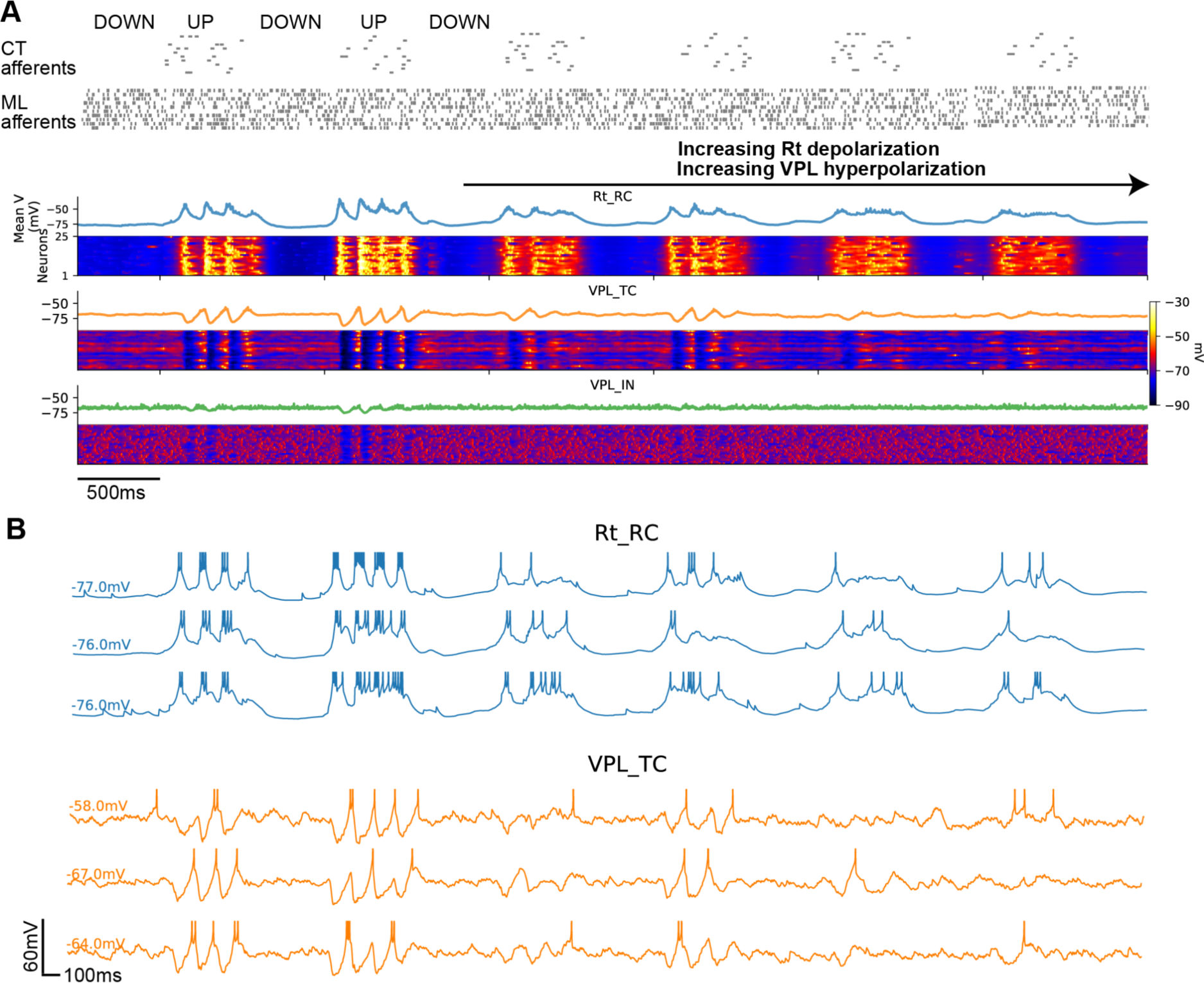
Spindle-like oscillations cease when simulating the effect of neuromodulation on thalamic and reticular neurons. (**A**) Simulated UP and DOWN states evoke reticular and thalamic depolarizations through afferent input, resulting in the “ping-pong” generation of spindle-like oscillations (as seen in Fig. 11). To approximate the differential effects of neuromodulators (e.g. acetylcholine) onto Rt_RC and VPL_TC we applied constant currents to depolarize Rt_RC and hyperpolarize VPL_TC cells. This resulted in spindle-like oscillations (left) being abolished (right). (**B**) Example single cell recordings from the simulation in A, note that while Rt_RC cells fire preferentially low threshold bursts during the cortical UP states (left), they transition to single spike modes when depolarized (right). The change in Rt_RC firing mode and hyperpolarization of VPL_TC cells resulted in a significant decrease of large amplitude IPSPs in VPL_TC cells and rebound bursts.

To calculate the oscillation duration (in ms) we used firing rate histograms for all the neurons and extracted their peaks, using the *scipy.signal.find_peaks* function (Fig. 14). Peaks were counted only if they were significantly higher than baseline firing rates. Oscillation duration was then calculated as the time difference between the last and the first peak.

To calculate oscillation frequency (Fig. 14), we computed the normalized autocorrelation of the firing rate histograms and extracted the time (oscillation period) corresponding to the first non-zero peak (Sohal and Huguenard, 2003). The inverse of the oscillation period corresponded to the oscillation frequency.

### 2.10 Thalamoreticular Microcircuit Portal and data integration in a FAIR knowledge graph

Experimental data, model entities and metadata are made available in the Thalamoreticular Microcircuit Portal (https://identifiers.org/bbkg:thalamus/studios/e9ceee28-b2c2-4c4d-bff9-d16f43c3eb0f). The portal includes data and model entities, including single neuron models, circuit files in SONATA format (Dai et al., 2020) and simulation output.

All data were integrated, and aligned to FAIR principles (Findible, Accessible, Interoperable, Reusable) using the Blue Brain Nexus software (RRID:SCR_022029). At the center of Blue Brain Nexus lies a knowledge graph which supports W3-standard “linked data” (https://www.w3.org/standards/semanticweb/data) storage and indexing. In the context of the knowledge graph, W3C (World Wide Web Consortium) Shapes Constraint Language (https://www.w3.org/TR/shacl/) was used to define FAIR data models (i.e. ‘shape’ the data and apply constraints). To support indexation of the datasets, specific data models were developed. Each individual data type was modeled according to a schema, available from Neuroshapes (http://neuroshapes.org/), to ensure standardization across different projects. This process made sure existing schemas, semantic markups, existing ontologies and controlled vocabularies were used.

For each data type, a minimum set of metadata was required to guarantee reusability of the data. We provide a concrete example of integration for an exemplar dataset consisting of neuron morphological reconstructions:

1. Identification of the dataset: 99 morphological reconstructions collected from acute brain slices through whole-cell patch clamp recording and biocytin filling, stored in .asc Neurolucida (rrid:SCR_001775) file format;
2. Identification of the metadata: A spreadsheet containing all the information related to the specimen, experimental protocol, date of the experiment, human agents involved in the experiment and reconstruction;
3. Creation of a data model: A schematic was developed according to the W3C PROV-O specification (https://www.w3.org/2011/prov/wiki/Diagrams), describing how the morphology was obtained in the form of a provenance graph;
4. Vocabulary and ontologies to integrate the dataset and metadata: Cell type terms from InterLex (rrid:SCR_016178) were used. InterLex is a dynamic lexicon of biomedical terms. For brain region, terms from the Allen Common Coordinate Framework version 3 were used (Wang et al., 2020; rrid:SCR_020999). To store species information, the NCBI organismal classification was used (Federhen, 2012). For storing information about strain, the Mouse Genomics Informatics database (http://www.informatics.jax.org/downloads/reports/MGI_Strain.rpt) was used. To store sex information, the <u>Phenotype And Trait Ontology</u> was used.

## 3 Results

### 3.1 Reconstructing thalamic and reticular morpho-electric neuron types

#### 3.1.1 Morphological types of thalamic and reticular neurons

To build a detailed model of thalamic microcircuitry, we started by collecting 157 three-dimensional reconstructions of neuronal morphologies of thalamocortical neurons, thalamic interneurons, and neurons of the reticular nucleus of the thalamus, from the mouse.

Neuron morphologies were reconstructed from both *in vitro* and *in vivo* labeling experiments in the mouse and included data from in-house experiments, and from open access datasets, such as the MouseLight Project at Janelia (Winnbust et al., 2019) (see Methods). For the purpose of this work, we classified the morphologies into three m-types (Fig. 2A). We grouped all thalamocortical (TC) neuron morphologies in the ventral posterolateral (VPL) nucleus microcircuit as one m-type, called VPL_TC, and all interneurons (IN) were grouped as VPL_IN. All neurons in the associated reticular nucleus (Rt) were grouped as a single m-type, Rt_RC.

We extended an existing algorithm to statistically recover dendrites from slicing artifacts (H. Anwar et al., 2009; Markram et al., 2015a) of *in vitro* labeled neurons and validated the results against *in vivo* labeled reconstructions. We then used a validated pipeline to generate a large dataset of thalamic and reticular morphologies (n = 92,970) that respected the biological variability (Markram et al., 2015), in a process called morphology diversification (see Methods).

#### 3.1.2 Electrical types of thalamic and reticular neurons

We characterized the firing behavior of over 100 thalamocortical neurons, interneurons and reticular neurons through patch-clamp recordings in brain slices of the mouse. TC and IN neurons were located mainly in the VPL and VPM nuclei of the thalamus and Rt neurons in the somatosensory sector of the Rt (Lam and Sherman, 2011; Pinault and Deschênes, 1998). VPL_TC and Rt_RC neurons were recorded with the injection of two different holding currents, to characterize their two firing modes: low-threshold bursting and tonic firing at hyperpolarized and depolarized membrane potentials, respectively.

We classified TC neurons as adapting (cAD_ltb) and non-adapting e-types (cNAD_ltb) by considering their tonic firing responses (Fig. 2B). These e-types were similar to those identified in rat TC cells (Iavarone et al., 2019). We found similar adapting and non-adapting responses in Rt_RCs. Other firing patterns have been described for reticular neurons, when considering low-threshold firing responses (Clemente-Perez et al., 2017; S. E. Lee et al., 2014; Lee et al., 2007). In our dataset, we mainly observed neurons with an intermediate burst propensity, which fired typically 1-2 bursts, followed by tonic firing for higher stimulus amplitude, similar to the “typical burst” type, which is more common in somatosensory sectors of the Rt (Lee et al., 2007). With our standard set of stimuli protocols (see Methods), interneurons mainly exhibited an accommodating firing behavior. Since they tended to initially fire action potentials with short interspike intervals, we classified all IN recordings as one e-type, called burst accommodating (bAC). When deeply hyperpolarized, some IN responded with rebound bursts and, in rare cases, they generated spontaneous oscillations (Simko and Markram, 2021).

#### 3.1.3 Morpho-electrical models of thalamic and reticular neurons

We previously demonstrated that a multi-objective optimization pipeline can be applied to capture different firing modes of thalamocortical neurons (Iavarone et al., 2019a). We applied a similar strategy to build electrical models (e-models) for the e-types shown in Fig. 2B (see Methods). In brief, we used a multi-objective optimization algorithm (Druckmann et al., 2007; Hay et al., 2011b; Iavarone et al., 2019a) with a set of electrical features (e.g. spike amplitude, firing frequency, afterhyperpolarization depth) extracted from the experimental recordings as parameters for the objective functions, in combination with an exemplar morphology, to optimize for ion channel densities (or peak conductances). All model neurons included models of transient sodium, persistent sodium, A-type transient potassium, delayed potassium, low-threshold calcium, high-threshold calcium, calcium-activated potassium (SK-type), h-current ion channel currents. All cellular compartments (i.e. somata, dendrites and axon initial segments) contained active membrane mechanisms. Rt_RC e-models followed the same approach as for VPL_TC cells, as previously published (Iavarone et al., 2019a), with the additional electrical features to quantify the deeper post-burst afterhyperpolarization observed in Rt_RCs (Avanzini et al., 1989; Cueni et al., 2008). To validate and test the generalization of the neuron models, we used features from current stimuli not used during the optimization phase (see Methods for detail).

In this way, we created five e-models, one for each e-type, and combined them with the 92,970 morphologies generated during the morphology diversification step to generate 142,678 unique morpho-electrical models. We assessed the quality of each morphology-electrical model combination (me-model) using a battery of test stimuli and rejected those having electrical features and firing behavior significantly different from the experimental data (see Methods).

### 3.2 Reconstructing thalamoreticular microcircuit structure

To establish the structure of the thalamoreticular microcircuit, we acquired neuron density data, defined the microcircuit volumetric dimensions, reconstructed the neuron numbers and composition, and established the soma positions and morphology alignment in the volume.

#### 3.2.1 Measured neuron densities

To obtain neuron counts, we used semi-automated cell counting in consecutive sagittal slices of a mouse brain and divided those numbers by the calculated volume (see Methods). We found an average cell density of 68,750 ± 1,976 cells/mm^3^ for Rt and 57,467 ± 5,201 cells/mm^3^ for the VPL (mean±std, n=37 slices).

#### 3.2.2 Defining microcircuit dimensions

To define the dimensions of a thalamic microcircuit, we started by identifying a subvolume spanning a portion of the ventral posterolateral nucleus of the thalamus (VPL) and the reticular nucleus of the thalamus (Rt). Its longest (vertical) dimension encompasses the two nuclei, with the Rt as the topmost region (Fig. 3). We chose the VPL nucleus because it receives information from the hindlimb (Joseph T Francis et al., 2008) and relays it to the somatosensory cortex; the corresponding cortical microcircuit was reconstructed in a previous model (Markram et al., 2015a). The Rt is intimately associated with the different thalamic nuclei and it is the primary source of inhibition to the thalamus (Cavdar et al., 2013; de Biasi et al., 1986; Houser et al., 1980; Pinault, 2004). Since the VPL does not have a clear modular organization, we followed a previously published approach to define the microcircuit radius and its height (Markram et al., 2015a).

To constrain the dimensions, we used the 3D reconstructed morphologies of the Rt_RCs to calculate the horizontal dimension of the microcircuit and the vertical dimension of VPL and Rt from the Allen Brain Atlas for its height (Goldowitz, 2010). The dendrites of Rt_RCs were used as they have the largest reach along the horizontal dimension (see Methods).

The estimated dimensions correspond to a microcircuit with a volume of ∼0.22 mm^3^, a base of 323 μm in length, and a total height of 800 μm (250 μm corresponds to the Rt and 550 μm to the VPL, see Methods for details).

#### 3.2.3 Reconstructing neuron numbers and composition

Having established the model microcircuit volume, we distributed the neurons according to densities and excitatory/inhibitory ratios measured experimentally (Fig. 3).

We next collected information on the inhibitory and excitatory neuron ratios in the microcircuit regions (Fig. 3.2C). As reported in many studies (de Biasi et al., 1986; Houser et al., 1980; Pinault, 2004), 100% of neurons in Rt were inhibitory. In this first draft, we included 0.5% of inhibitory cells in the VPL, according to the estimates in the Blue Brain Cell Atlas (RRID:SCR_019266), an open-access cell atlas for the mouse brain (Erö et al., 2018). This atlas is a resource that integrates whole brain Nissl and gene expression stains to predict neuron densities and positions.

As a result, the microcircuit was populated with 4,909 ± 283 Rt_RCs, 8,952 ± 517 VPL_TCs and 47 ± 2 VPL_INs, for a total of ∼14,000 neurons. The numbers correspond to the mean and standard deviation of the different microcircuit instances (see Methods).

Once we had obtained the inhibitory and excitatory cells densities, the morphological and electrical types composition (me-composition) was directly determined (Fig. 3.2D): the reticular nucleus (Rt) had 100% of neurons of m-type Rt_RC. Each Rt_RC neuron had one of two e-types (57% cAD_ltb and 43% cNAD_ltb). In the VPL, all inhibitory neurons corresponded to the VPL_IN m-type and bAC e-type. The excitatory neurons in VPL, which corresponded to thalamocortical cells (m-type VPL_TC), had e-type cAD_ltb (64%) or cNAD_ltb (36%). The percentage of e-types for each m-type (e-type fractions) were derived from our *in vitro* electrophysiological recordings.

#### 3.2.4 Soma positions and morphology placement

After establishing the dimensions and the number of neurons for each me-type, we generated somata positions using an algorithm to fill the space, ensuring that somata did not overlap and that neurons were uniformly spaced between each other (Fig. 3E,F). Once neuron positions were defined, we used a second algorithm to select the morphology that best fulfilled the anatomical constraints of the microcircuit (see Methods). The logic followed was based on experimental findings showing that the axons of reticular neurons project towards the different thalamic nuclei of the thalamus (Pinault and Deschênes, 1998) and that thalamocortical cells have axonal collaterals in the reticular nucleus (Harris, 1987; Monconduit et al., 2006). To choose the best morphology for each position, we manually annotated these patterns on each reconstructed morphology and calculated their overlap with the microcircuit subregions (see Methods).

### 3.3 Reconstructing and validating synaptic connectivity

The detailed connectivity between individual neurons in the microcircuit was built by adapting an existing algorithm (Markram et al., 2015a; Reimann et al., 2015). Detailed anatomical studies found a linear relation between the available dendritic surface in the thalamus and the bouton numbers on reticular axons (Pinault and Deschênes, 1998). This finding suggested that potential synaptic locations in the thalamus between Rt and TC neurons can be predicted by the statistical overlap between neurites (S. L. Hill et al., 2012).

To establish synaptic connectivity (Fig. 4), we used neuron morphologies placed in the microcircuit and the available data on axonal bouton densities from 3D reconstructed morphologies (number of boutons/axonal length) as constraints. We included synapses from ML and CT afferents by using volumetric bouton densities as constraints (number of boutons/tissue volume).

We found that the model reproduced findings from electron microscope studies that had not been used as constraints. We compared the synapse convergence onto reticular neurons (Liu and Jones, 1999) and the distribution of number of synapses per connections (Morgan and Lichtman, 2020)) and both gave results comparable to the experimental counterpart (Fig. 4D).

#### 3.3.1 Reconstructing intrathalamic connectivity

Potential synaptic locations were identified by computing all appositions between the neuron morphologies, i.e. potential synapses, filtering using additional experimental constraints (see Methods for detail). Along with the physical surface area available on the morphologies, we used as a constraint the bouton densities (number of boutons / axonal length) on the axons of Rt_RCs and VPL_TCs (Fig. 4A). In our experimental dataset, we found that TCs had on average 0.102 ± 0.021 boutons / μm (n = 9 axons) and Rt neurons 0.124 ± 0.002 boutons / μm (n = 2 axons).

Synaptic connections in the model were formed between all m-types, except for VPL_TCs to VPL_TCs and from VPL_TCs to VPL_INs. Connections between VPL_TCs are likely to disappear during development (Lee et al., 2010), while we did not find any experimental evidence of connections between TCs and INs, neither in the ventrobasal thalamus, nor in the visual thalamus, where INs are present in higher proportions (Arcelli et al., 1997; Evangelio et al., 2018; Jager et al., 2021). Connections between all m-types were formed by presynaptic axons and postsynaptic dendrites and somata, while those formed by INs were largely established by presynaptic dendrites (Fig. 4B), as shown in the visual thalamus (Morgan and Lichtman, 2020; Sherman, 2004).

An important difference between the reconstruction of connectivity in the cortical microcircuit (Reimann et al., 2015; Markram et al., 2015) and the present model is that we did not explicitly remove connections between neurons if they shared only one contact, as we did not have any evidence of this constraint. Rather, a recent electron microscope reconstruction of an IN in the visual thalamus, showed that most connections from INs involve only one functional synapse (Morgan and Lichtman, 2020).

The resulting distributions of the number of functional synapses per connection, i.e. the number of functional contacts between a pair of neurons, followed geometric distributions, similar to the one shown in Fig. 4D1. Most of the m-type to m-type connections had 60-70% of pairs with one synapse, with the exception of Rt_RCs to VPL_TCs, where most of the connections had 2 (30%) or more synapses (70%).

#### 3.3.2 Reconstructing connectivity from lemniscal and corticothalamic afferents

We included synapses from extrathalamic sources, i.e. from the sensory periphery through the medial lemniscus (ML) and cortex (corticothalamic afferents, CT). We used as experimental constraints the volumetric bouton densities of lemniscal synapses in the mouse VPM (Takeuchi et al., 2017), since data for the VPL was not available (Fig. 4C). For CT afferents, we used the relative proportions of corticothalamic to lemniscal synapses onto TCs in the VB (around 12) and the ratio of CT to TCs synapses in the reticular nucleus (around 2.8) (Çavdar et al., 2011; Mineff and Weinberg, 2000). Lemniscal synapses were assigned to 2,601 virtual fibers (see Methods for details); this number was estimated by taking into account the number of TCs in the model, the number of VPL neurons and number of dorsal column nuclei (DCN) projecting to the thalamus (Shishido and Toda, 2017). We used a mouse cell atlas (Erö et al., 2018) to determine the number of excitatory neurons in the VPL and DCN. The number of CT fibers was 75,325, consistent with data reporting a ratio of ∼10 between CT afferents and the corresponding TC neurons (Crandall et al., 2015; Monconduit et al., 2006; Sherman and Koch, 1986).

#### 3.3.3 Predicted synapse numbers, afferent, and efferent neuron numbers

This microcircuit model is a tool to predict the convergence and divergence of the different thalamic m-types. We found that each neuron in the microcircuit projected on average to 246 ± 88 other neurons (mean ± std, sample of 1,000 neurons); each Rt_RC neuron projected to 64 ± 28 Rt_RCs and 136 ± 60 VPL_TCs; each VPL_TC projected to 34 ± 46 Rt_RC neurons; VPL_IN sent efferents on average to 220 VPL_TCs (± 78.0). The total number of intra-thalamic synapses in a microcircuit was 4.77 million. Afferent lemniscal fibers made a total of 17,998 synapses in the VPL portion of the microcircuit, while synapses from corticothalamic fibers were 40,905 (sum for VPL and Rt in the microcircuit). In a mesocircuit, comprised of a central microcircuit and surrounded by six others (see Fig. 6A), we found a total of 44.2 million synapses. Each neuron received inputs on average from 203 ± 41.0 other neurons. Each Rt_RC neuron receives inputs from 28 ± 15 other Rt_RCs and 74 ± 28 VPL_TCs, while VPL_TCs are contacted on average by 75 ± 32.0 Rt_RCs.

In a preliminary investigation, we determined the relative proportions of closed and open-loop configurations between Rt_RCs and VPL_TCs. We found that closed-loops were present in our model and that they were a minority of the connections, in qualitative agreement with experimental findings (Gentet and Ulrich, 2003; Pinault and Deschênes, 1998; Shosaku et al., 1989). As a starting point for this analysis, we considered all connected VPL_TC to Rt_RC (or Rt_RC to VPL_TC) pairs and for each presynaptic neuron we counted how many among its postsynaptic neurons it received input from. In this way we found that the percentage of closed-loops was always lower than 10%.

#### 3.3.4 Reconstructing and validating synapse physiology

After establishing the anatomically constrained connectivity, we modeled synapse physiology with available data from in-house experiments and the literature on short-term plasticity, post-synaptic potential amplitudes (PSPs), time constant of synaptic currents and reversal potentials (Fig. 5).

We identified three short-term synaptic plasticity types: inhibitory depressing (I2), excitatory depressing (E2) and excitatory facilitating (E1) (Ecker et al., 2020; Markram et al., 2015a). We used synapse models featuring stochastic transmission and short-term plasticity, (Fuhrmann et al., 2002; Markram et al., 2015a) and constrained the parameters of the Tsodyks-Markram model of short-term plasticity with available thalamic data (see Table 1 and Fig. 5A-B). In this way, we provide a first comprehensive map of synapse types in a thalamic microcircuit with the main external afferents (Fig. 5B). For instance, we took into account that synapses between VPL_TC and Rt_RC depress more quickly than synapses between Rt_RC and VPL_TCs are more rapidly depressing than those between VPL_TCs and Rt_RCs (Cox et al., 1997; Gentet and Ulrich, 2003) (Fig. 5D). Connections from presynaptic interneurons have never been characterized in the somatosensory thalamus of the rodent and we found that they were depressing as well (Simko and Markram, 2021). Interestingly, synapses from corticothalamic afferents were described as being facilitating in first-order thalamic nuclei and in the reticular nucleus (Connelly et al., 2016; Crandall et al., 2015; Cruikshank et al., 2010; Jurgens et al., 2012; Landisman and Connors, 2007; Miyata, 2007). For some connections, such as extrinsic synapses from CT fibers, we found information in the literature regarding their short-term plasticity types, but experiments were limited to the analysis of the first 2-4 consecutive EPSPs. In those cases, parameters were predicted from similar pathways from our recordings or from the neocortical microcircuit model (see Methods for details, Fig. 5B).

Similarly, we used available paired-recording data and generalization principles (see Methods) to assign synaptic conductance values (*g_syn_*) to match experimentally recorded PSP amplitudes. We predicted that single *g_syn_* from inhibitory neurons are in general small (e.g. 0.9 ± 0.23 nS for VPL_IN to VPL_IN), while conductances from VPL_TCs and lemniscal afferents are larger (>2 nS), consistent with being “driver” synapses (Mo et al., 2017; Sherman and Guillery, 1998) (Mo et al., 2017; Sherman and Guillery, 1998), while corticothalamic synapses have small *g_syn_* (<0.5 nS), but are facilitating (Fig. 5D).

To model synapse kinetics, we used existing models of synaptic currents (Markram et al., 2015) and included literature findings on decay time constants, reversal potentials and the relative contribution of AMPA, NMDA, GABA_A_ and GABA_B_ currents, summarized in Table 1 (Arsenault and Zhang, 2006; Deleuze and Huguenard, 2016; Miyata and Imoto, 2006; Warren et al., 1994; Zhu et al., 1999). In the case of inhibitory synapses, we included only GABA_A_ currents in this first draft, since our existing GABA model does not take into account the non-linear dependence of GABA_B_ activation on presynaptic activity (Destexhe and Sejnowski, 1995; Kim et al., 1997; Wang et al., 1995).

We found that the model was able to reproduce data that were not used so far: the stochastic nature of synaptic release was assessed against the coefficient of variation of the first PSP amplitudes (Table 2 and Fig. 5C1). Moreover, when we performed *in silico* paired-recording on neuron pairs that were not considered when assigning the *g_syn_* values, we found that they reproduced very closely the experimental counterpart (Table 3 and Fig. 5C2). This validation was particularly important, since PSP values emerge not only from the specific *g_syn_* values, but also from the dendritic properties of the single neuron models, the distance of the synapses from the soma and the initial synapse release probability.

**Table 3.**
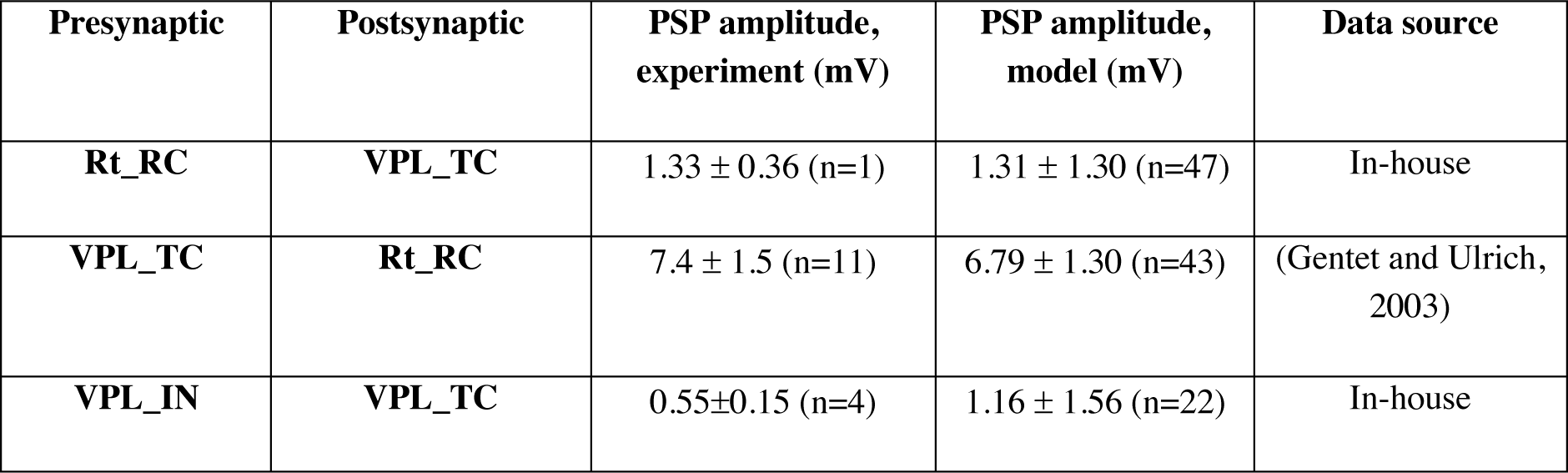
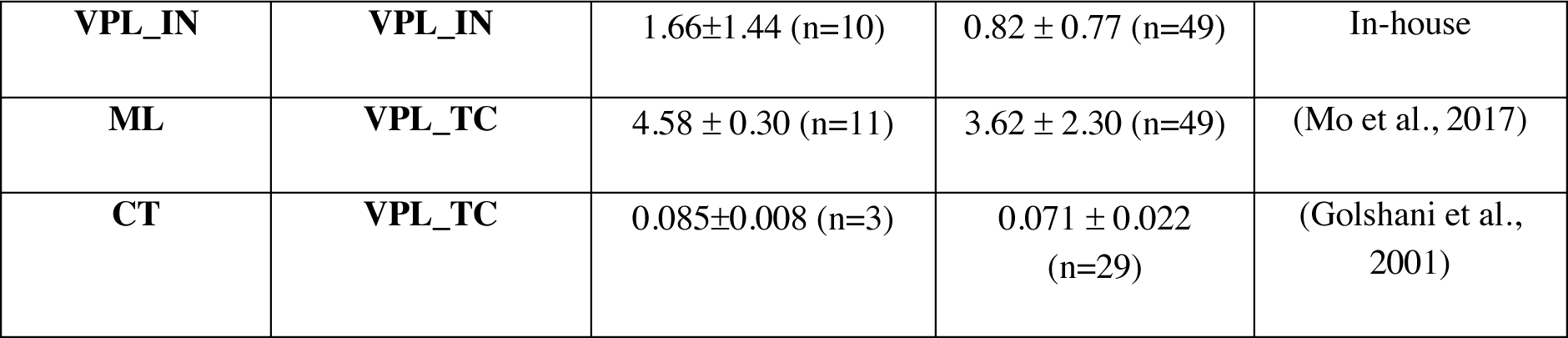
Postsynaptic potential (PSP) amplitudes. PSP amplitude values as characterized experimentally through *in vitro* paired recordings. Values are reported as mean ± standard deviation (of multiple pairs). (Related to Fig. 5C2).

| Presynaptic | Postsynaptic | PSP amplitude, experiment (mV) | PSP amplitude, model (mV) | Data source |
| --- | --- | --- | --- | --- |
| **Rt_RC** | **VPL_TC** | 1.33 ± 0.36 (n=1) | 1.31 ± 1.30 (n=47) | In-house |
| **VPL_TC** | **Rt_RC** | 7.4 ± 1.5 (n=11) | 6.79 ± 1.30 (n=43) | (Gentet and Ulrich, 2003) |
| **VPL_IN** | **VPL_TC** | 0.55±0.15 (n=4) | 1.16 ± 1.56 (n=22) | In-house |
| VPL_IN | VPL_IN | 1.66±1.44 (n=10) | 0.82 ± 0.77 (n=49) | In-house |
| ML | VPL_TC | 4.58 ± 0.30 (n=11) | 3.62 ± 2.30 (n=49) | (Mo et al., 2017) |
| CT | VPL_TC | 0.085±0.008 (n=3) | 0.071 ± 0.022<br>(n=29) | (Golshani et al., 2001) |

### 3.4 Reconstructing and validating gap junction connectivity

Neurons in the reticular nucleus of the thalamus are functionally connected through electrical synapses, providing a basis for neuronal synchronization in thalamic networks (Haas et al., 2011; Landisman et al., 2002; S.-C. Lee et al., 2014; Long et al., 2004). Different studies in rodents estimated that gap junctions-coupled neurons represent between 30-50% of the total neuron population in the Rt (Crabtree, 2018; Deleuze and Huguenard, 2006; Lam et al., 2006; S.-C. Lee et al., 2014). Along with intra-thalamic GABAergic synapses, they play a crucial role in the balance between synchronization and desynchronization in thalamic networks and rhythm generation (Beenhakker and Huguenard, 2009).

Our computational model provides a basis to study the functional impact of gap junction connectivity on network dynamics and rhythm generation, overcoming important limitations of experiments in the slice. For instance, difficulties in detecting the activity of gap junctions that are remote from the recording site in the soma or loss of dendritic mass due to slicing artifacts (Long et al., 2004).

#### 3.4.1 Gap junction connectivity between reticular neurons is predicted by dendrodendritic appositions

We identified gap junction locations (GJs) by following a similar approach as was used to identify synapse locations. We started by finding all possible appositions between Rt_RCs dendrites and somata. By doing so, the resulting neuron divergence (number of first-order postsynaptic neurons) was significantly higher than values reported in the literature (Lee et al., 2014) (Fig. 6A1). Moreover, the mean number of GJs per neuron was higher than values reported in other species or brain regions of ∼300 per neuron (Amsalem et al., 2016). We therefore removed a random fraction of appositions until we matched the experimental neuron divergence (Fig. 6A2). We found that the model best matched the experimental data when retaining 30% of the potential dendrodendritic appositions.

We then validated the resulting connectivity by comparing the extent of anatomical coupling in the model with available dye-coupling data in the mouse Rt (Lee et al., 2014). Consistent with the experiment findings, we find that, in the model, each neuron is directly coupled with 2-20 other neurons and that the majority of coupled neurons was at 50-100 μm from the primary injected neuron (Fig 6B). When analyzing single model neurons, most of the coupled neurons were located at 40-120 μm, as found experimentally (Fig. 6B). Rarely, coupled neurons can be found at distances of 300-400 μm, consistent with the extent of some Rt_RC neuron morphologies and the experimental findings (Fig. 6B). These results indicate that many aspects of GJ connectivity, in particular its distance-dependence, can be predicted by the morphological properties of Rt_RC dendrites.

We assumed a gap junction conductance of 0.2 nS based on prior work (Amsalem et al., 2016). To validate the coupling strength, we computed the coupling coefficients (CCs) between pairs of neurons with simulated paired recordings (Fig. 6C). In the model, we found that electrically coupled Rt_RC neurons had CCs values of 0.023 ± 0.008 (mean ± standard deviation, Fig. 6C) and that the mean value fell within the reported variability in mouse Rt neurons (Landisman et al., 2002).

### 3.5 Spontaneous and evoked activity in the model thalamoreticular microcircuit

We first explored spontaneous and evoked activity in the microcircuit in a simulated *in vivo* wakefulness-like condition (Fig. 7, see Methods). In this condition, the model reproduces the distribution of firing rates in first-order thalamic and reticular nucleus neurons during quiet wakefulness, characterized by low firing rates (<10 Hz) (Born et al., 2021; Hartings et al., 2000; Nestvogel and McCormick, 2021). The simulation exhibits uncorrelated firing activity in all m-types, with higher firing rates in VPL_INs due to their lower spiking threshold in the model (Fig. 7A and Methods). Single cell activities are dominated by single tonic spikes (Fig. 7B), consistent with their predominance over low-threshold bursts in wakefulness-like states (Urbain et al., 2015).

We then compared spontaneous activity with sensory evoked responses with brief activation of a subset of ML fibers, simulating a whisker flick or a brief electrical stimulation to the hindlimb (Kimura, 2017). We found increased population firing rates in VPL_TC neurons, with a peak in firing in a short time window (5-10 ms) following the stimulus (Fig. 7 and <u>Video</u> <u>1</u>). This activity is followed by a period of silence in some cells, lasting 100-200 ms Fig. 7B). Rt_RCs exhibit an increase in firing rate compared to their spontaneous activity as a result of the excitation they receive from VPL_TCs (Fig. 7B). The increased activity in Rt_RCs lasts for 50-100 ms after the stimulus and gradually decreases to baseline levels, resulting in a clear hyperpolarization of VPL_TCs (Fig. 7A). The longer activation of Rt_RCs is in line with experimental findings and suggests an important role of the Rt in limiting sensory responses and focusing them on rapid perturbations (Hartings et al., 2000). Single neuron membrane potentials of VPL cells show heterogeneous responses (spikes, associated or not with IPSPs from the Rt, IPSPs only) (Fig. 7C).

#### 3.5.1 Increasing sensory stimulus size increases surround inhibition in the thalamus

Receptive fields in the VPL vary in size and are more focal in subregions representing individual digits and broader for regions representing the body or limbs (Joseph T. Francis et al., 2008). Inhibition from the Rt comprises an important contribution in shaping the temporal and spatial properties of thalamic receptive fields (Born et al., 2021; Lee et al., 1994; Shosaku, 1986; Shosaku et al., 1989; Soto-Sánchez et al., 2017). Therefore, we explored the effect of inhibitory feedback from the Rt on network responses by simulating sensory stimuli of increasing size.

We simulated the activation of an increasing number of afferent ML fibers, and identified a threshold for minimal activation (increased population responses) of both Rt and VPL of 60 ML fibers. Further increasing the number of afferent fibers, increased the population responses in the Rt and VPL, with marked Rt-mediated hyperpolarization when 160-260 fibers were activated (Fig. 8A). Both response latency and its variability decreased with increasing stimulus size, indicating increasingly synchronous responses (Fig. 8B). Interestingly, population firing rates in the Rt increased more than in the VPL when more than 260 ML fibers were activated (Fig. 8A), confirming the high responsiveness of reticular neurons to sensory stimuli (Hartings et al., 2000). The model shows that not only cells directly activated by the lemniscal input exhibit stimulus size-dependent responses, but also the indirect inhibition from Rt_RC cells is stimulus size-dependent: at increasing stimulus size more Rt_RCs are recruited and cause greater hyperpolarisation in the VPL (Fig. 8A, C).

We then investigated the impact of the recruitment of VPL and Rt neurons at the topographical level (Fig. 8D and <u>Video 1</u>). Central VPL cells were depolarized or spiked, while the activation in the Rt was broader, with some degree of VPL-mediated depolarization in the periphery of the Rt. This result suggests that receptive fields in the Rt are somewhat larger than in the VPL, yet they provide topographically aligned inhibition to the thalamus in a focal manner, as shown for visual reticular neurons (Born et al., 2021; Soto-Sanchez et al., 2017). The activity in the Rt results in broad inhibition of the VPL, extending beyond the area directly activated by the stimulus (surround inhibition). This result unveils an important role of the Rt in the control of receptive field size, which does not strictly require extra-thalamic inputs, such as cortical feedback. The Rt contributes to precise responses in the VPL not only by rapidly inhibiting directly responding neurons, but also by limiting the response in the surrounding area.

#### 3.5.2 Thalamic responses to sensory input exhibit adaptation and cortical enhancement

Numerous studies have shown adaptation to sensory stimuli in the lemniscal pathway due to short-term depression of lemniscal EPSPs (Manuel A Castro-Alamancos, 2002; Manuel A. Castro-Alamancos, 2002; Diamond et al., 1992; Martin-Cortecero and Nuñez, 2014; Simons and Carvell, 1989). In the model, we were able to reproduce sensory adaptation at different frequencies and a recent study showing that depressed responses in the somatosensory thalamus can be enhanced by cortical activation in anesthetized mice (Mease et al., 2014).

We studied the model responses to repetitive sensory stimuli and with activation of the corticothalamic afferents before and during the sensory stimuli (Mease et al., 2014). Activating the lemniscal fibers with trains of stimuli at 8 Hz, results in high response probability to the first stimuli, while subsequent EPSPs exhibit decreased amplitudes (Fig. 9A). The activation of cortical afferents, 200 ms before the sensory stimulus with noisy input at a mean firing rate of 4 Hz, increased the firing probability to all the other stimuli in the train, thus counterbalancing the adaptation, at least in part (Fig. 9B), as shown in the corresponding experiment (Mease et al., 2014).

With ML stimulation only, we also found that the adaptation was minimal at 2 Hz (firing probability ∼0.8 for all the stimuli), but was already evident at 4 Hz (firing probability of ∼0.6 for the 3^rd^-to the last stimuli in the train) (Fig. 9B). For a cell, a firing probability of 0.8 means that it fired in response to the stimulus in 20 out of the 25 trials tested. Sensory adaptation is thus frequency dependent, as was already shown in experiments (Manuel A Castro-Alamancos, 2002; Martin-Cortecero and Nuñez, 2014). Not only VPL_TCs, but also Rt_RCs receive input from cortex and inhibit VPL_TCs cells when indirectly activated by sensory inputs, as shown previously (Fig. 8) and could potentially suppress thalamic activity.

We thus wondered why the net effect of cortical activation was enhancement (and not suppression) of sensory responses. We found that Rt activity was marked in response to the first two-three stimuli in the train and tended to decrease with successive stimuli (Fig. 9C), as a result of synaptic depression in the synapses between VPL and Rt neurons (Fig. 5). Moreover, we found that the topographical properties of the Rt-mediated inhibition were affected by CT activation, with an increase of the extent of surround inhibition in the VPL when the cortex was activated (see <u>Video 2</u>). This result shows how the cortex can shape thalamic receptive fields, via direct excitation of the thalamus and indirect inhibition via the Rt.

The model also reproduces the experimental results with single sensory stimuli (Fig. S2). We found that VPL_TCs respond to lemniscal activation with subthreshold EPSPs, single or multiple spikes (Fig. S2A). The increased depolarization, caused by the activation of CT afferents, results in a decrease in EPSP amplitude, suggesting a contribution from the low-threshold Ca^2+^ current at baseline (Fig. S2B-E) (Fig. 9C).

#### 3.5.3 Cortical enhancement of sensory responses is frequency dependent and frequency selective

Corticothalamic effects on thalamic activity are dynamic and depend on the mean firing rate and pattern of cortical activation (Crandall et al., 2015; Kirchgessner et al., 2020). We thus explored how the results above, showing net cortical enhancement of sensory responses, generalized to different frequencies of sensory and cortical activation.

Above, we saw that when sensory stimuli of elevated frequencies are presented (e.g. 10 - 20 Hz) thalamic responses are highly depressed with a greatly reduced probability (∼0.1) of firing (Fig. 9B). We then tested cortical activation at different mean firing rates and found that increased cortical firing corresponded to an increased efficacy of CT inputs in counterbalancing sensory adaptation, i.e. increase in firing probability (Fig. 9D). Surprisingly, this happens for sensory stimuli up to 10 Hz, while at 20 Hz the input from CT afferents is no longer sufficient to counterbalance the adaptation, and maximal firing probabilities were <0.5.

We found that cortical enhancement was frequency selective and maximal when both ML and CT afferents were activated at ∼10 Hz (Fig. 9D). The enhanced gain of thalamic input at 10 Hz ML inputs is due to Rt inhibition and the properties of subthreshold responses in TCs (Fig. 9E and Fig. S2F-G). Around 10 Hz, the stimuli are well-timed with the indirect inhibition coming from the Rt and activation of the low-threshold Ca^2+^ current in TC cells. For higher frequency of sensory inputs EPSPs decrease in amplitude, due to synaptic short-term depression and limited activation of the low-threshold Ca^2+^ current, and the cortical input is less effective in counterbalancing sensory adaptation (Fig. 9E). Furthermore, increased activation of the Rt also contributes to inhibiting VPL_TCs responses to 20 Hz ML stimuli.

### 3.6 Spindle-like oscillations emerge in the thalamoreticular microcircuit model

It is well established that sleep spindles are generated through Rt and TC reciprocal interactions (Steriade et al., 1985; von Krosigk et al., 1993), and numerous *in vivo*, *in vitro,* and *in silico* studies have explored the thalamoreticular mechanisms underlying the generation of spindle-like oscillations (T. Bal et al., 1995; Destexhe et al., 1993, 1996; Golomb et al., 1996; Kim et al., 1995; von Krosigk et al., 1993; Wang et al., 1995). Spindle generation entails the activation of Rt neurons which hyperpolarize TC cells via Rt-TC inhibitory connections (Sanchez-Vives and McCormick, 1997; Steriade et al., 1985). This hyperpolarization primes TC cells for rebound bursts (via intrinsic pacemaker I_h_ and low-threshold Ca^2+^ currents), which perpetuate the spindle cycle through a “ping-pong” interaction between the Rt and TC (Sohal et al., 2006). These cellular and synaptic interactions can all be influenced by CT afferent inputs and diverse neuromodulatory contributions (Fernandez and Luthi, 2019). Recent technical advances have enabled the study of spindle oscillations using optogenetic activation of the Rt *in vivo* (Halassa et al., 2011; Kim et al., 2012; Thankachan et al., 2019). We use these prior studies to validate that the model recapitulates known spindle generation mechanisms, without having been explicitly built to do so.

#### 3.6.1 Activating the reticular nucleus increases thalamic bursting and initiates spindle-like oscillations

The Rt nucleus has long been recognized as the spindle pacemaker for its ability to generate oscillations in thalamic networks, acting in concert with post-inhibitory thalamic bursts (T Bal et al., 1995; Jahnsen and Llinás, 1984; Steriade et al., 1985). We thus investigated whether the model can reproduce experimental findings, showing that optogenetic activation of Rt neurons results in increased thalamic bursting and spindle oscillations (Halassa et al., 2011).

We activated a subset of neighboring Rt_RC cells in the model with a 20 ms current pulse, to simulate a brief optogenetic activation of the reticular nucleus (Fig, 10).We activated 750 central Rt_RCs in the *in vivo*-like condition (see Methods). We found that activation of the Rt results in brief (∼250 ms) oscillatory responses in both Rt and VPL cells (Fig. 10A). Local field potential (LFP), calculated in the VPL (Fig. 10B) resemble the average membrane potential of VPL_TCs (Fig. 10A) and reveals oscillations at ∼10 Hz (Fig. 10C), consistent with the spindle frequency range (7-15 Hz) *in vivo* (Halassa et al., 2011). Single cell responses show increased bursting in Rt_RCs as well as VPL_TCs (Fig. 10D)

We found that Rt_RC cells respond to the stimulus with increased spiking activity, followed by a second peak ∼125 ms after the stimulus (Fig. 10E). The second peak is mainly generated by the post-inhibitory rebound responses in VPL_TCs. In line with the experimental findings, we found an increase in VPL_TCs burst in the 100 ms following the Rt stimulus (Fig. 10F). While the firing rate in VPL_TCs cells does not increase significantly after the stimulus, the firing mode changes, with increased burst probability after the stimulus (Fig. 10G).

#### 3.6.2 Simulated cortical UP and DOWN states initiate spindle-like oscillations

The correlation between cortical and thalamic oscillatory activity has been extensively studied *in vivo* in anesthetized animals and, more recently, in naturally sleeping rodents (Contreras and Steriade, 1996, 1995; Fernandez et al., 2018; Slézia et al., 2011; Steriade, 2006; Urbain et al., 2019). Spindles often occur during cortical UP states, when cortical neurons are more active (Destexhe et al., 2007; McCormick and Bal, 1997; Steriade et al., 1993). Synchronous Rt bursts are often initiated by inputs from layer 6 cortical neurons, which provide a major source of excitatory input to the Rt (Fuentealba et al., 2005; Fuentealba and Steriade, 2005). The simulations in the previous section confirmed that spindle-like oscillations can be initiated by directly activating the reticular nucleus. Here, we show that cortical UP states, that provide excitation to the Rt as well as the VPL, can generate thalamic spindle-like oscillations.

We simulated a transition from a wakefulness-like to NREM-like state characterized by alternating DOWN and UP states. We simulated the DOWN and UP states by periodically removing the firing background from the cortical afferents for 500 ms and reactivating it for another 500 ms to the same level as the one used for our standard *in vivo* wakefulness-like condition (Fig. 11 and <u>Video 3</u>).

Interestingly, during the DOWN state Rt_RCs are highly hyperpolarized (< -₋70 mV), while VPL_TCs are less affected (Fig. 11A). This result showed that, in the model, the major drive of activity in the Rt is coming from CT afferents. Moreover, although Rt activity is markedly reduced, thus removing inhibitory inputs to the VPL, the net effect of a cortical DOWN state is mild hyperpolarization of the VPL. This suggests the contribution of cortical activity, along with spontaneous lemniscal firing, in driving activity in the VPL. During the simulated cortical

UP state, Rt_RCs fire robustly, predominantly low-threshold bursts, causing deep IPSPs in the VPL_TCs, that in turn often respond with post-inhibitory rebound bursts, the hallmark of spindle-like activity in the thalamus (Fig. 11B).

Taken together, these results show that cortical input has a dramatic impact on thalamic and reticular activity, depending on its temporal structure: when it provides continuous drive, it promotes uncorrelated firing, such as the one we observed during wakefulness-like states. During periodic activity, causing an alternation of hyperpolarization and depolarization in the Rt, for example during DOWN and UP states, it promotes synchronous spindle-like oscillations. Prolonged hyperpolarization in the reticular nucleus has been shown to precede spindle sequences in anesthetized cats (Fuentealba et al., 2004).

#### 3.6.3 Spindle-like oscillations are maintained by “ping-pong” interactions between the reticular nucleus and thalamus

After verifying that the model was able to generate spindle-like oscillations after stimulation of the Rt and with simulated cortical UP and DOWN states, we dissected the mechanisms underlying their initiation, duration and termination, in the isolated thalamoreticular network (without firing activity from ML and CT afferents).

Without firing from the afferents, all the neurons are hyperpolarized (Fig. 12). We began by simulating the network at rest with spontaneous synaptic release as the only source of input and activating a set of neighboring Rt_RCs with a 20 ms pulse as before (see Fig. 10 and <u>Video</u> <u>4</u>). We confirmed that synchronized burst firing in Rt cells is a potent trigger of spindle oscillations. Synchronous bursts in Rt_RCs cells cause large amplitude IPSPs in VPL_TCs that last ∼100 ms, which have the optimal duration to remove the inactivation of the low-threshold Ca^2+^ current in VPL_TCs and activate the I_h_ current that, in turn, fire post-inhibitory rebound spikes and bursts. This result confirms the important role of these intrinsic neuronal mechanisms in the generation of spindle-like oscillations (Astori et al., 2011; Liu et al., 2011; Pellegrini et al., 2016; Talley et al., 1999, Bal et al., 1995; Jahnsen and Llinas, 1984). Furthermore, the negative reversal potential for Cl^-^ in the thalamus contributes to large amplitude GABA_A_-mediated IPSPs (Huguenard and Prince, 1994; Ulrich and Huguenard, 1997). All these intrinsic and synaptic properties were taken into account in our neuron and synapse models (see Methods).

We then varied the synapse release probability (P_rel_) in the connections between VPL_TCs and Rt_RCs and vice versa. The back-and-forth of activity lasts longer for *in vitro*-like P_rel_ and generates network oscillations at a frequency of ∼5-6 Hz, visible as increased peaks of activity in the average membrane potentials (Fig. 12 B2). The frequency is similar to some of the barrages of IPSPs in TC cells recorded during spindle waves in ferrets *in vitro* (T. Bal et al., 1995).

These results confirmed that the “ping-pong” of activity is necessary to generate spindle-like oscillations in thalamic networks for a wide range of synapse efficacy in the Rt to VPL connections. While the reliable excitation of the Rt from the thalamus has been investigated *in vitro* (Gentet and Ulrich, 2003), we further show that a minimal degree of synaptic efficacy of the Rt_RC to VPL_TC connections is necessary to initiate the oscillation.

#### 3.6.4 The duration of spindle-like oscillations is determined by the efficacy of reticulothalamic connections

We found that a number of synaptic mechanisms, or instance P_rel_ and short-term synaptic depression, have an effect on the duration of spindle-like oscillations. To the best of our knowledge, this is the first account of how synapse efficacy between the Rt and the thalamus can affect the duration of spindle-like oscillations.

The release probability of Rt to TC connections has a marked effect on the oscillation duration, which lasted less than 500 ms for P_rel_ corresponding to *in vivo*-like conditions and 700-800 ms for P_rel_ corresponding to the *in vitro*-like condition (Fig. 12B). The PSD peak, calculated from the firing rate histograms (see Methods), increased smoothly when increasing P_rel_ between Rt and VPL (Fig. 12A).

Short-term synaptic depression causes P_rel_ to decrease over time and its recovery is governed by the recovery time constant from synaptic depression (τ_RecDep_). When we varied τ_RecDep_ in both connections between Rt_RCs and VPL_TCs (and vice versa), we found again a predominant effect of the Rt_RC to VPL_TC connection, with long oscillations when τ_RecDep_ was small. Unlike the variation of the PSD peak with P_rel_, in the case of τ_RecDep_ the change was discontinuous, in particular when we varied the value in the Rt_RCs to VPL_TCs connections.

#### 3.6.5 Gap junctions increase the duration of spindle-like oscillations by propagating low-threshold bursts across the reticular network

Gap junctions (GJs) between reticular neurons can efficiently transmit low-threshold bursts between cells, promote spiking correlations when the coupling is strong and synchronize the activity in the reticular nucleus *in vitro* (Landisman et al., 2002; Long et al., 2004). GJs have been hypothesized to contribute to the maintenance of network oscillations, through network synchronization (Beenhakker and Huguenard, 2009; Fernandez and Luthi, 2019). Here we show a direct contribution of GJs to the duration of spindle-like oscillations, along with the spatial properties of the Rt recruitment.

Starting from the *in vitro*-like P_rel_ condition used previously (Fig. 12B2), we studied the effect of removing GJs on spindle-like oscillations (Fig. 13). We then activated a central subset of Rt_RC with a current pulse as before. When the GJs conductance was set to 0 the activation of a subset of Rt_RCs was not sufficient anymore to elicit spinde-like oscillations. When we compared the spiking activity of Rt_RCs we found that, when GJs are present, there is some extra activity after the stimulus (see arrow in Fig. 13A).

We then compared the spatio-temporal activities in the Rt and VPL in the two conditions (control and GJs removed, see <u>Video 5</u>). While the responses during the stimulation period are comparable, with only the central Rt_RCs being active, the spatio-temporal activity is different after 40-50 ms (Fig. 13B). When GJs are present, cells that are not directly stimulated are recruited and cause extra spiking activity in the Rt. When we analyzed the membrane potential along the lateral dimension of the microcircuit in the two conditions, we found that the excitation in the Rt spreads to more lateral neurons and lasts for a longer time, with corresponding increased inhibition in the VPL, thus increasing the probability of rebound responses in VPL_TCs (Fig. 13C-D). This lateral spread in the Rt propagates from central cells to peripheral ones with the latter being activated later after the stimulus (Fig. 13C,E). Slow signals, such as low-threshold spikes are transmitted efficiently via GJs, and their amplitude decreases in more peripheral cells (Fig. 13E). When GJs are present, the IPSPs from the Rt are visible in central as well as more peripheral cells in the VPL, indicating that the activity propagates broadly along the Rt axons.

We have shown previously that GJs connect neurons that are up to 250-300 μm apart (see Fig. 6). These simulations further highlight that GJs have functional effects along the lateral extent of the microcircuit. Low-threshold spikes are transmitted along the dendrites of Rt_RCs and contribute to the maintenance and duration of spindle-like oscillations.

#### 3.6.6 The termination of spindle-like oscillations is determined by short-term depression and buildup of intra-reticular inhibition

Different intra-thalamic cellular and synaptic mechanisms have been shown to play a role in spindle termination, including Ca^2+^-dependent upregulation of *I_h_* current in TC cells (Bal and McCormick, 1996; Lüthi and McCormick, 1998), progressive hyperpolarization of reticular neurons for the activation by Na^+^- and Ca^++^-dependent K^+^ current in ferret (Kim and McCormick, 1998). External inputs, such as desynchronized cortical activity and noradrenergic input from the locus coeruleus could play a role as well (Aston-Jones and Bloom, 1981; Bonjean et al., 2011).

In our model, we found additional cellular and synaptic mechanisms that are sufficient to cause the termination of spindle-like oscillations, that have not been directly investigated so far. All intrathalamic connections in the model are governed by depressing synapses (Fig. 5), which contributes to the decrease in efficacy between Rt and VPL neurons over time. By systematically scaling the time constant of recovery (from short-term synaptic depression) between 0 (no depression) and 1 (normal depression) we found that at 85% of the control value (299±39 ms) in the Rt_RC to VPL_TC connections, the oscillation no longer terminates (Fig. 12D-E). On the other hand, short-term depression in the VPL_TC to Rt_TC pathway has very limited effect on the termination and could be even removed (scale value of 0 in Fig. 12D), if it is still present between Rt_RCs and VPL_TCs.

We saw that the bursting activity of Rt_RCs is one of the key determinants in the initiation and duration of spindle-like oscillations. However, Rt_RCs cells inhibit each other through GABAergic synapses and this can contrast their activity. The importance of inhibition between reticular neurons has been already suggested in experimental work (Beenhakker and Huguenard, 2009; Makinson et al., 2017; Sohal and Huguenard, 2003), although some studies have questioned the existence of inhibitory Rt-Rt synapses (Hou et al., 2016). In the model, we were able to directly study the impact of Rt-Rt inhibition on network oscillation. We show that decreasing the conductance in these synapses results in non-terminating oscillations (Fig. 12F). When comparing the firing activity, we found that the percentage of Rt_RC cells participating in each oscillation cycle increases when mutual inhibition is reduced (Fig. 12F).

#### 3.6.7 Waxing and waning of spindle-like oscillations emerges due to intrinsic cellular and synaptic mechanisms

Waxing and waning are defining characteristics of spontaneous sleep spindles recorded at the cortical level in the EEG, local field potential and in thalamic recordings (T. Bal et al., 1995). Despite this characteristic pattern of activity, the mechanisms underlying the spindle-shaped oscillation, is not completely clear (Clawson et al., 2016) A line of research has proposed that cortical feedback is necessary in the first cycles of the oscillation (waxing), when the involvement of TC cells is still small. During the waxing, increasing activity of TC neurons would cause progressive activation of cortical neurons and subsequent stronger CT thalamic feedback and synchronous activity in the thalamus (Kandel and Buzsáki, 1997; Lüthi, 2014). During the waning phase, cortex would provide desynchronizing input to the thalamoreticular network (Bonjean et al., 2011; Timofeev et al., 2001).

Although the bidirectional interaction between thalamus and cortex can contribute to the waxing and waning of the oscillation, in our model we found that it can also be sustained by the thalamoreticular microcircuit alone (Fig. 12G left). Specifically, the waxing of a spindle oscillation is created by the rhythmic recruitment of additional neurons, first in the reticular nucleus, and second through the “ping-pong” interaction with the thalamus - each “ping” from the Rt successively recruits additional neurons and generate a stronger “pong”, via low-threshold bursts in VPL_TC cells. Gap junctions play a key role in this process through their ability to rapidly recruit Rt neurons. To generate the ping-pong, a minimum degree of inhibition from the Rt is necessary, as we have shown when we varied the release probability in the Rt_RC to VPL_TC connections (Fig. 12A), along with reticular low-threshold bursts. In the model, only a small fraction (<10%) of VPL_TCs is active in each cycle (Fig. 12F), however their excitatory impact on Rt_RC cells is sufficient to maintain the “ping-pong” interaction between the populations.

The waning of the spindle-like oscillation is a result of the progressive reduction in the probability of synaptic release (due to short term synaptic depression), the subsequent decrease in PSP amplitudes, and the consequential reduction in recruitment of neuron firing during the “ping-pong” interaction between Rt and VPL neurons. At the same time, mutual inhibition between Rt_RCs progressively builds up, as more Rt_RCs are recruited, and prevents the spread of the activity, acting as a self-limiting mechanism. We showed that when we compared the number of Rt_RCs participating in each cycle of the oscillation in control condition and when mutual inhibition was reduced (Fig. 12F). The greatest difference was at the 5^th^ cycle of the oscillation: while in both cases the firing tended to decrease compared to the previous cycle, the activity was enough to start a new cycle of waxing and prevent the termination of the oscillation, when mutual inhibition was reduced.

From the perspective of different connections, we found that the excitatory pathway between VPL_TCs and Rt_RCs increases the oscillation, while mutual inhibition in the Rt decreases it (Fig. 12G right). The inhibition from Rt_RCs to VPL_TCs has positive or negative effects on the oscillations depending on the firing mode and timing of Rt_RCs: while synchronous bursting generated IPSPs with the optimal amplitude and duration for rebound responses in VPL_TCs (see above), single tonic spikes would just inhibit VPL activity, especially if Rt activity is not synchronized between cells. GJs contribute to the waxing of the oscillations, in particular when only a subset of Rt_RCs is active (e.g. at the start of the oscillation, see also

<u>Video 5</u>). By contributing to the build up of inhibition inside the Rt, they also play an indirect role in the waning.

#### 3.6.8 Depolarizing the reticular nucleus decreases spindle-like oscillation duration

Spindle oscillations in naturally sleeping (i.e. non-anesthetized) rodents are more easily evoked when thalamic activity is mildly synchronized and thalamic neurons are less active (Barthó et al., 2014; Halassa et al., 2011). Their features, such as frequency and duration, evolve during NREM episodes (Barthó et al., 2014; Urbain et al., 2019). In the model we show that the duration can vary as a result of membrane potential dynamics in the thalamus and the reticular nucleus.

We explored how the membrane potential level in the Rt and VPL affects the duration, frequency and peak firing during spindle-like oscillations (Fig. 14), starting from the *in vitro*-like condition (as in Fig. 12-13). We studied network dynamics over a wide range of depolarization levels in the Rt and VPL, through noisy current injection into Rt_RCs and VPL_TCs (see Methods), as an approximation of neuromodulatory influences onto thalamic and reticular activity (McCormick, 1992). The baseline potentials explored, went from hyperpolarized (resting membrane potential of the *in vitro*-like condition), mildly depolarized to close to firing threshold. As in previous simulations, 750 Rt_RCs located at the center of the microcircuit were stimulated with a 20 ms current pulse, which resulted in different spiking responses depending on the depolarization levels.

We first studied how oscillation duration and frequency were affected when the Rt or VPL alone were depolarized. When Rt_RCs are depolarized from their resting potential the duration of evoked spindle-like oscillations decreases (Fig. 14A). This result is in agreement with experimental findings showing how initial network state influences spindle duration, through the activity of reticular neurons (Barthó et al., 2014). Depolarization of Rt_RCs decreases their firing in response to the stimulus, as a result of decreased burst occurrence or a smaller number of spikes per burst (Fig. 14B). An increase in membrane potential, as small as 2-3 mV, is sufficient to observe this effect. The result is reduced inhibition to the VPL and decreased firing probability of VPL_TCs. Furthermore, the waxing-and-waning in firing responses tends to become a predominantly waning response when the Rt is depolarized (Fig. 14A).

#### 3.6.9 Depolarizing the thalamus increases spindle-like oscillation duration and frequency

When the membrane potential in the Rt is held constant (around −76 mV) and the VPL is depolarized, the oscillation increases in duration and frequency (Fig. 14C). VPL depolarization results in deeper and faster IPSPs (Fig. 14D). Deeper IPSPs result in stronger post-inhibitory rebound responses in VPL_TCs, which in turn excite more Rt_RCs causing a longer period of “ping-pong” interactions between the two populations, increasing the oscillation duration. Faster repolarization after the IPSPs is associated with post-inhibitory rebound responses occurring at shorter intervals, driving the “ping-pong” of activity at higher frequencies.

#### 3.6.10 Differential depolarization of the reticular nucleus and thalamus modulates spindle properties

For each combination of membrane potentials in Rt_RCs and VPL_TCs, we calculated oscillation duration, frequency and peak firing (Fig. 14E). The oscillation duration map shows that clear spindle-like oscillations (duration >= 500 ms) can only be evoked in a region, where Rt_RC membrane potentials are below −75 mV and VPL_TCs are more depolarized than Rt_RCs. If we assume that VPL_TCs neurons are in general more depolarized during wakefulness than during sleep, and that Rt_RC neurons are more hyperpolarized, this result suggests that spindle-like oscillations are easier to evoke at the transition between wakefulness-like to sleep-like states. When both VPL_TCs and Rt_RCs are at their baseline membrane potential, the frequency decreases to 5-6 Hz.

Modulation of spindle frequency and duration in the somatosensory cortex has been shown to vary during non-REM sleep in naturally sleeping mice; more specifically, the frequency decreases during NREM sleep along with TC neurons membrane potentials (Urbain et al., 2019) as shown in our simulations (Fig. 14D-E). Our results suggest that this modulation of frequency is already present at the thalamic level and can be transmitted to the cortex via thalamocortical projections. We also found that the Rt depolarization has a negligible effect on oscillation frequency (Fig. 14E).

#### 3.6.11 Simulating the effects of neuromodulation on thalamic and reticular neurons causes spindle-like oscillations to cease

Sleep spindles are a defining characteristic of stage 2 non-REM sleep, and appear less frequently in deeper stages of NREM sleep (Purcell et al, 2017, Cox et al., 2017; Fernandez and Luthi, 2019). The change in the incidence of spindles are due, at least in part, to neuromodulatory changes (Destexhe et al., 1994a; Osorio-Forero et al., 2021; Vyazovskiy et al., 2004). Our previous simulations, in the *in vitro*-like condition, showed that membrane potentials have a strong effect on oscillation duration and frequency (Fig. 14). They also indicated that when the VPL is hyperpolarized and the Rt is depolarized, spindles are less easy to evoke. We hypothesized that such differential polarization levels of the Rt and VPL would resemble the transition to deeper stages of NREM sleep, and result in decreased incidence of spindles.

We tested this hypothesis with the model in the *in vivo*-like condition using simulated cortical UP and DOWN states (as in Fig. 11). We then progressively depolarized the Rt and hyperpolarized the VPL, approximating the differential effect of neuromodulators on thalamic and reticular populations (Beierlein, 2014; Boucetta et al., 2014; McCormick and Prince, 1987). These simulated neuromodulatory changes cause the spindle-like oscillations to occur with reduced amplitude then to cease, while the cortically-generated up-down states continue.

## 4 Discussion

We developed the first morphologically and biophysically detailed model of a thalamoreticular microcircuit, and validated its behavior across different simulated conditions, including wakefulness and sleep. It was constructed and validated using experimental data by extending a workflow previously used to model cortical microcircuitry (Markram et al., 2015). The microcircuit model integrates experimental measurements of the detailed anatomy and physiology of single neurons, the three-dimensional organization of the reticular and VPL nuclei, neuron densities, synaptic anatomy and physiology, and electrical connectivity mediated by gap-junctions. The model gives rise to network-level phenomena observed in *in vivo* studies, including sensory adaptation, cortical enhancement, and rhythm generation (spindle-like oscillations), although the model was only fit to cellular and synaptic data. We found key roles for short-term synaptic plasticity, mutual inhibition within the reticular nucleus in the termination of spindle-like oscillations. We demonstrated that gap junctions, corticothalamic feedback, and membrane potentials play a key role in modulating the duration and frequency of spindle-like oscillations in the model. This model provides a comprehensive account of thalamoreticular network dynamics across different states, providing a tool to interpret alterations in corticothalamic feedback and rhythm generation in the thalamoreticular microcircuit at cellular and synaptic levels in health and disease. The full circuit model and its accompanying data are openly available (See Data and Model Availability below) to facilitate integration of new data and accelerate future studies.

The model differs from previous ones in several aspects, including scale (in terms of number of neurons), the level of biological detail and scope (Bazhenov et al., 2000; Bonjean et al., 2011; Brown et al., 2020; Bús et al., 2018; A. Destexhe et al., 1998; Destexhe et al., 1996, 1994b; Golomb et al., 1996). Prior models have largely used single compartment neurons, while this model uses reconstructed 3-dimensional neuron morphologies to constrain the biophysical models. The neuron morphologies provide further constraints on this model, highly constraining synaptic connectivity, rather than assigning an average connection probability. This model is additionally constrained by 3-dimensional estimates of Rt and TC cell density and the relative proportions of cell types in the context of an anatomical atlas, whereas prior models typically used 1-dimensional or 2-dimensional arrangements of cells and connectivity.

This model should be considered a first-draft reconstruction of thalamoreticular microcircuitry, and is made openly available to facilitate future refinements. Different thalamic nuclei have different mixtures of excitatory cells that have unique project characteristics to the cortex (Clascá et al., 2012; Jones, 2009). This model is representative of a primary somatosensory microcircuit and, therefore, includes only the “core” cells characteristic of sensory thalamic nuclei that preferentially target cortical layers III and IV. Higher-order thalamic nuclei contain additional excitatory cell populations that send “matrix” projections that preferentially target supragranular layers (I-III). As such, the present model represents a thalamic nucleus dominated by neurons with “core”, rather than “matrix” projections to the cortex. Future refinements could integrate additional such excitatory populations and their properties.

The cellular e-types in the present model do not capture the full diversity of known ion channels implicated in bursting behavior in reticular neurons or dendritic properties of thalamic interneurons. Inclusion of additional and more specific ion channel mechanisms, (e.g. variants of low-threshold Ca^2+^ Cav3.1, Cav3.2, and Cav3.3) in TC and Rt neurons could more accurately reproduce differences in bursting behavior in thalamic and reticular neurons, as well as dendritic properties of thalamic interneurons (Acuna-Goycolea et al., 2008; Astori et al., 2011; Huguenard and Prince, 1992; Lüthi and McCormick, 1998; Pellegrini et al., 2016).

Further refining the definition of morphological types and modeling the distribution of different me-types within both thalamic and reticular domains, could constrain further our derived connectivity (Deleuze and Huguenard, 2006; Krahe et al., 2011; Martinez-Garcia et al., 2020; Spreafico et al., 1991). Also the heterogeneity and laminar structure of the Rt can be taken into account in future refinements (Li et al., 2020; Martinez-Garcia et al., 2020). With such refinement, additional validation of the total neurite densities and synapses densities in the model could be used to further validate the connectivity (Kubota et al., 2018; Yin et al., 2020).

We found that many properties of intra-thalamic connectivity can be predicted by the axodendritic and dendrodendritic overlap of neuron morphologies, such as the distribution of synapse locations and the number of synapses per connections established by thalamic interneurons, as well as the convergence of synapses from thalamocortical axon collaterals, other reticular neurons, and corticothalamic afferents onto reticular neurons. An electron microscope study showed that most of the connections established by thalamic interneurons consist of single synaptic contacts (Morgan and Lichtman, 2020). The model showed the same characteristic, while accurately recreating the full distribution of single and multi synaptic connections accounting for different fractions of the connections depending on the specific pathway (e.g. 70% of connections between Rt and VPL neurons had more than two contacts, while for the other pathways, single synapse contacts were the majority).

Interestingly, the model recapitulates distance-dependent connectivity between Rt neurons as observed in dye-coupling experiments (Lee et al., 2014). We found that dendrodendritic appositions provide a sufficient basis for determining gap-junction locations and that the distance-dependent distribution of gap junctions between reticular neurons was determined by the extent of their dendrodendritic overlap.

We explored network dynamics in the model starting from spontaneous and sensory-evoked activity in wakefulness-like states. Our model makes it possible to simultaneously record direct and indirect sensory responses in the thalamus as well as in the reticular nucleus. Responses to stimuli were clearly visible in the reticular nucleus, as shown experimentally with sensory, auditory or visual stimuli (Hartings et al., 2000; Kimura, 2017; McAlonan et al., 2006). We found that robust inhibition from the Rt generates strong surround inhibition in the VPL. We found that cortical activation sharpens the spatial properties of sensory inputs, by evoking stronger Rt-mediated surround inhibitions, as recently shown in the visual thalamus (Born et al., 2021). This can contribute to the precise timing of VPL sensory responses, which were suppressed for ∼100 ms following the stimulus and in shaping the spatial properties of thalamic receptive fields, with the inhibition of the surrounding neurons.

As previously reported in anesthetized animals, the responses to trains of sensory stimuli were adapting for relatively low frequencies of 4-5 Hz, and the degree of adaptation increased with the increase of stimulus frequency (Diamond et al., 1992; Simons and Carvell, 1989; Castro-Alamancos 2002). The adaptation was reduced by simulated cortical activation, approximating cortical activity during activated states, allowing the transmission of high frequency inputs (Mease et al., 2014).

We found that the thalamoreticular microcircuit exhibits frequency-selective cortical enhancement. When simulating different frequencies of input and corticothalamic activation, we found peak enhancement of thalamic responses occurs for lemniscal inputs around ∼10 Hz. This enhancement occurs because cortical activation recruits sufficient Rt inhibition to activate *I_h_* and low-threshold Ca^2+^ currents, resulting in enhanced gain of input to TC cells at ∼10 Hz. This is striking, because rhythmic activity around this frequency emerges in thalamocortical networks during different behavioral states, such as sleep spindles and alpha oscillations during attention (Chen et al., 2016). Our simulations suggest that ∼10 Hz rhythms are intrinsic to thalamoreticular networks, and could be responsible for enhanced gain of sensory inputs and thalamocortical activity around that frequency. These results are consistent with the observation of overlapping thalamic mechanisms between sensory processing, attention, and sleep (Chen et al., 2016). The model, therefore, provides a self-consistent account of common cellular and synaptic mechanisms underlying thalamic gain and spindle generation across states.

This model gives rise to evoked spindle-like oscillations, without being explicitly built for this purpose. Although we did not optimize the model to reproduce spindle-like oscillations, they do emerge in the model and are robustly generated for parameters within experimentally plausible ranges. Furthermore, although the model was mainly based on *in vitro* findings, many aspects closely resemble thalamic activities during spindle oscillations *in vivo* in rodents (Barthó et al., 2014; Rovó et al., 2014; Urbain et al., 2019).

Consistent with previous experimental and modeling studies, spindle oscillations in the model are generated through a combination of intrinsic mechanisms, namely low-threshold calcium bursting in reticular neurons (Astori et al., 2011; Pellegrini et al., 2016) and the synaptic interactions between reticular and thalamocortical neurons (Destexhe et al., 1993, 1996; Li et al., 2017).

The model also provides novel insights into the mechanisms underlying the generation of sleep spindles in thalamic networks, including the role of TC-Rt synapse efficacy, short-term synaptic depression, mutual inhibition between Rt neurons and gap junctions in the thalamic generation of waxing-and-waning spindle-like oscillations..

Dendrodendritic gap junctions between reticular neurons have been known to transmit low-threshold bursts between cells, promote spiking correlations and synchronized activity (Landisman et al., 2002; Long et al., 2004). The model shows that gap junctions also influence spindle duration. Furthermore, the spatial organization of Rt dendrites and their connections through gap junctions also shows a clear functional role in propagating the stimuli along the horizontal dimension of the microcircuit and enhancing spindle-like oscillations.

The model accounted for spindle termination through synaptic mechanisms alone, and therefore did not require specific ionic mechanisms, such as Ca^2+^-dependent upregulation of the *I_h_*current in TC cells or desynchronizing cortical input (Bonjean et al., 2011; Bús et al., 2018; Destexhe et al., 1996, 1998). We show that the modeled cells and circuit have mechanisms that limit the duration of the oscillation, including short-term synaptic depression and mutual inhibition between reticular neurons. While the role of mutual inhibition has been already hypothesized based on experimental findings (Beenhakker and Huguenard, 2009; Fogerson and Huguenard, 2016; Kim et al., 1995; Makinson et al., 2017; Sohal and Huguenard, 2003), short-term synaptic depression has not previously been proposed (to the best of our knowledge) as a mechanism that contributes to spindle termination. Based on the role of Rt-Rt mutual inhibition, which decreases spindle duration and GJs, that tend to prolong it, we propose that contribution of Rt cells to spindle oscillations could have self-limiting factor: on the one hand, through GJs coupling, they promote synchronized IPSPs and post-inhibitory excitatory responses from the TCs, thus recruiting more and more Rt neurons; on the other hand, when a critical recruitment in the Rt is reached, the overall excitation is overcome by reciprocal inhibition between Rt neurons, short-term synaptic depression, and the oscillatory activity would limit itself.

The gradual increase in amplitude (waxing) and the gradual decline (waning) of spindle oscillations has been hypothesized to be due to changes in cortical activity impinging on the reticular nucleus and thalamus (Bonjean et al., 2011; Kandel and Buzsáki, 1997; Lüthi, 2014; Timofeev et al., 2001). However, the model demonstrates that cellular and synaptic mechanisms of thalamoreticular circuitry that underlie the increased recruitment of additional neurons during each cycle of a spindle can produce the waxing phenomena, while synaptic depression and the buildup of inhibition within the Rt are sufficient to explain the waning phenomena.

We also found that the isolated reticular nucleus was able to sustain spindle-like oscillations (not shown), if it was depolarized, simulating the presence of neuromodulators or if the inhibitory synapses between Rt_RCs were stronger, as suggested in previous models (Destexhe et al., 1994a). We also found that increased gap junction coupling can contribute to the generation of spindle-like oscillations in the isolated Rt (not shown). It may be that strong inhibitory interactions and gap-junctions coupling are prevalent in the specific foci of Rt (Fuentealba and Steriade, 2005), where spindles can be recorded when this nucleus is isolated.

We found that the depolarization level of thalamic and reticular neurons had a direct effect on oscillation frequency, duration, strength, and incidence of spindle-like oscillations. In the intact brain, different mechanisms contribute to modulate the membrane potential of thalamic neurons, such as neuromodulators or corticothalamic feedback. Although our depolarization paradigm is an approximation of the dynamic change of thalamic network states at the transition between wakefulness and sleep, this result builds upon known mechanisms of action of neuromodulators, such as acetylcholine (ACh). ACh neurons projecting to the thalamus and reticular neurons are particularly active during wakefulness, arousal and REM sleep (Boucetta et al., 2014), while NREM sleep is associated with a decrease in ACh release in the thalamus and Rt. Interestingly, ACh has opposite effects on thalamocortical neurons, where it is depolarizing, and in reticular neurons, where it causes hyperpolarization (Beierlein, 2014; McCormick and Prince, 1987). The decrease in ACh during transition from wakefulness to NREM sleep would be thus associated with hyperpolarization of the Rt and depolarization of the thalamus, as shown in our simulations. Noradrenaline (NA), has also been shown to depolarize thalamic and some reticular neurons *in vitro* (Lee and McCormick, 1996). Recently, NA levels have been found to fluctuate during NREM sleep *in vivo*, resulting in depolarized membrane potentials that impact the low-threshold bursting activity that underlies sleep spindles (Osorio-Forero et al., 2021). This NA-mediated depolarization reduces the incidence of spindles (Osorio-Forero et al., 2021). Other alterations in thalamic and reticular excitability could be caused by increases or decreases in the rate of synaptic activity impinging on thalamic and reticular cells, as well as the plasticity of corticothalamic and corticoreticular projections.

These findings have important implications when considering alterations of sleep spindles frequency, density and amplitude as a potential biomarker of disease. The model could serve as a tool to assess causal mechanisms affecting spindle oscillations and their properties during sleep in schizophrenia (Castelnovo et al., 2018; Ferrarelli et al., 2010, 2007; Manoach et al., 2016, 2014), neurodevelopmental disorders (Gruber and Wise, 2016), attention deficit hyperactivity disorder (Saito et al., 2019), Alzheimer’s disease (Weng et al., 2020), among others. For instance, spindle densities have been found to decrease in patients with Parkinson’s disease, while their density and duration are known to be sensitive to Alzheimer’s disease, along with other parameters quantifying their time-varying microstructure (Christensen et al., 2014; Kam et al., 2019; Ktonas et al., 2009; Weng et al., 2020). Furthermore, alterations to fast (12-15 Hz) spindles appeared to be more common in Alzheimer’s disease compared to slow spindles (9-12 Hz) (Weng et al., 2020). Previous studies have suggested that cortical mechanisms are required for the generation of fast and slow spindles (Timofeev and Chauvette, 2013), however the model shows that the thalamus itself can modulate the frequency of spindle-like oscillations, suggesting that thalamic mechanisms may be sufficient to generate the two classes of spindles.

In summary, we have developed a first-draft large-scale model of thalamic and reticular microcircuitry. Although it is, to the best of our knowledge, the most detailed model of its kind created so far, it is only a first step. Future studies will further refine the thalamoreticular model to take into account newly observed cellular and synaptic properties (Li et al., 2020; Martinez-Garcia et al., 2020), and explore thalamoreticular contributions to other functions, such as attention and the generation of the alpha rhythm (Ahrens et al., 2015; Chen et al., 2016; Gu et al., 2021; Makinson and Huguenard, 2015; Nestvogel and McCormick, 2021), when considered in the broader context of the thalamocortical loop.

## Data and model availability

The experimental data, ion channel models, single neuron models, synapse models, and circuit model are all available under an open access license. The portal includes data and model entities, including single neuron models, circuit files in SONATA format (Dai et al., 2020) and simulation output. The Thalamoreticular Microcircuit Portal is accessible at: (https://identifiers.org/bbkg:thalamus/studios/e9ceee28-b2c2-4c4d-bff9-d16f43c3eb0f).

## Supplementary figures

**Figure S1, related to Figure 2.**
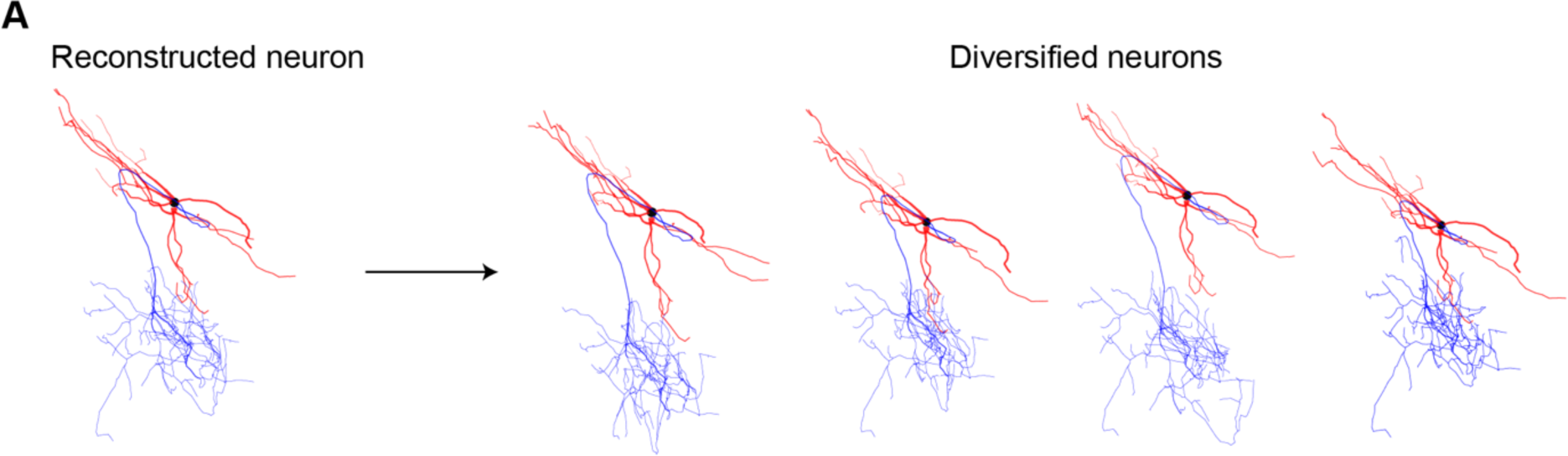
Morphology diversification. **(A)** The reconstructed neuron on the left was diversified to generate a sample of unique morphologies, by introducing variability (jittering in f branch lengths of 0 ± 20% and jittering in branch rotations of 0° ± 20°, mean ± standard deviation, see Methods) to the branch lengths and angles (see Methods.)

**Figure S2.**
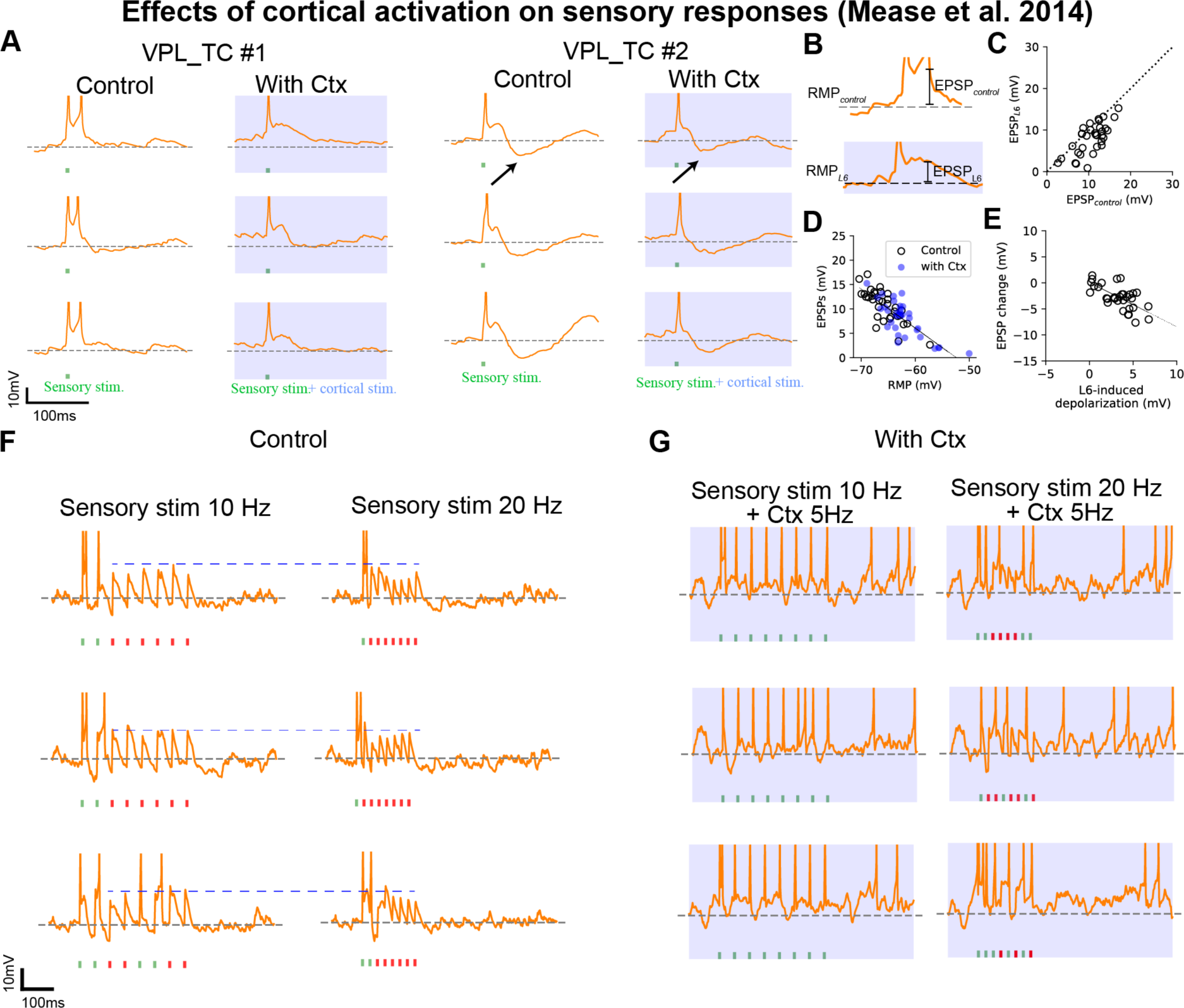
Cortical activation decreases sensory adaptation by depolarizing VPL_TCs and enhances responses to stimuli at ∼10Hz preferentially. **(A)** Left: Single cell recording of a VPL_TC neuron (3 of 25 repetitions are shown) that responded to the sensory stimulus (green) with a burst of two spikes. The sensory stimulus was generated with brief synchronous activation of 160 ML afferent fibers. Right: activation of the CT afferent fibers (blue) stimulus depolarized the cells and shifted their responses to single spikes. VPL_TC #2 showed a marked IPSP (arrow) following the stimulus-evoked spike, which was reduced with cortical activation. CT fibers were activated with noisy input at 4 Hz, 200 ms before the sensory stimulus to approximate the optogenetic protocol in Mease et al., 2014. (**B**) Illustration of different metrics used to quantify subthreshold responses (in a time window of 50 ms after the stimulus to the sensory stimuli (cfr. Mease et al, 2014). (**C**) Population analysis of VPL_TC cells (n=50, values are median of the 25 repetitions for each cell) showing the decrease of EPSP amplitude with cortical activation (EPSP_L6_). This effect is due to partial inactivation of the low-threshold Ca^2+^ conductance, but inhibition from the Rt can’t be excluded. (**D**) The amplitude of the EPSPs (both with and without cortical activation) is negatively correlated with the resting potential of the cell (r=−0.8). This is due to a greater availability of ionic currents activated at hyperpolarized potentials and greater driving force of excitatory conductances (whose reversal potential is 0 mV). (**E**) Correlation between the magnitude of sensory response change (EPSP_L6_ − EPSP with sensory stimulus only) and the depolarization induced by the cortical activation. The line shows the best fit (r=−0.8). This shows that greater cortical activation corresponds to decreased responses to sensory input (for single stimulation). (**F**) Single cell recordings of a VPL_TC neuron (3 of 25 repetitions are shown) with sensory stimulus at 10 Hz (left) and 20 Hz (right). Note the smaller amplitudes of the EPSPs in response to the 20 Hz stimulus. (**G**) Same cell as in F, with activation of cortex (noisy input at 5 Hz). Note a higher number of spiking failures (red ticks) with 20 Hz sensory stimulation. With 10 Hz stimulus, cortical activation made the cell fire in response to each pulse of the stimulus (green ticks).

## Movies

<u>Video 1</u>: Evoked sensory activity, *in vivo*-like condition (related to Fig. 8):

Simulation of evoked sensory activity with brief activation of 160 lemniscal fibers located at the center of the microcircuit. The microcircuit is displayed from the VPL side.

<u>Video 2</u>: Sensory adaptation, control vs. cortical input, *in vivo*-like condition (related to Fig. 9):

Simulation of sensory adaptation and the enhancement of sensory responses by cortical activation. The microcircuit is displayed from the VPL side. The stimulus consists of brief activation of 160 lemniscal fibers, repeated 8 times at 8 Hz (left). On the right, the same stimulus is delivered during cortical activation (4 Hz of noise stimulus from the corticothalamic fibers).

<u>Video 3</u>: Transition from wakefulness-like states to simulated cortical UP and DOWN activity, with spindle-like oscillations appearing during the UP state (related to Fig. 11) At the start of the simulation the network is in an wakefulness-like state, with spontaneous activity generated by spiking from lemniscal and corticothalamic fibers. The microcircuit is displayed from the front side, the upper part corresponds to the Rt and the lower one to the VPL. A cortical DOWN phase is simulated by interrupting the spiking from the corticothalamic fibers, while it is reactivated during the UP phase. During the UP state, spindle-like oscillations emerge in the microcircuit as activity “ping-pong” between the Rt and the VPL.

<u>Video 4</u>: Spindle-like oscillations, *in vitro*-like condition (related to Fig. 12)

Simulation of the *in vitro*-like condition, without activity from the lemniscal and corticothalamic fibers. The microcircuit is displayed from the front side, the upper part corresponds to the Rt and the lower one to the VPL. A brief pulse of current is delivered to the center of the Rt and generates the ping-pong of activity between the Rt and VPL.

**<u>Video 5</u>: Spindle-like oscillations, control vs. gap-junctions removed (related to** **Fig. 13**): Simulation in the *in vitro*-like condition. The microcircuit is displayed from the Rt side and Rt_RC dendrites are shown (5% of their density). A pulse of current is delivered to the center of the Rt in the control condition (left, with gap-junctions present) and with gap junctions removed (right).

## Acknowledgments

We thank Karin Holm from the Blue Brain Project for her assistance with manuscript editing and submission and our experimental collaborators: Mario Rubio and César Porrero (Autonomous University of Madrid), and Yun Wang and the team at Wenzhou Medical University for their contributions with morphology reconstructions.

